# Modulation of Avian Iridescence via Melanogenesis

**DOI:** 10.64898/2026.01.26.701322

**Authors:** Soraia Barbosa, Carina Bittner, Roberto Arbore, Pedro M. Araújo, Paulo Pereira, Pedro Andrade, Csaba Fekete, Rita Afonso, Sandra Afonso, M. Francisca Amorim, Cristiana I. Marques, Michaël P. J. Nicolaï, João Carita, Jindřich Brejcha, Ricardo J. Lopes, Joel M. Alves, Fernando Cruz, Jèssica Gómez-Garrido, Cristina Zamarreño, Marta Gut, Tyler Alioto, Muhammad Akhtar Ali, Andrea Hilpert, Anita Hecht, Erdmann Spiecker, Benjamin Apeleo Zubiri, Shosuke Ito, Kazumasa Wakamatsu, Leif Andersson, Joseph C. Corbo, Nicolas Vogel, Miguel Carneiro

**Author notes:** Correspondence (M.C.), (N.V.); (S.B.). Co-first authors (Soraia Barbosa, Carina Bittner, Roberto Arbore, Pedro M. Araújo). Co-senior authors (Nicolas Vogel, Miguel Carneiro).

## Abstract

The iridescent colors of birds originate in nanoscale feather structures that interact with light. Although the physical principles governing avian iridescence are well-established, the molecular mechanisms assembling these nanostructures into photonic materials remain unknown. Here, we integrated genomic, cellular, and optical approaches to investigate the molecular basis of iridescent color formation in birds. We leveraged mutations that arose in the domesticated Indian peafowl and natural color variation in wild birds. In peafowl, we identified eight genes underlying gains, losses, and shifts of iridescence, whereas in two wild species, we used single-cell transcriptomics to profile asymmetric feathers in which iridescent and non-iridescent barbules develop on opposite sides of the same feather. We found that all peafowl mutations mapped to melanogenesis genes, showing that variation in this pathway can modify diverse geometric features of feather photonic nanostructures. Mutations altering melanin composition collapsed multilayered photonic systems and yielded non-iridescent tissues, whereas changes in melanosome abundance, elongation, or deposition timing generated multilayer architectures with variable periodicity and color. We further show in peafowl that transitions from non-iridescent to iridescent plumage were associated with single-nucleotide mutations in a melanogenesis gene, suggesting that, in species already capable of forming multilayered nanostructures, iridescence can be gained by relatively small genetic changes. Single-cell transcriptomes from wild species also supported extensive melanocyte-centered regulatory rewiring associated with iridescence, including changes in melanosome maturation, trafficking, and intercellular signaling. Together, these results show that the nanoscale order underlying iridescence is developmentally plastic and highly responsive to the melanogenic environment.

**Significance Statement:** Bird feathers can produce brilliant, shifting colors, but how these colors form during development has remained poorly understood. By comparing the genomes of color variants of domesticated peafowl and transcriptomes during feather development in wild birds, and combining these with chemical and optical analyses, we show that mutations in genes associated with melanogenesis can profoundly alter the amount, shape, and arrangement of the melanin granules inside feathers that interact with light and produce iridescence. These changes can switch feather appearance between dull and iridescent, or generate entirely new colors by restructuring how those granules are layered at the microscopic scale. Our results reveal a direct and flexible link between pigment production and the physical structures that generate iridescence in birds.

## Introduction

From butterfly wings to bird feathers, animal coloration arises through two primary mechanisms: pigmentary and structural coloration (1–4). In birds, pigments generate hues ranging from blacks and browns to bright reds and yellows (5–9), and their genetic and biochemical underpinnings are increasingly well understood (*6–9*). However, some of the most visually striking feather effects—glossy blacks, bright blues, and iridescence—result from structural coloration (10–12). This form of coloration originates from the interaction of light with highly organized nanoscale features within feathers (10, 13), and evolves more rapidly than pigment-based colors (14–19). Notable examples include the elaborate plumage of hummingbirds and peacocks, whose eyespot feathers have become textbook examples of sexual selection. Structural colors are crucial to avian biology, supporting functions such as communication and camouflage (10, 20, 21), and serve as models for bioinspired technologies (22–24).

Iridescence in bird feathers is a form of structural coloration that produces brilliant colors that vary with the angle of observation or illumination (10, 25). Avian iridescence arises from multilayer nanostructures consisting of keratin and melanosome layers within feather barbules. Melanosomes are intracellular organelles produced by pigment cells (melanocytes) that synthesize and store melanin, and can vary in shape and internal structure, including rod- or platelet-shaped melanosomes, which can be either solid or hollow (10, 25–28). For example, many hummingbirds possess hollow melanosomes, sometimes with a platelet-like morphology, whereas platelet-shaped melanosomes are also found in some starlings and birds-of-paradise. Here, we focus on the most widespread mechanism of avian iridescence, exemplified in peacock feathers, where high-refractive-index rod-shaped and solid melanosomes are embedded within a lower-refractive-index matrix of keratin and air (29–31). During feather growth, these components become organized into highly ordered multilayered nanostructures that reflect light through constructive interference (when reflected light waves reinforce one another), producing iridescent colors (4, 10, 25, 32). Variation in melanosome morphology and arrangement, as well as barbule geometry, accounts for much of the nanostructural and optical diversity of iridescent plumage (15, 25, 27, 29, 30, 33). Despite decades of optical and anatomical work clarifying the physical basis of avian iridescence, the developmental and molecular mechanisms that generate these nanostructures remain essentially unknown (2, 11). Although physicochemical self-assembly is thought to play an important role in their formation (26), the extent to which their assembly is also shaped by direct cellular patterning or other developmental processes remains unclear.

In this study, we investigated the molecular basis of avian iridescence using complementary systems and approaches. First, we studied eight color variants of the domesticated Indian peafowl (*Pavo cristatus*), in which Mendelian mutations cause losses and shifts in iridescence across the whole body, including the eyespot (**Fig. 1**; **fig. S1**). These color varieties, each a single mutational step from wild-type, offer a powerful system to link genotype, nanostructure, and optical phenotype. By combining genetic mapping, pigment chemistry, ultrastructural analyses, and optical measurements, we linked these mutations to changes in feather photonic nanostructures. We then examined the Black-shoulder mutation, another domesticated variant that confers a localized gain of iridescent coloration in males, which shows that minimal genetic changes in melanogenic genes can generate new photonic structures with iridescent color. Finally, to determine whether similar mechanisms operate in nature, we profiled asymmetric feathers from wild birds using single-cell transcriptomics, comparing iridescent and non-iridescent barbules developing within the same feather. Complementing the mutational insights provided by the peafowl system, these wild-bird analyses reveal how iridescent coloration is generated through the integration of multiple developmental processes, including melanogenesis, keratinization, and intercellular signaling. Our study provides molecular insights into the generation of iridescent colors of bird feathers, a phenomenon that is better understood from physical and anatomical perspectives.

**Figure 1.**
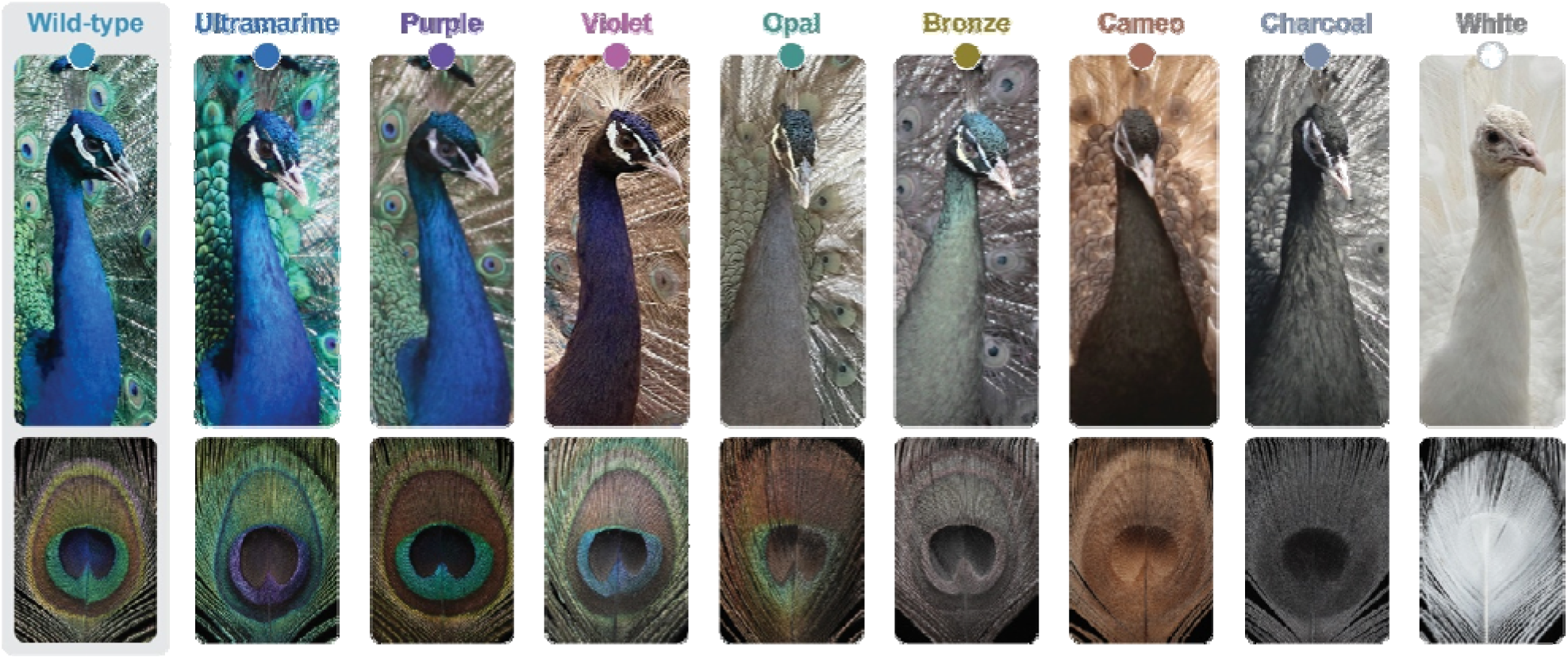
Mendelian color mutants in the Indian peafowl (*Pavo cristatus*). This figure illustrates the striking phenotypic effects of Mendelian color mutations on the overall appearance and eyespot coloration in peacocks. From left to right: wild-type, Ultramarine, Purple, Violet, Opal, Bronze, Cameo, Charcoal, and White. Additional photographs showing the full-body plumage phenotype of each color variety are provided in **fig. S1**.

## Results

### Peafowl color variation maps to melanogenesis genes

Our current understanding of the anatomical organization of photonic structures in avian feathers provides a framework to hypothesize mechanisms through which structural coloration might evolve in birds. Modifications in melanin biosynthesis, keratin production, feather morphogenesis, or spatial signaling pathways that coordinate intracellular organization are all potential mechanisms that might affect the resultant plumage color (11, 34). However, progress in testing these hypotheses has been limited because iridescent traits rarely segregate within species, restricting the discovery of functionally relevant genetic variants (35, 36). Domesticated color mutants of the Indian peafowl offer a rare opportunity to directly interrogate the molecular determinants of structural coloration.

To investigate the molecular basis of color variation in peafowl, we began by generating a *de novo* reference genome (**table S1; figs. S2 and S3**). Using long-read Nanopore sequencing, 10x Genomics linked-reads, and Hi-C chromatin conformation capture data, we assembled a chromosome-scale genome (1.04 Gb, N50 = 93.07 Mb) with 96.2% completeness of single-copy avian orthologs. Gene annotation, assisted by transcriptome data from five tissues, identified 21,618 protein-coding genes. We further resequenced the genomes of 368 birds representing wild-type, color mutants known to be governed by single Mendelian loci, and color varieties likely arising from combinations of two or more mutations (mean depth = 10.1× ± 5.3 SD). For a subset of individuals representing wild-type and several mutant varieties (*n* = 27), we profiled gene expression in regenerating feather follicles using RNA-sequencing.

Genome-wide association studies (GWAS) recovered significant loci for eight Mendelian peafowl varieties (**Fig. 2A**), thus genetically characterizing the domesticated variants analyzed below for their ultrastructural, cellular, biochemical, and optical phenotypes. For seven varieties, the associated loci contained a single compelling candidate gene: White (*EDNRB2*), Cameo (*AP3S1*), Bronze (*HPS3*), Opal (*MLPH*), Charcoal (*LYST*), and Purple and Violet (*TYRP1*). For Ultramarine, several genomic regions showed association with the phenotype, likely reflecting the lower number of sampled individuals for this variety (*n =* 11) together with extensive intra-flock breeding practices following the recent origin of the mutation, which produces stronger population structure. Among the top candidates, *MITF* represents the strongest based on its well-established role in pigmentation and the autosomal inheritance of the trait. The candidate genes across phenotypes span diverse steps of the melanogenic pathway, including melanoblast differentiation, regulation of melanogenesis, melanosome biogenesis and trafficking, melanosome transfer to keratinocytes, and melanin biosynthesis (**Fig. 2B**).

**Figure 2.**
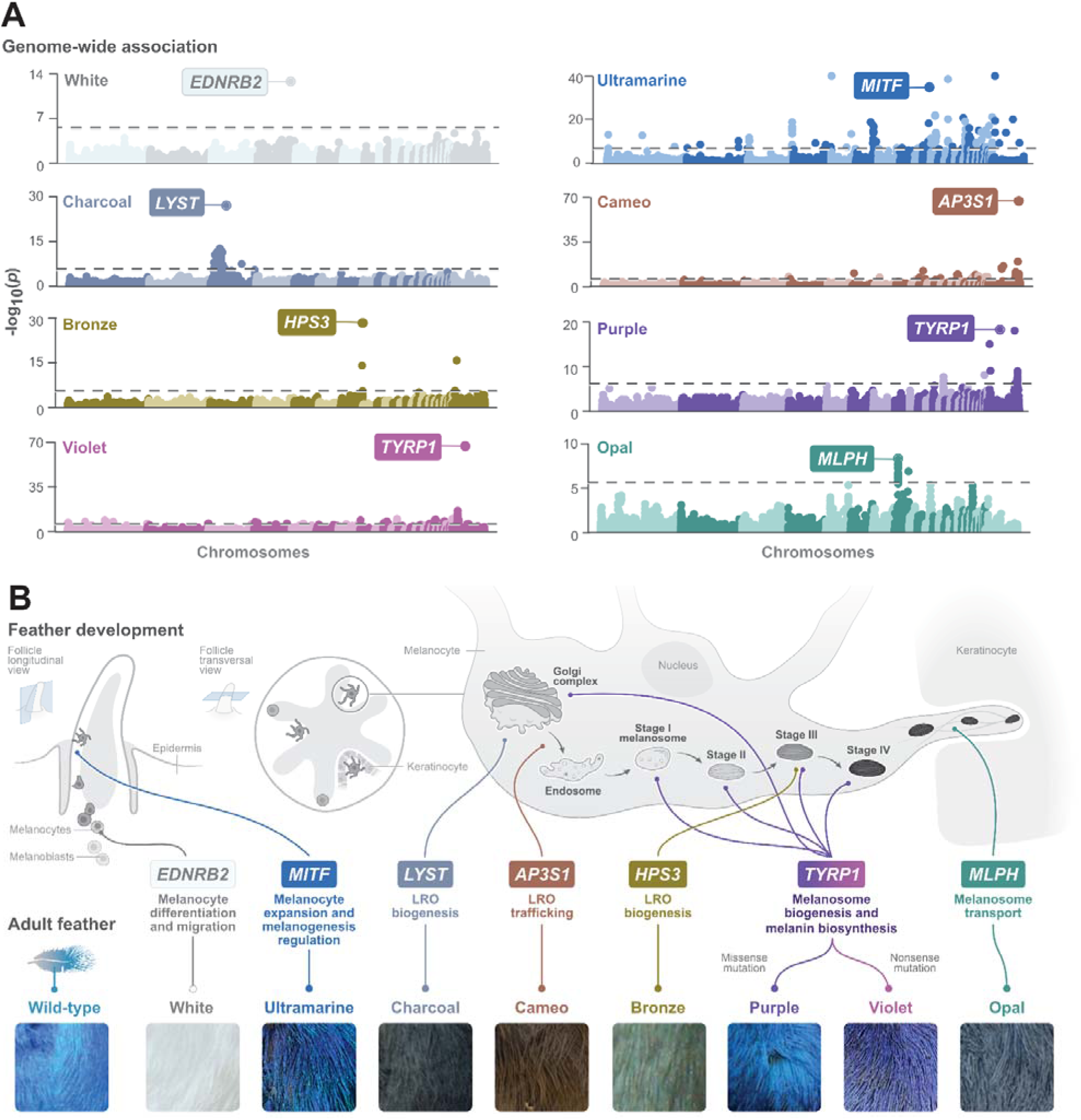
Genetic mapping of color variants. **(A)** Manhattan plots representing the genome-wide association analyses (GWAS) for plumage color. Each point represents a variant, with the *x*-axis showing genomic position (chromosomes in alternating colors, ordered according to the Indian peafowl reference genome) and the *y*-axis indicating statistical significance as –log_10_(*p*). The horizontal dashed line indicates the Bonferroni-corrected genome-wide significance threshold. The most significantly associated variant in each plot is annotated with the name of the gene in which it resides. (**B)** Schematic overview of melanogenesis in feather development, highlighting candidate genes identified by GWAS. On the top left, longitudinal and cross-sectional views of a developing feather follicle depict the spatial arrangement of melanocytes during feather formation. In the center and top right, a melanocyte is illustrated with its key cellular components involved in melanogenesis, the melanosome maturation stages, and the dendritic projections connecting to keratinocytes, where melanosomes are finally deposited. Colored lines link genes to their respective subcellular locations and cellular processes associated with melanocytes, lysosome-related organelles (LRO), and melanosomes. *TYRP1* is associated with two phenotypes and the mutation type (e.g., missense, nonsense) is indicated. Along the bottom, magnified images of adult neck feathers (∼2 × 2 cm) display the macroscopic color phenotypes corresponding to each mutation.

Within each candidate region, we identified putative causal variants, including missense substitutions, premature stop codons, and exon deletions (**table S2**). In several cases, their functional relevance was supported by expression data or their overlap with evolutionarily conserved amino acid positions across avian species (**fig. S4**; **table S2**). Genotyping across our full cohort of samples revealed that four additional peafowl varieties result from combinations of these mutations (**table S2**), illustrating how epistatic interactions further expand color diversity. Collectively, these results show that variation in melanogenesis genes is associated with a wide range of effects on iridescent color phenotypes in bird feathers, and establish a basis for linking specific mutations to feather nanostructure and optical phenotype.

### Photonic lattice geometry is highly sensitive to changes in melanogenesis

Next, we hypothesized that genetic variation in peafowl influences feather coloration by altering the organization of photonic nanostructures. To test this, we sampled six chromatic zones of the peacock eyespot (A1-A6; **Fig. 3A**) (27, 30, 37), and examined cross-sectional and longitudinal preparations of barbules from wild-type and the genetically characterized varieties using transmission electron microscopy (TEM). We quantified seven structural parameters that collectively describe the nanostructural organization of the barbule photonic lattice, encompassing melanosome geometry, air pocket dimensions, and lattice organization (**Fig. 3A; fig S5; table S3**). We related ultrastructural features to optics via angle-dependent reflectance spectroscopy, which measures how feather color changes when viewed or illuminated from different angles.

**Figure 3.**
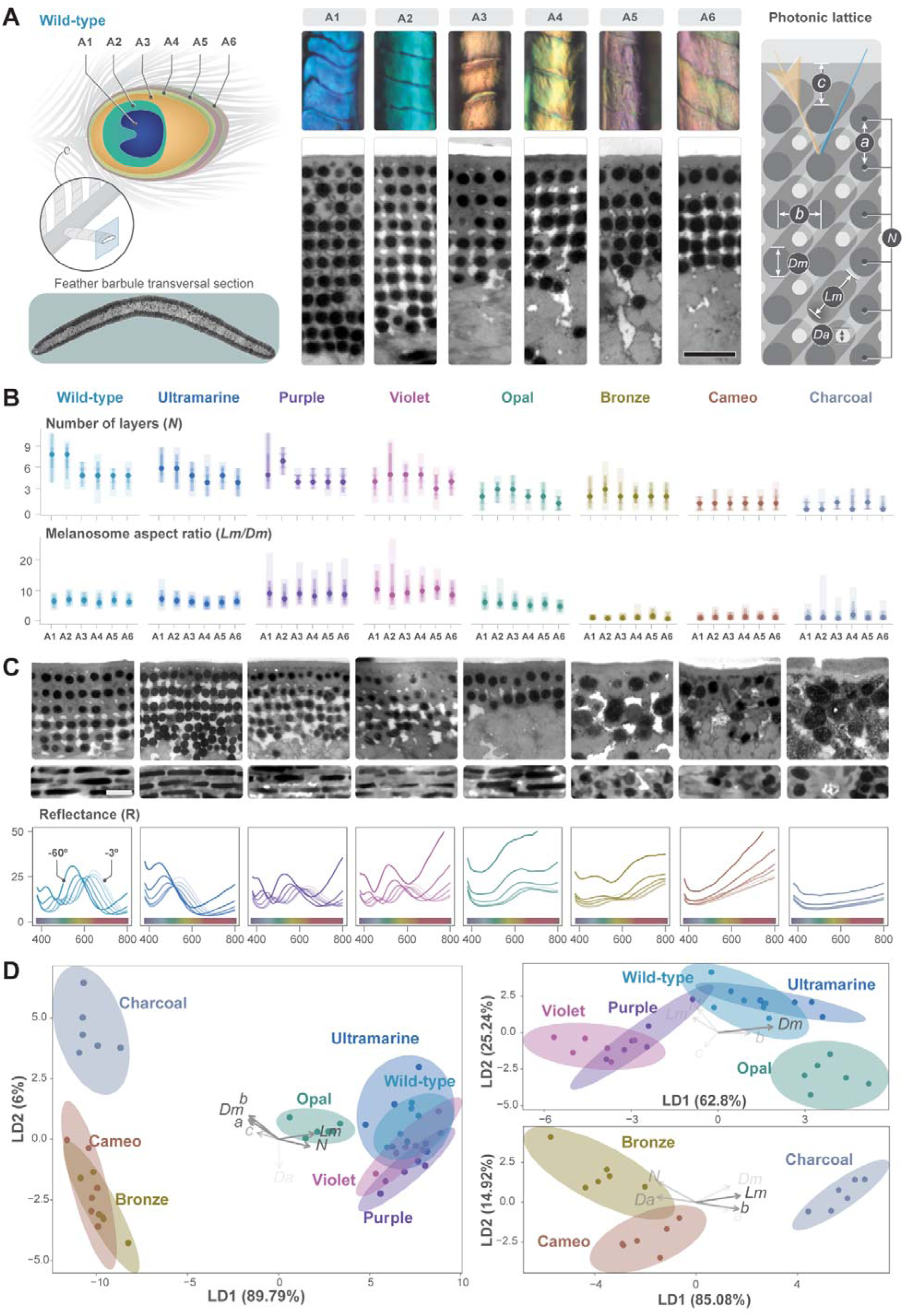
Ultrastructural modifications and photonic impact in eyespot feathers. **(A)** Schematic representation of feather structure, sampling procedure, and measured parameters of the photonic lattice. Left: A diagram of the wild-type peacock eyespot, divided into six chromatic regions (A1–A6), alongside a transverse section of a feather barbule exemplifying the transmission electron microscopy (TEM) imaging used for structural analysis. Center: TEM images illustrating the ultrastructural arrangement of melanosomes across the six regions corresponding to different barbule colors in wild-type feathers (scale = 500 nm). Right: a 3D schematic representation of the key structural measurements from the photonic lattice of feather barbules, including: distance between melanosome layers (*a*); distance between melanosomes within layer (*b*); cortex thickness (*c*); melanosomes diameter (*Dm*); air pocket diameter (*Da*); number of melanosome layers (*N*) and length of melanosomes (*Lm*) (scale bar = 500 nm)**. (B)** Gradient plots illustrate differences in melanosome layer number (*N*) and aspect ratio (*Lm*/*Dm*) across A1–A6 in wild-type and mutants. Each circle represents an average measurement for a specific region, narrow bars represent quantile intervals at the 80% and 95% levels, and the wider bar represents the range of minimum and maximum for each area. Additional parameters are presented in **fig. S6**. **(C)** TEM of transversal (top) and longitudinal (bottom) cuts of feather barbules in A3 in wild-type and seven mutants (scale bar = 500 nm). Images for the remaining eyespot regions are available in **fig. S5**. Below, reflectance (R) profiles of A3 and variation (minimum– maximum) of the peak reflectance wavelength of the angle-dependent measurement from -60° to -3° (in increasing transparency). Measurements were taken at 10° intervals, with the exception of an additional measurement at −3° to capture the reflection near the surface normal. **(D)** Linear discriminant analyses (LDA) of structural measurements for all mutants on the left, and for iridescent (top) and non-iridescent (bottom) mutants on the right, with oval shading representing the variation per mutant across areas. The arrows represent the parameters explaining a significant amount of variance in LD1 of each analysis (*P* < 0.01), and the shading reflects their individual weight.

TEM imaging of wild-type feathers revealed that rod-shaped melanosomes formed a highly ordered photonic lattice, with their long axes aligned parallel to the barbule, with variation in layer spacing and number generating the distinct hues across the eyespot (**Fig. 3A; fig. S5**), as previously documented (27, 29, 37). In contrast, color mutants exhibited pronounced deviations from the wild-type ultrastructure (**Fig. 3A- C; figs. S5 and S6**), and we applied linear discriminant analysis (LDA), a statistical approach that identifies which variables best distinguish among groups, to determine which features most strongly contribute to these differences (**Fig. 3D; tables S4 and S5)**. This analysis revealed a clear separation between two main clusters along the first discriminant axis (LD1), which accounted for 90% of total variance (**Fig. 3D**). Melanosome length was the strongest contributor (*Lm*, *P* = 6.1×10^−37^) (**table S4**), which was inversely correlated with melanosome diameter (*Dm*) (*r* = –0.76), together defining the melanosome aspect ratio (rod-like versus spherical). Bronze, Cameo, and Charcoal mutants clustered on the negative LD1 axis, and TEM images revealed a collapse of the photonic lattice, with spherical and poorly ordered or disordered melanosomes replacing the aligned rods typical of wild-type iridescent feathers (**Fig. 3B and C; figs. S5 and S6**). Their reflectance spectra—measurements of the wavelengths of light reflected by the feather— were flat or displayed weak features (**Fig. 3C; fig. S7**), indicating a loss of constructive interference. Our results thus reveal a gradient among peafowl mutants in which progressive melanosome rounding or irregularity corresponds to increasing lattice disorder (strongly iridescent types < Bronze < Cameo < Charcoal) (**Fig. 3B and C**; **table S5**).

The Opal variety exhibited a partially ordered lattice with fewer layers, yielding weaker iridescence (**Fig. 3B and C; figs. S5-S7**). Violet, Purple, and Ultramarine mutants retained well-organized multilayered lattices (**Fig. 3C**; **fig. S5**) and clustered close to wild-type (**Fig. 3D**). Within this iridescent group, LD1 (62.8% of variance) was driven by melanosome diameter (*Dm*; *r* = 0.94, *P* = 1.2 × 10^−14^), with additional contributions from inter-melanosome spacing (*b*; *r* = 0.54, *P* = 0.002) and inter-layer distance (*a*; *r* = 0.38, *P* = 0.04) (**Fig. 3D**; **table S4**). LD2 (25.2% of variance) was shaped mainly by layer number (*N*; *r* = 0.81, *P* = 7.3 × 10^−8^). Comparative analyses relative to wild-type highlighted the phenotype-specific shifts (**table S5**). Opal differed most strongly in layer number (*N*; *P* = 1.1 × 10 ^4^), whereas Purple and Violet diverged mainly in melanosome diameter (*Dm*; *P* = 2.8 × 10 ^6^) and in inter-melanosome spacing within layers (*b*; *P* < 3.0 × 10 ^2^). Ultramarine showed differences only in inter-melanosomal spacing (*b*; P = 2.0 × 10 ^2^). The number of layers (*N*) also differed significantly in Violet (*P* = 4.4 × 10 ^2^) and, although not significant overall, exhibited pronounced regional variation in Purple and Ultramarine (**figs. S5 and S6**).

These ultrastructural changes were closely associated with optical variation. A blue shift in peak reflectance from Purple to Violet to Ultramarine (**fig. S7**) is associated with decreasing inter-melanosome spacing within layers, which shortens the optical path and shifts constructive interference toward shorter wavelengths. Across chromatic zones and iridescent varieties, layer number (*N*) was positively associated with color saturation (84%; *t* = 5.7, *df* = 94; *P* = 1.3 × 10^−7^), consistent with enhanced constructive interference in barbules with more layers (**fig. S8**). Thus, repetition (layer number) and periodicity (melanosome spacing) are tunable features of photonic lattices that vary predictably with mutations in melanogenesis genes and are associated with differences in hue, saturation, and spectral bandwidth in iridescent plumage. Our findings also reveal a continuum of nanostructural integrity among peafowl color variants, in which some mutations are associated with pronounced disruption of lattice organization, whereas others are associated with subtler changes in lattice geometry and color.

### Cellular and biochemical basis of iridescence

Because the candidate genes identified in peafowl are associated with melanogenesis, we next investigated how these mutations affect melanin production and other melanogenesis-related cellular processes underlying iridescent feather development. To determine how such changes shape feather nanostructure, we analyzed regenerating feather follicles and mature neck feathers from wild-type and mutant birds. Longitudinal follicle sections were examined by light microscopy to assess melanin distribution in melanocyte and keratinocyte layers, as well as the melanocyte dendritic branching (**Fig. 4A**), which mediates melanosome transfer into keratinocytes that become the feather’s structural matrix. In parallel, we performed chemical analysis of mature feathers to evaluate how mutations affect melanin biosynthesis and deposition (**Fig. 4B**; **table S6**). We focused on neck feathers because they can be induced to regenerate by plucking independently of the molting cycle and exhibit uniform coloration, providing a simplified and more tractable system than the complex color patterning of the eyespot and facilitating comparisons of cellular and biochemical phenotypes among varieties.

**Figure 4.**
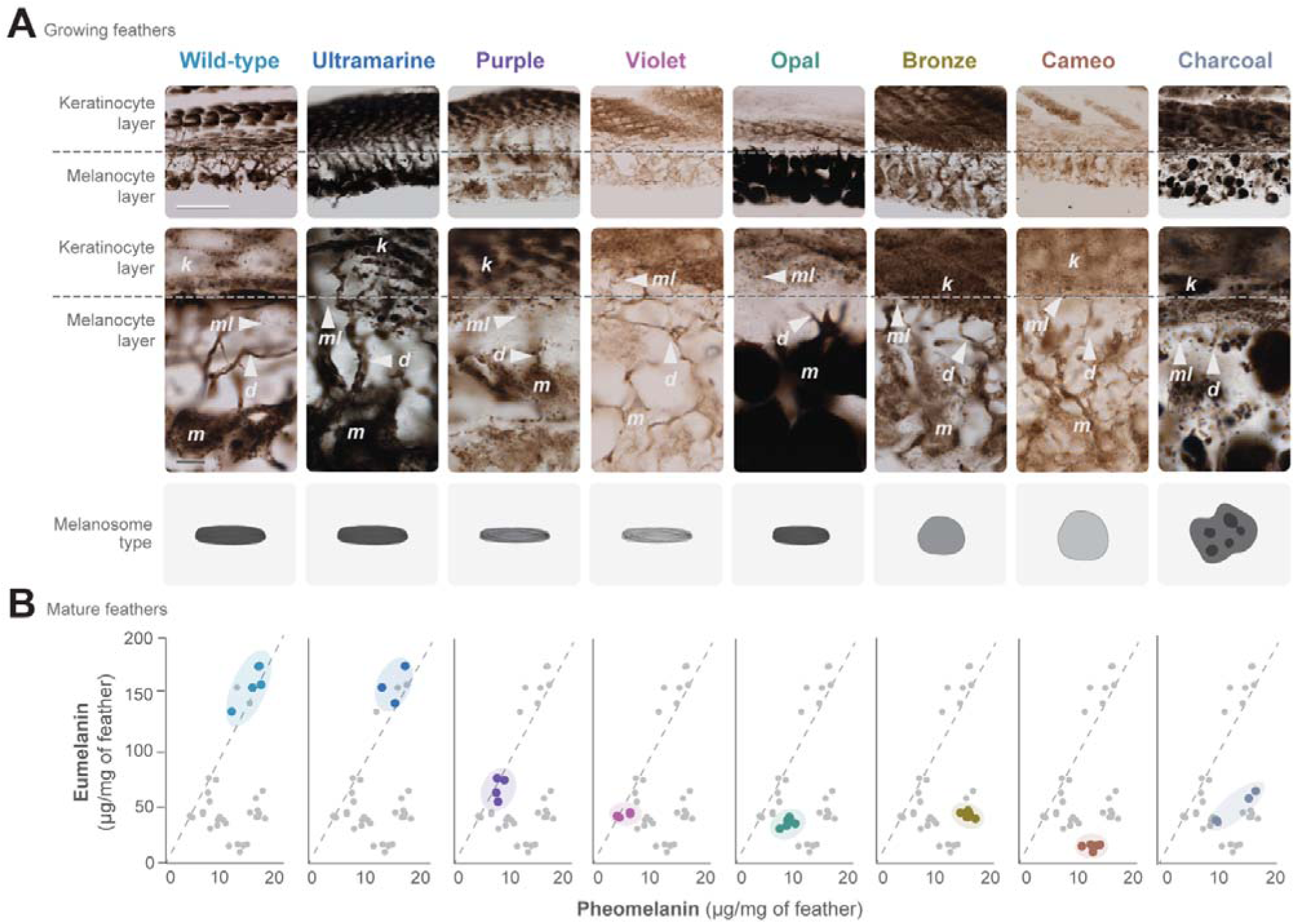
Histological and biochemical characterization of peafowl mutants. (**A**) Longitudinal sections of developing feather follicles of wild-type and mutant birds showing the organization of a melanocyte layer below the dashed line and a keratinocyte layer above the dashed line (top, scale bar = 50 µm). An enlarged view of the boundary between the two layers is shown (bottom, scale bar represents 10 µm), and various structures are highlighted: melanocytes (*m*), keratinocytes (*k*), dendrites (*d*), and melanosomes (*ml*). Across the bottom are schematic illustrations of melanosome morphology derived from TEM images (**fig. S5**). (**B**) Melanin quantification in mature neck feathers from wild-type and mutant phenotypes (detailed values provided in **table S6**). Graphs show quantification of pheomelanin (*x*-axis) and eumelanin (*y*-axis). All datapoints across all genotypes are plotted in each panel to enable direct visual comparison. Points belonging to the genotype shown in that panel are highlighted in color, while all other datapoints are shown in gray. The dashed line represents a eumelanin-to-pheomelanin ratio of 10:1.

In wild-type follicles, melanosomes were abundant and efficiently transferred from melanocytes to keratinocytes through dendritic projections, producing a uniform pigment distribution (**Fig. 4A**). Biochemically, wild-type feathers had a high eumelanin-to-pheomelanin ratio (∼10:1; **Fig. 4B**). The weakly iridescent (Bronze) and non-iridescent mutants (Cameo and Charcoal) showed diverse cellular and biochemical changes. In Charcoal follicles, dendrites and melanosome transfer were present, but melanosomes formed aggregates within melanocytes and keratinocytes (**Fig. 4A**). Despite appearing visually darkest, Charcoal contained 67% less total melanin than wild-type feathers (*t* = 8.4, *df* = 6, *P* = 1.6×10^−4^) and comparable levels to Violet and Purple (**Fig. 4B**), which appear far more vibrant, illustrating a disconnect between pigment abundance and perceived brightness arising from structural organization. Cameo and Bronze displayed disrupted melanin distribution, with clustered pigment deposits within keratinocytes indicating melanosome aggregation (**Fig. 4A**). In Cameo, pigment-free regions within melanocytes suggested the presence of large intracellular vacuoles. Importantly, across all three mutants, eumelanin content was markedly reduced, but the pheomelanin levels remained almost unaltered (**Fig. 4B**), hence reducing the eumelanin-to-pheomelanin ratio (∼1.5:1–4:1). These compositional shifts coincided with the transformation of rod-shaped melanosomes into more spherical forms (**Fig. 3B and C**). Although the underlying genes act at distinct steps of melanogenesis, none are known to directly regulate melanosome morphology. The repeated emergence of spherical melanosomes is instead consistent with the possibility that altered melanin composition influences melanosome shape, either directly or by perturbing normal melanosome maturation. These changes in melanosome shape, in turn, were consistently associated with compromised photonic lattice integrity.

The Opal mutant showed a distinct cellular phenotype, characterized by melanocytes densely filled with melanin but lacking visible dendritic projections to keratinocytes, which receive few melanosomes (**Fig. 4A**). This suggests disrupted melanosome transfer. Biochemically, Opal feathers exhibited a large reduction in total melanin (–73%; *t* = 11.4, *df* = 7, *P* = 8.9 × 10^−6^; **Fig. 4B**), but unlike Cameo, Bronze, and Charcoal, this decrease affects both eumelanin and pheomelanin (**Fig. 4B**). This more balanced loss appears to preserve the rod-shaped melanosomes typical of wild-type, in contrast to the spherical forms that characterize other low-melanin mutants. Consequently, despite melanin levels comparable to Charcoal, the retention of rod-shaped melanosomes seemingly permits partial nanostructural organization, making Opal appear lighter and more vivid in color.

Hue-shifted iridescent variants (Purple, Violet, and Ultramarine) showed feather follicles with cellular organization comparable to wild-type, with only slight differences (**Fig. 4A**). Purple and Violet follicles contained a higher proportion of light brown melanosomes, indicative of delayed or incomplete maturation as expected from *TYRP1* mutations (38), whereas Ultramarine follicles exhibited dense pigmentation. All three phenotypes maintained wild-type-like eumelanin-to-pheomelanin ratios (Purple: 9:1; Violet: 8.5:1; Ultramarine: 11.4:1) and retained the characteristic rod-shaped melanosomes (**Fig. 4B**).

Despite these similarities, these mutants fall into two distinct mechanistic categories. Purple and Violet feathers showed large reductions in total melanin content (Purple: –50%, *t* = 6.7, *df* = 6, *P* = 5.3 × 10^-4^; Violet: –72%, *t* = 9.4, *df* = 6, *P* = 8.5 × 10^−5^) (**Fig. 4B**), yet sufficient to support ordered multilayered structures (**Fig. 3C**). Both mutants also exhibited reduced inter-melanosome spacing (see above). Because *TYRP1* encodes a core enzyme in eumelanin synthesis and contributes to melanosome maturation rather than spatial organization (38), it is unlikely to directly regulate inter-melanosome packing. Instead, the packing changes are most plausibly an indirect consequence of the increased melanosome aspect ratios resulting from reduced melanosome diameter (*Dm*) in both mutants compared to wild-type (**Fig. 3B**). This is consistent with known effects of *TYRP1* on melanosome morphology (39). These observations suggest that variation in melanosome aspect ratio may represent an additional feature of melanogenesis associated with photonic assembly and optical performance. The observed differences between Purple and Violet feathers likely reflect an additional layer of interaction between melanosome abundance and melanosome shape.

In contrast, we found that Ultramarine feathers, likely caused by an *MITF* mutation, showed no significant change in pigment abundance (+0.06%, *t* = 4.9 × 10^−3^, *df* = 5, *P* = 0.1) (**Fig. 4B**), despite intense cellular pigmentation in the follicle sections (**Fig. 4A**). Analyses of melanosome morphology and pigment chemistry likewise revealed no detectable differences from wild-type (**Fig. 4B**; **fig. S6**), indicating that pigment quantity or composition cannot account for the observed optical properties. Our ultrastructural data show that the Ultramarine photonic lattice is densely packed, with significantly reduced inter-melanosome distance (*b*) (**Fig. 3C; figs. S5 and S6**). TEM images further indicated a unique organizational anomaly in Ultramarine, which is absent in wild-type and other mutants, consisting of clusters of disorganized melanosomes near the basal boundary of the photonic crystal and occasionally dispersed melanosomes within the medullary region of the barbule (**fig. S9**). The anomaly suggests a developmental mechanism in which photonic lattice assembly was prematurely arrested and melanosomes failed to integrate into the lattice within the appropriate temporal window. A loss of temporal coordination between melanosome delivery and lattice formation could also underlie the observed alterations in periodicity, possibly reflecting melanosome deposition during phases of altered keratin matrix viscosity. This interpretation aligns with several known functions of *MITF*, including its roles in regulating melanocyte differentiation and the timing of melanosome maturation (40).

Taken together, our biochemical, ultrastructural, and photonic analyses indicate that feather nanostructure is a developmentally plastic trait associated with melanosome availability and morphology, as well as with the temporal dynamics of pigment deposition during feather development.

### Single mutations in melanogenesis genes can induce iridescent plumage

The Black-shoulder phenotype is another domesticated peafowl variant (41). Unlike the previously described mutations that modify existing iridescent colors throughout the body, Black-shoulder produces a localized gain of iridescence in males within plumage regions that are normally weakly iridescent or non-iridescent. In wild-type peacocks, the shoulders and upper back display a light-dark barred pattern with little iridescence, whereas Black-shoulder males replace this pattern with vivid green and blue plumage (**Fig. 5A**). This mutation also has strikingly sex-specific effects: Black-shoulder females are predominantly depigmented and lack the normal brown body plumage and iridescence, appearing nearly white to pale cream due to pheomelanin deposition (**Fig. 5A**). Thus, the same mutation produces a localized gain of iridescence in males but a reduction in melanin pigmentation and iridescence in females. The male phenotype offers an opportunity to investigate the molecular and developmental basis of the emergence of novel biological photonic structures.

**Figure 5.**
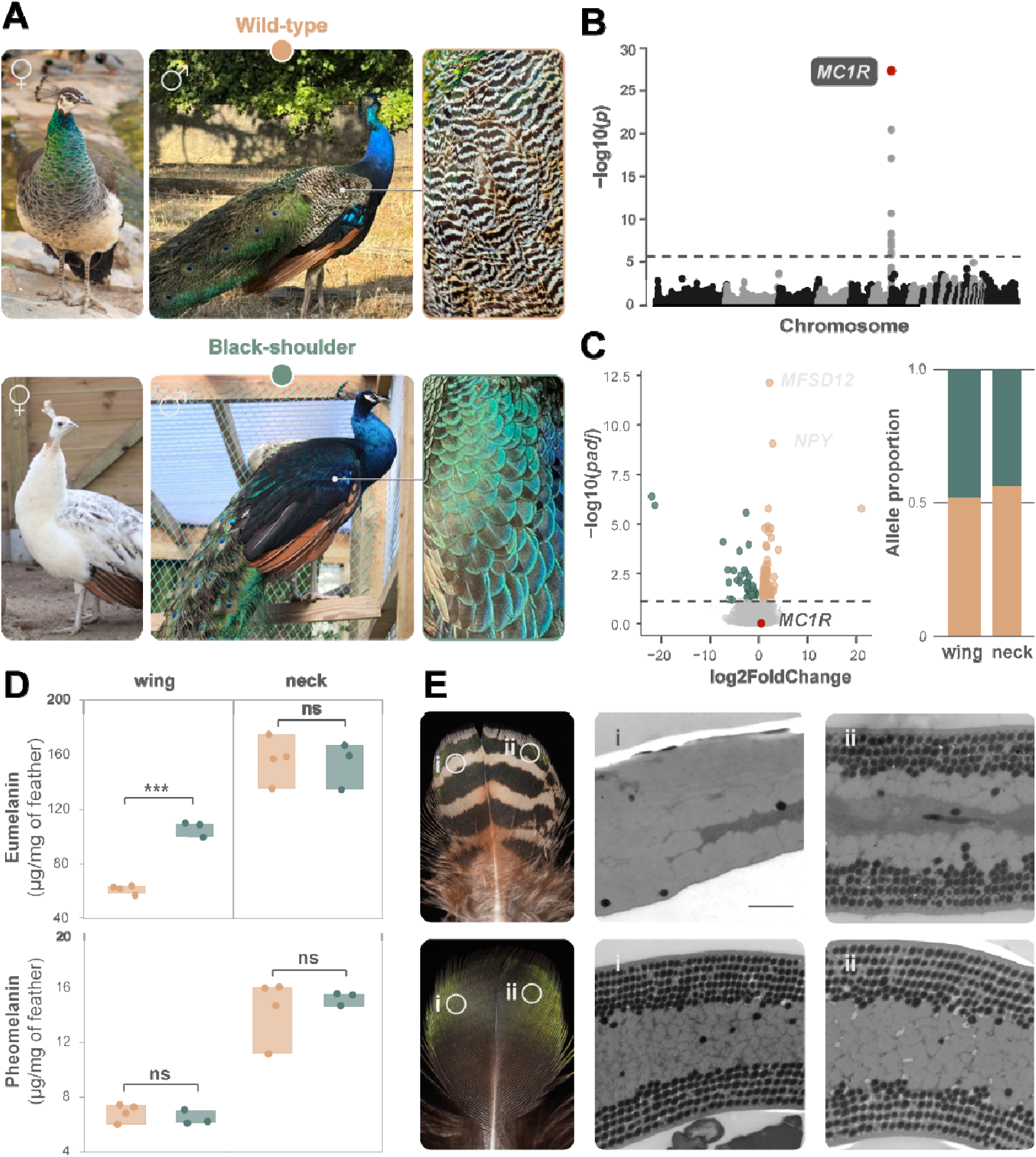
A patch-specific gain of iridescence. **(A)** Pictures illustrating wild-type (top) and Black-shoulder (bottom) peafowls, both females (peahens, left) and males (peacocks, center), and a close-up of peacock wing covert feathers (right). **(B)** Manhattan plot displaying whole-genome association results. Each dot represents the −log_10_ (*p*) value from the likelihood ratio test for a genetic variant. The red horizontal line indicates the Bonferroni-corrected genome-wide significance threshold. The strongest association corresponds to a missense mutation in *MC1R* (red circle). **(C)** Left panel: a volcano plot comparing RNA-seq expression profiles of regenerating feather follicles from the wing of wild-type (*n* = 4) and Black-shoulder (*n* = 4) males. The horizontal dashed line marks the significance threshold (FDR = 0.1). Green and beige circles represent genes upregulated in Black-shoulder and wild-type individuals, respectively. The red circle represents the *MC1R* gene. Significantly differentially expressed genes with known functions in melanogenesis are labeled in grey. Right panel: allelic imbalance analysis of the wild-type (beige) and Black-shoulder (green) *MC1R* alleles in feather follicles from the wing and neck of male individuals heterozygous for the missense mutation associated with the Black-shoulder phenotype. **(D)** Comparisons of eumelanin (top) and pheomelanin (bottom) levels in wing and neck feathers of wild-type (beige) and Black-shoulder (green) male feathers (detailed values provided in **table S6**). Significant differences are indicated (*** *P* < 0.001), while "ns" denotes non-significant differences (*P* > 0.05), as determined by a two-sample *t*-test assuming unequal variances. **(E)** On the left, representative wing covert feathers from wild-type (top row) and Black-shoulder (bottom row) males, highlighting equivalent locations chosen for ultrastructural analyses of feather barbules. The center and right panels show TEM images of transversal cuts of feather barbules from the selected regions, illustrating melanosome organization, and where scale represents 1 µm and is maintained across figures.

To identify the causal variant, we conducted a GWAS comparing Black-shoulder (*n* = 107) and wild-type birds (*n* = 244, scored for wing color phenotype). This analysis revealed a significant association (*P* = 5.0 × 10 ^28^) overlapping the *MC1R* gene (**Fig. 5B; fig. S10**), which encodes a receptor that regulates melanin synthesis via the cAMP pathway (42). We next assessed gene expression in regenerating wing covert feather follicles from Black-shoulder (*n* = 4) and wild-type (*n* = 6) males using RNA-seq. *MC1R* expression was similar between phenotypes (**Fig. 5C**), suggesting that functional differences arise from coding changes rather than changes in gene regulation, despite the patch-specific effects. Examination of the *MC1R* coding sequence revealed two mutations: a missense mutation (Trp251Arg and the GWAS lead variant) and a frameshift mutation (**table S2**). Genotyping showed that Black-shoulder individuals were either homozygous for one mutation or compound heterozygous, indicating genetic heterogeneity underlying the same phenotype. If the two variants resided on the same haplotype, they would always be inherited together, making individuals homozygous for only one of the two mutations impossible. Black-shoulder individuals homozygous for the missense mutations were always wild-type for the frameshift mutations and vice versa, which demonstrates that the variants reside on distinct haplotypes. Because the missense allele is substantially more frequent among Black-shoulder individuals (76.8%), it generates the strongest association signal in the GWAS. Both mutations are predicted to disrupt *MC1R* function, given its canonical effects of eumelanin loss in females (**Fig. 5A**). The frameshift alters all amino acid residues of transmembrane domain (TM) 4 and 5 and a premature stop codon excludes TM 6 and 7 from the MC1R protein. The missense mutation affects an amino acid residue that is universally conserved among birds (**table S2**). The very similar phenotypes associated with both alleles suggest that they alter *MC1R* signaling in a similar manner, but the specific functional consequences of each mutation remain to be experimentally validated and considered in the context of the expanding functional landscape of *MC1R* in vertebrates (43–44). Allelic imbalance tests, which assess whether one gene copy is expressed more strongly than the other in heterozygote birds, showed no expression bias (**Fig. 5C**), reinforcing that the phenotype arises from altered protein function.

To understand the biochemical basis of this gain of iridescence, we analyzed melanin content as above. Chemical analyses of male wing covert feathers showed a ∼1.7-fold increase in eumelanin in Black- shoulder relative to the wild-type (*t* = 12.3, *df* = 3, *P* = 4.0 × 10^−5^), whereas eumelanin in neck feathers and pheomelanin in either region showed no significant difference (*t* < 0.4, *df* = 3-5, *P* > 0.4; **Fig. 5D**). Because these measurements represent the total eumelanin content of whole feathers, they cannot distinguish whether the increased eumelanin in wing covert feathers reflects increase melanogenesis or simply a greater proportion of the feather undergoing eumelanization. In light of what is known about *MC1R* function in mammals and some birds (45–46), the effect of the Black-shoulder mutations is surprising. Loss-of-function mutations in *MC1R* or mutations that reduce its activity are typically associated with red or lighter pigmentation and tend to affect pigmentation broadly across the body rather than in localized regions. This expected behavior is in fact observed in Black-shoulder females, which are lightly colored and retain pheomelanin pigmentation. The striking contrast between peacocks and peahens, therefore, suggests that *MC1R* can interact with sex-specific developmental programs to produce sexually dimorphic effects on pigmentation and iridescence. A growing number of studies in birds have reported similar effects of *MC1R* that differ from the canonical mammalian model (47–50) (see Discussion).

To determine the photonic basis of this gain of iridescence in the wings of Black-shoulder males, we next examined feather ultrastructure. TEM imaging showed organized rod-shaped melanosomes in dark bands of wild-type feathers, but light bands were sparsely pigmented and lacked discernible structure (**Fig. 5E**). In the feathers of male Black-shoulder, in contrast, melanosome density and layering were often increased toward feather tips, and the multilayered organization extended into regions corresponding to the wild-type light bands. Both features show an increase in ordered ultrastructure that underpins the enhanced photonic properties of Black-shoulder feathers (**Fig. 5E**). These findings suggest that melanin deposition beyond a threshold can trigger the formation of ordered multilayered nanostructures, even in feather barbs previously devoid of such structures. Thus, peafowl, and perhaps other birds, may possess a latent capacity to form photonic nanostructures that can be triggered by changes in the melanogenesis pathway.

### Transcriptomic drivers of iridescence in nature

To investigate the developmental basis of naturally occurring iridescence and the formation of its underlying photonic structures, and to determine whether melanocyte-centered processes more broadly contribute to this trait in nature, we performed single-cell RNA sequencing (scRNA-seq) on regenerating feathers of the emerald dove (*Chalcophaps indica*) and mallard duck (*Anas platyrhynchos*). These species represent two additional avian orders and develop asymmetric feathers in their wings, where iridescent and non-iridescent barbules differentiate on opposite sides of the same feather (**Fig. 6A**). This within-feather design enables direct comparisons of cell composition and gene regulation between barbule types within an otherwise identical developmental context. TEM confirmed ordered nanostructures within iridescent barbules and their absence in non-iridescent ones (**Fig. 6A**; **fig. S11**). To identify molecular drivers of these differences, we microdissected regenerating follicles to isolate the sides expressing iridescent and non-iridescent colors and generated eight scRNA-seq libraries (two replicates per barbule type per species).

**Figure 6.**
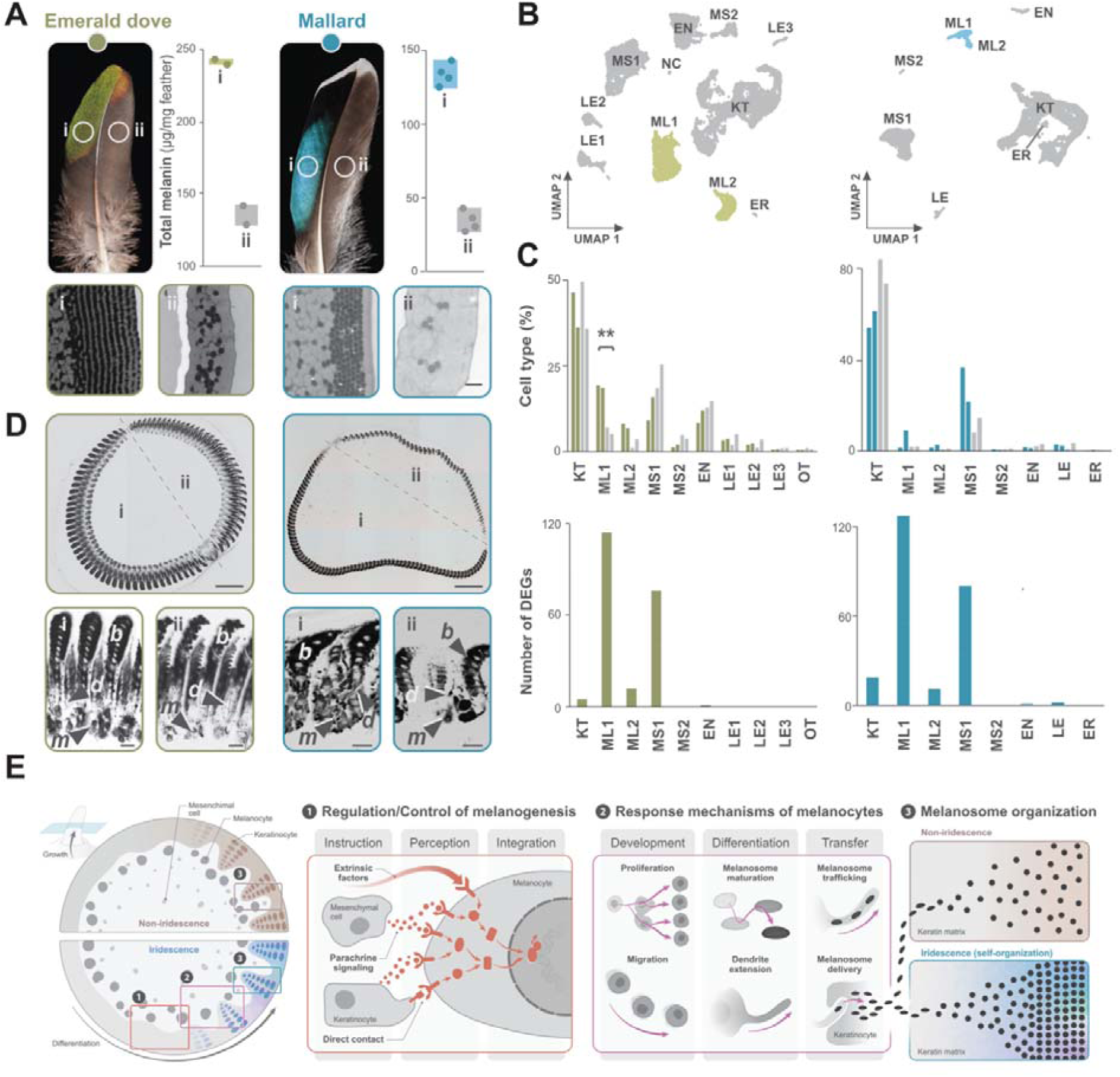
Modulation of iridescence via melanogenesis in wild bird species. The emerald dove (*Chalcophaps indica*) and the mallard duck (*Anas platyrhynchos*) develop asymmetric feathers containing both iridescent (i) and non-iridescent (ii) barbules. **(A)** *Top:* total melanin content per sample in each of the two feather regions. *Bottom:* TEM images of barbule cross-sections in iridescent and non-iridescent barbules (scale bar for all panels: 1 µm). **(B–C)** *scRNA-seq* of developing feathers from emerald dove (left) and mallard (right): ML, melanocytes; KT, keratinocytes; MS, mesenchymal cells; EN, endothelial cells; LE, leukocytes; ER, erythrocytes; NC, neural crest-derived cells; OT, others (ER+NC) **(B)** UMAP projections showing cell clustering. **(C)** *Top:* proportion of major cell types identified. *Bottom:* number of differentially expressed genes (DEGs) per cell type. **(D)** *Histological analyses* of developing feathers. *Top:* cross-sections of feathers showing overall morphology (scale bars: left = 200 µm; right = 500 µm). The dashed line indicates the boundary within the regenerating feather follicle separating regions that give rise to iridescent (i) and non-iridescent (ii) barbules. *Bottom*: high-magnification images of barb ridges (scale bar = 20 µm) reveal increased melanization in melanocytes (*m*) and their dendrites (*d*), associated with iridescent barbule keratinocytes (*b*). **(E)** Schematic summary of proposed mechanisms controlling iridescent feather color formation inferred from the scRNA-seq data. Cross-section representation of growing feathers developing non-iridescent (top) or iridescent (bottom) colors. Throughout barbule differentiation, depicted as a gradient from left to right, melanocytes receive, perceive, and integrate environmental cues from neighboring cells and extrinsic factors (e.g., paracrine signals or growth factors) (1). These instructions modulate melanocyte behavior (2), and the resulting availability, shape, composition, and transfer of melanosomes influence their assembly into photonic structures (3).

We identified 31,204 high-quality cells representing all major cell types in growing feathers (**Fig. 6B; figs. S12-15**). Melanocytes formed two transcriptionally distinct clusters corresponding to eumelanin- (melanocytes 1 or ML1) and pheomelanin-producing (melanocytes 2 or ML2) subtypes (see Methods). Using all cell populations, we compared iridescent and non-iridescent barbules to identify differentially expressed genes (DEGs) and differences in cellular composition. In the emerald dove, melanocytes exhibited the most noteworthy changes in both cellular composition and transcriptional profile (**Fig. 6C**). They were threefold more abundant in iridescent barbules (26.7% vs. 8.6%; *P* = 1.7 × 10^-2^) and showed the largest number of DEGs. This expression divergence was independent of cell-type abundance and more pronounced within the eumelanin-producing subtype ML1. In mallard, we saw no consistent compositional differences, yet ML1 again showed the largest transcriptional divergence in number of DEGs (**Fig. 6C**). We validated these findings with histological analysis of regenerating feathers and pigment quantification in mature feathers, which confirmed increased cellular pigmentation and greater melanin deposition in the iridescent side (**Fig. 6A and D**). These results suggest that cellular and transcriptional changes in melanocytes are central to iridescent color development in wild birds, consistent with our findings in peafowl.

Because melanocytes showed the strongest transcriptional divergence, we first focused on their DEGs (**Data S1 and S2**). When the melanocyte subclusters were analyzed either separately or together, only eight genes were shared between both species (*CCZ1*, *CHN1, DCLK1*, *DCX*, *DLC1*, *FTH1*, *GPNMB*, and *NCALD*), indicating that iridescent coloration is produced by distinct gene sets in each lineage despite their convergent nanostructural architecture and optical function. None of the DEGs encoded melanin-synthesis enzymes; instead, expression changes implicated melanocyte signaling, ion homeostasis, melanosome dynamics, and dendritic transfer. In the emerald dove, v-ATPase subunits (*ATP6V1E1*, *ATP6V0A1*), the cation channel *TPCN2*, and the potassium channel *KCNJ13* showed differential expression, consistent with active tuning of melanosome pH and ion balance (processes essential for melanogenesis) in iridescent barbules. In mallard, *cAMP*-responsive effectors downstream of *MC1R* (a DEG in the dove) were elevated (*RAPGEF4* and *ARPP21*), along with trafficking and autophagy regulators (*UVRAG*) and cytoskeletal/adhesion genes involved in dendrite elaboration and vesicle transport (*TMSB4X*, *CHN1*, *KANK1*, *LPP*). Together with increased *GPNMB* and *CCZ1* expression (DEGs shared between both species), these signatures point to more active melanosome maturation and intracellular trafficking as a common feature of iridescent barbule formation. Intercellular signaling also shifted in iridescent feathers and was reflected in gene ontology (GO) enrichment (**fig. S16**): genes associated with GTPase–mediated and ephrin- receptor signaling were elevated in mallard iridescent barbules (*RAPGEF4, CHN1, EPHA7*); WNT-axis components were reduced in both species (*DAAM2* in mallard; *LGR4* in dove); and integrin signaling decreased in dove (*ITGA4*). Three of the genes shared by both species (*CHN1*, *DCLK1*, *DCX*) and several top-10 DEGs (*NBEA*, *VAT1*/*VAT1L*) were linked to neuronal and synaptic processes (GO analyses also confirm an enrichment of many neuron/synapse-related terms; **fig. S16**), raising the hypothesis that synaptic- like vesicular processes contribute to differences in melanosome transfer along melanocyte dendrites between barbule types. These data indicate extensive rewiring of melanocyte biology between iridescent and non-iridescent barbules, while expression of the core enzymatic machinery of melanogenesis remains largely unchanged.

To assess gene regulation beyond melanocytes, we examined DEGs in keratinocytes and mesenchymal cells (**Data S1 and S2**). Many were functionally linked to melanogenesis through their regulatory effects on melanocyte behavior (51, 52), indicating that altered cross-talk between pigment cells and neighboring tissues distinguishes iridescent from non-iridescent barbules. Among the 15 keratinocyte DEGs with well- characterized proteins, six have known connections to melanocyte regulation, including paracrine factors upregulated in iridescent barbules that enhance melanocyte recruitment and proliferation (*KITL* and *NRG1* in mallard) and adhesive molecules mediating keratinocyte-melanocyte communication (*COL17A1* in dove). Mesenchymal cell DEGs likewise showed transcriptional signatures of differential paracrine regulation with secreted factors known to be implicated in melanocyte differentiation, migration, and pigment output, including *NDP* and *BMP2* in dove and *ASIP* and *FST* in mallard. Finally, several DEGs may contribute to iridescence through effects on the keratin matrix and tissue mechanics (11). For example, keratinocyte DEGs included feather structural components encoded by keratin genes that were consistently downregulated in iridescent barbules (*KRT1L*, *KRT*β*L*, and *KRT6A*). Taken together, our findings suggest that multiple developmental processes contribute to the formation of feather photonic structures in nature, with melanogenesis-associated changes representing one important component (**Fig. 6E**).

## Discussion

The formation of iridescent coloration requires an astonishing degree of structural precision. By analyzing nine peafowl color mutations, we connect genetic and biochemical variation to specific macroscopic optical properties. Some mutations produce whole-body losses or shifts in iridescent coloration, whereas Black- shoulder generates a notable, localized gain in iridescence restricted to particular feather tracts in males. Across these phenotypes, we genetically mapped mutations in melanogenesis genes that alter the composition, shape, and organization of photonic nanostructures, producing color shifts ranging from subtle hue differences to complete loss or gain of iridescence. To assess the formation of iridescent coloration in nature, we complemented the analyses of domesticated peacocks with single-cell transcriptomic studies of two wild species with asymmetric feathers, in which iridescent and non-iridescent barbules develop on opposite sides of the same feather. This within-feather comparison is informative because both barbule types develop in the same feather and therefore share much of their broader developmental context. These comparisons likewise highlighted extensive changes in melanocyte biology associated with iridescence, particularly in pathways related to melanosome maturation, intracellular transport, and communication between melanocytes and neighboring cells. Although other cellular processes can influence feather nanostructures (11, 26, 35, 53)—and our single-cell data in wild species reveal signatures of some of these processes, including expression changes in genes encoding keratins—our findings in domesticated and natural systems suggest that melanogenesis is one important contributor to variation in iridescent color. We propose that melanogenesis may be effective at tuning iridescence because its multistage pathway, governed by dozens to hundreds of interacting factors (45), constitutes a broad mutational target in which even minor perturbations can induce substantial nanostructural and optical shifts, as illustrated by the peafowl mutants.

In contrast to comparative studies in several bird species, our analysis leverages the shallow genetic divergence among peafowl varieties that exhibit cellular, nanostructural, and optical changes arising from single mutational events. The genes identified in mutants that reduce or abolish iridescence, Cameo (*AP3S1*), Bronze (*HPS3*), and Charcoal (*LYST*), span several stages of melanogenesis, including melanosome biogenesis and trafficking, and melanosome maturation. Despite encoding genes with different functions, these mutants were consistently associated with changes in melanin composition, particularly the balance of pheomelanin and eumelanin. Whenever eumelanin declined while pheomelanin remained largely unchanged, the characteristic rod-shaped melanosomes of the wild type were replaced by a more rounded, spherical form, indicating that altered melanin composition may influence melanosome morphology without additional genetic changes directly specifying organelle shape. This observation has been previously described in mammals and other birds (54, 55), where shifts in the eumelanin-to-pheomelanin ratio have also been associated with changes in melanosome shape. In the three peafowl mutants described above, spherical melanosomes were in turn associated with disruption of the photonic lattice and replacement of ordered multilayers by haphazard arrangements that produce black or brown plumage. A useful contrast is provided by the Opal variety, which is controlled by a gene thought to mediate the transport of melanosomes from melanocytes to keratinocytes (*MLPH*). Despite greatly reduced overall pigment levels, Opal feathers exhibit a higher eumelanin-to-pheomelanin ratio that is close to the wild-type, retain rod-shaped melanosomes, and preserve a partially ordered lattice together with residual iridescence. These results suggest that melanosome shape, which is at least in part determined by melanin composition, is an important determinant of photonic ultrastructure organization. This may reflect the fact that rod-shaped melanosomes self-assemble more efficiently into ordered lattice arrangements than spherical ones, possibly through depletion attraction forces, which act more strongly on elongated particles, like rods, to promote their alignment and regular packing. More broadly, the association between melanin composition, melanosome shape, and lattice integrity observed across these single-gene peafowl variants may help explain why pheomelanin-rich feathers rarely display iridescence across birds (11, 25) and suggests that shifts in melanogenesis could lead to the rapid loss of iridescence.

In cases where photonic lattices are preserved, we identified several additional genes in peafowl that modify melanin abundance and produce subtler changes in rod-shaped melanosome morphology. These include *MITF*, a key transcriptional regulator of melanocyte development and melanogenic gene expression, and *TYRP1*, an enzyme involved in eumelanin synthesis. Variation at these loci was associated with shifts in reflected wavelength and color intensity through their effects on lattice periodicity and melanosome stacking, illustrating how changes in melanosome shape, density, and arrangement can shift or generate new iridescent colors without disrupting the lattice itself. Our observations in the Ultramarine variant further raise the possibility that the temporal coordination of melanosome delivery also influences lattice organization. Together, these observations suggest that feather nanoarchitecture is a developmentally plastic trait, shaped by the cellular and biochemical environment rather than strictly specified by fixed genetic programs. In this context, the Black-shoulder mutation that maps to *MC1R* illustrates how changes in melanogenesis, in this case in males alone, can trigger the appearance of iridescent plumage in feather regions that normally show little or no iridescence, suggesting that some birds may possess a latent capacity for gains and shifts in iridescent coloration through changes in melanogenesis. This developmental plasticity is consistent with models in which photonic lattices emerge in part through physicochemical self- organization, where melanosome properties (shape, size, and density) have the potential to bias the outcome of lattice assembly during feather development (26).

This study also has several limitations. First, although our results in peafowl suggest that feather nanoarchitecture is developmentally plastic and responsive to changes in melanogenesis, we did not directly observe the formation of photonic lattices during feather growth. Ultrastructural analyses across successive stages of feather regeneration in both wild-type and mutant backgrounds will therefore be necessary to determine how changes in melanin production, melanosome maturation, and the timing of pigment delivery influence lattice assembly, and to clarify the extent to which direct cellular patterning or other developmental mechanisms also contribute. Second, our conclusions are based on spontaneously occurring genetic variants and correlative comparisons from chemical, to ultrastructural, to optical phenotypes that suggest a strong causality, but that do not provide direct evidence. Direct experimental manipulation, such as feather follicle culture combined with genome editing or virally mediated gene perturbation *in vivo*, offers a means to dissect the contribution of individual genes or pathways to feather formation. Paired with developmental time series and quantitative analyses of lattice assembly, such approaches could reveal how melanogenesis and other molecular processes (e.g., keratinization) shape photonic structure formation during barbule development. Third, the functional and mechanistic effects of the *MC1R* alleles associated with Black-shoulder plumage remain partially unresolved, and further work will be required to determine their effects on melanogenesis. Although many avian *MC1R* mutations produce pigmentation phenotypes consistent with the canonical mammalian model (45–46), in several bird species the relationship between *MC1R* and melanogenesis appears to be more heterogeneous, often regulating pigmentation in a spatially restricted manner within individual feathers or body regions (47–50). For example, in turkeys, an *MC1R* allele predicted to strongly impair receptor function causes increased dark pigmentation restricted to the wings rather than generalized pheomelanic plumage (48), closely resembling the peafowl Black-shoulder mutation. In chickens, *MC1R* contributes to the spatial organization of pigmentation within feathers rather than simply to overall melanin production across the body, with local inhibition of *MC1R* by *ASIP* generating alternating dark and light regions within feathers called barring (49), similar to the pattern seen in the wings of wild-type peacocks. In pigeons, the recessive Smoky mutation, caused by a large 500-bp deletion that most likely abolishes *MC1R* function, does not eliminate major eumelanic pattern elements such as the wing and tail bars, indicating that at least some aspects of avian eumelanin production are controlled by *MC1R*-independent pathways (50). The sex-specific and patch-specific effects of Black- shoulder make it a good system for dissecting how *MC1R* function is modified by hormonal and developmental context. Clarifying this will be informative not only for understanding iridescence gain in peacocks but also for understanding how birds regulate melanogenesis more generally. Finally, our study focuses on the most widespread class of avian iridescence, in which rod-shaped melanosomes form multilayered photonic structures. It therefore remains unclear to what extent similar melanogenesis- associated mechanisms, and the same degree of developmental plasticity, also contribute to other forms of iridescent coloration based on hollow or platelet-shaped melanosomes.

## Methods

### Genome sequencing and annotation of the Indian peafowl

A chromosome-level reference genome for the Indian peafowl (*Pavo cristatus*) was generated from a wild-type male using Oxford Nanopore long reads, Illumina linked reads, and Hi-C sequencing. Hybrid assembly, iterative scaffolding, and manual curation were used to produce the final assembly, and completeness was assessed with BUSCO. Genome annotation combined multi-tissue RNA-seq data, protein homology, repeat annotation, and *ab initio* gene prediction approaches to infer coding and non-coding gene models and assign functional annotations.

### Whole-genome resequencing and genetic mapping

Whole-genome Illumina resequencing was performed on 352 captive peafowl representing wild-type birds and multiple plumage color mutants, together with green peafowl (*Pavo muticus*) and hybrid individuals. Variants were identified using GATK (Broad Institute), annotated for predicted functional effects, and analyzed using mixed-model genome-wide association analyses to map loci associated with plumage phenotypes.

### Bulk RNA-sequencing

RNA-seq was conducted on regenerating feather follicles from wild-type and color mutants. Differential gene expression analyses were performed to identify transcriptional differences associated with feather coloration, and allele-specific expression analyses were used to quantify allelic imbalance at candidate loci.

### Feather imaging, spectroscopy, and ultrastructure analyses

Macroscopic imaging, optical microscopy, reflectance spectroscopy, and angle-dependent spectrometry were used to characterize feather coloration and optical properties or wild-type and color mutants. Feather nanostructure was examined with transmission electron microscopy, followed by quantitative measurements and multivariate statistical analyses of melanosome organization.

### Histology and melanin characterization

Histological sections of regenerating feather follicles were prepared to examine tissue organization during feather development. Chemical analyses of feathers were performed to quantify eumelanin and pheomelanin composition.

### Single-cell RNA-sequencing

Single-cell RNA-seq was performed on regenerating feathers from emerald dove (*Chalcophaps indica*) and mallard (*Anas platyrhynchos*) to compare iridescent and non-iridescent regions of asymmetric feathers during feather development. Dissociated cells were processed using the 10x Genomics platform, followed by clustering, cell-type annotation, tissue composition analyses, differential expression testing, and gene ontology enrichment analyses.

Detailed methods can be found on the Supporting Information document.

## Data availability

The whole-genome sequencing (Illumina), RNA sequencing, and single-cell RNA sequencing data generated in this study are available at the NCBI BioProject under accession number PRJNA1216069. The reference genome assembly and associated data, including Oxford Nanopore sequencing, Hi-C, and linked-read sequencing, are available under the ENA project accession PRJEB87866. The raw data supporting the nanostructural and optical analyses are available in the public repository Zenodo at the following DOI: https://doi.org/10.5281/zenodo.21456601.

## Acknowledgements

We thank the numerous breeders who kindly provided feather and tissue samples for data analyses. We are also grateful to Jens Poulsen, Natalia Gusakova (Unsplash), and Clucking Fens Farm (www.cluckingfens.co.uk) for peafowl photographs used in Fig. 1, Fig. 5, and fig. S1. We thank Ibanidis Lda (Versele-Laga distributor), ZooService Lda (Manitoba distributor), and Papa d’Ovo (Domus Molinari distributor) as sponsoring partners of food, husbandry cages, and accessories. We thank Joana C. Carvalho for helping produce the main figures. We thank Nico Nees for kindly providing the code used for the chroma calculations, Bettina Winzer for her assistance in aligning the angle-dependent spectrometer, and Carmen Rubach for helpful discussions. This work was supported by the European Research Council under the European Union’s Horizon 2020 research and innovation program to MC (grant agreement No. 101000504); by the Portuguese Foundation for Science and Technology (FCT, https://www.fct.pt) research contracts to MC (CEECINST/00014/2018/CP1512/CT0002), PA (2020.01405.CEECIND/CP1601/CT0011), and PMA (2020.01494.CEECIND); by research fellowships to CIM (SFRH/BD/147030/2019) and RAf (UI/BD/154451/2022), in the scope of the Biodiversity, Genetics and Evolution (BIODIV) PhD program; by the Deutsche Forschungsgemeinschaft (DFG, German Research Foundation; Project-ID 416229255 – SFB 1411, and Project-ID 539724755 – FPS Core Facility); by an EASI-Genomics project under CNAG-CRG coordination, funded by the European Union’s Horizon 2020 research and innovation programme (Grant Agreement No. 824110); by national funds through FCT to CE3C (UIDB/00329/2020, https://doi.org/10.54499/UIDB/00329/2020); and by funding from Vetenskapsrådet (VR 2017-02907), and the Knut and Alice Wallenberg Foundation (KAW 2023.0160 and KAW 2024.0058) to LA.

## Author Contributions

Conceptualization: M.C., N.V., J.C.C., L.A., S.B., C.B., R.A., P.M.A.

Methodology: S.B., P.M.A., P.P., P.A., R.Af., S.Af., J.Ca., R.J.L., J.M.A., M.F.A., R.A., C.B., M.P.J.N., J.B., A.Hi., A.He., E.S., B.A.Z., C.I.M., C.F., M.A.A., F.C., J.G.G., C.Z., M.G., T.A., S.I., K.W.

Investigation: S.B., P.M.A., P.P., P.A., R.Af., S.Af., J.Ca., R.J.L., J.M.A., M.F.A., R.A., C.B., M.P.J.N., J.B., A.Hi., A.He., E.S., B.A.Z., C.I.M., C.F., M.A.A., F.C., J.G.G., C.Z., M.G., T.A., S.I., K.W.

Data curation: S.B., P.M.A., P.P., P.A., R.A., C.B., J.B., A.Hi., A.He., E.S., B.A.Z., C.F., F.C., J.G.G., C.Z., M.G., T.A.

Formal analysis: S.B., M.F.A., R.A., P.P., C.B., M.P.J.N., J.B., A.Hi., A.He., E.S., B.A.Z.

Visualization: S.B., R.A., C.B., C.F., J.A.A., T.A., M.P.J.N., J.B., A.Hi., A.He., E.S., B.A.Z.

Supervision: M.C., N.V.

## References

1. 1. S. Kinoshita, Helen Ghiradella, Lars Olof Björn, “Spectral tuning in Biology II: Structural Color” in Photobiology, Björn L., Ed. (Springer, 2015).

2. I. C. Cuthill, et al., The biology of color. Science (1979). 357 (2017).

3. M. D. Shawkey, L. D’Alba, Interactions between colour-producing mechanisms and their effects on the integumentary colour palette. Philosophical Transactions of the Royal Society B: Biological Sciences 372, 20160536 (2017).

4. P. Vukusic, J. R. Sambles, Photonic structures in biology. Nature 424, 852–855 (2003).

5. 5. G. E. Hill, K. J. McGraw, Bird coloration. In Mechanisms and Measurements*, 1* (Harvard University Press, 2006).

6. T. F. Cooke, et al., Genetic mapping and biochemical basis of yellow feather pigmentation in budgerigars. Cell 171, 427–439 (2017).

7. R. Arbore, et al., A molecular mechanism for bright color variation in parrots. Science (1979). 386 (2024).

8. M. A. Gazda, et al., A genetic mechanism for sexual dichromatism in birds. Science (1979). 368, 1270–1274 (2020).

9. R. Price-Waldman, M. C. Stoddard, Avian coloration genetics: recent advances and emerging questions. Journal of Heredity 112, 395–416 (2021).

10. 10. R. O. Prum, “Anatomy, physics, and evolution of avian structural colors” in Bird Coloration, Volume 1: Mechanisms and Measurements, G. E. Hill, K. J. MCGraw, Eds. (Harvard University Press, 2006).

11. V. Saranathan, C. Finet, Cellular and developmental basis of avian structural coloration. Curr. Opin. Genet. Dev. 69, 56–64 (2021).

12. M. C. Stoddard, R. O. Prum, How colorful are birds? Evolution of the avian plumage color gamut. Behavioral Ecology 22, 1042–1052 (2011).

13. M. F. Land, The physics and biology of animal reflectors. Prog. Biophys. Mol. Biol. [Preprint] (1972).

14. C. M. Eliason, R. Maia, M. D. Shawkey, Modular color evolution facilitated by a complex nanostructure in birds. Evolution (N. Y). 69, 357–367 (2015).

15. R. Maia, D. R. Rubenstein, M. D. Shawkey, Key ornamental innovations facilitate diversification in an avian radiation. Proceedings of the National Academy of Sciences 110, 10687–10692 (2013).

16. M. Nicolai, et al., The evolution of brilliant iridescence in birds. [Preprint] (2025). 10.21203/rs.3.rs-8020267/v1

17. K. K. Nordén, et al., Melanosome diversity and convergence in the evolution of iridescent avian feathers- Implications for paleocolor reconstruction. Evolution (N. Y). 73, 15–27 (2019).

18. C. M. Eliason, et al., Transitions between colour mechanisms affect speciation dynamics and range distributions of birds. *Nat*. Ecol. Evol. 8, 1723–1734 (2024).

19. M. P. J. Nicolaï, et al., The evolution of multiple color mechanisms Is correlated with diversification in sunbirds (Nectariniidae). Syst. Biol. 73, 343–354 (2024).

20. R. Dakin, R. Montgomerie, Eye for an eyespot: how iridescent plumage ocelli influence peacock mating success. Behavioral Ecology 24, 1048–1057 (2013).

21. S. M. Doucet, M. G. Meadows, Iridescence: a functional perspective. J. R. Soc. Interface 6 (2009).

22. B. E. Droguet, et al., Large-scale fabrication of structurally coloured cellulose nanocrystal films and effect pigments. Nat. Mater. 21, 352–358 (2022).

23. N. Vogel, et al., Color from hierarchy: Diverse optical properties of micron-sized spherical colloidal assemblies. Proc. Natl. Acad. Sci. U. S. A. 112, 10845–10850 (2015).

24. V. P. Patil, J. D. Sandt, M. Kolle, J. Dunkel, Topological mechanics of knots and tangles. Science (1979). 367, 71–75 (2020).

25. K. K. Nordén, C. M. Eliason, M. C. Stoddard, Evolution of brilliant iridescent feather nanostructures. Elife 10 (2021).

26. R. Maia, R. H. F. Macedo, M. D. Shawkey, Nanostructural self-assembly of iridescent feather barbules through depletion attraction of melanosomes during keratinization. J. R. Soc. Interface 9, 734–743 (2012).

27. H. Durrer, Schillerfarben der Vogelfeder als Evolutionsproblem. Denkschr. Schweiz Naturforsch. Ges. 14, 1– 126 (1977).

28. C. M. Eliason, R. Maia, J. L. Parra, M. D. Shawkey, Signal evolution and morphological complexity in hummingbirds (Aves: Trochilidae). Evolution (N. Y). 74, 447–458 (2020).

29. J. Zi, et al., Coloration strategies in peacock feathers. Proc. Natl. Acad. Sci. U. S. A. 100, 12576–12578 (2003).

30. P. Freyer, D. G. Stavenga, Biophotonics of diversely coloured peacock tail feathers. Faraday Discuss. 223, 49– 62 (2020).

31. H. Durrer, Bau und Bildung der Augfeder des Pfaus (*Pavo cristatus* L.). Revue suisse de zoologie. (1965). 10.5962/bhl.part.75643.

32. D.-J. Jeon, S. Paik, S. Ji, J.-S. Yeo, Melanin-based structural coloration of birds and its biomimetic applications. Appl. Microsc. 51, 14 (2021).

33. T. Okazaki, Reflectance spectra of peacock feathers and the turning angles of melanin rods in barbules. Zoolog. Sci. 35, 86–91 (2018).

34. R. Maia, L. D’Alba, M. D. Shawkey, What makes a feather shine? A nanostructural basis for glossy black colours in feathers. Proceedings of the Royal Society B: Biological Sciences 278, 1973–1980 (2011).

35. D. R. Rubenstein, et al., Feather gene expression elucidates the developmental basis of plumage iridescence in African starlings. Journal of Heredity 112, 417–429 (2021).

36. G. Gao, et al., Comparative genomics and transcriptomics of Chrysolophus provide insights into the evolution of complex plumage colouration. Gigascience 7, giy113 (2018).

37. H. Dürrer, Schillerfarben beim Pfau (*Pavo Cristatus* L.). Naturf. Ges. Basel 73 (1962).

38. R. Sarangarajan, R. E. Boissy, *Tyrp1* and Oculocutaneous Albinism Type 3. Pigment Cell Res. 14, 437–444 (2001).

39. J. Li, et al., A missense mutation in *TYRP1* causes the chocolate plumage color in chicken and alters melanosome structure. Pigment Cell Melanoma Res. 32, 381–390 (2019).

40. C. R. Goding, H. Arnheiter, MITF—the first 25 years. Genes Dev. 33, 983–1007 (2019).

41. H. van Grouw, W. Dekkers, The taxonomic history of Black-shouldered Peafowl; with Darwin’s help downgraded from species to variation. Bull. Br. Ornithol. Club 143 (2023).

42. N. I. Mundy, A window on the genetics of evolution: *MC1R* and plumage colouration in birds. Proceedings of the Royal Society B: Biological Sciences 272, 1633–1640 (2005).

43. Adelmann, C. H. et al. *MFSD12* mediates the import of cysteine into melanosomes and lysosomes. Nature. 588, 699–704 (2020).

44. Ducrest, A. L. et al. Melanin and neurotransmitter signalling genes are differentially co-expressed in growing feathers of white and rufous barn owls. Pigment Cell Melanoma Res. 38, (2025).

45. H. E. Hoekstra, Genetics, development and evolution of adaptive pigmentation in vertebrates. Heredity (Edinb). 97, 222–234 (2006).

46. Guernsey, M. W., Ritscher, L., Miller, M. A., Smith, D. A. & Schöneberg, T. A Val85Met mutation in Melanocortin-1 Receptor is associated with reductions in eumelanic pigmentation and cell surface expression in domestic rock pigeons (*Columba livia*). PLoS One 8, 74475 (2013).

47. MacDougall-Shackleton, E. A., Blanchard, L. & Gibbs, H. L. Unmelanized plumage patterns in Old World leaf warblers do not correspond to sequence variation at the melanocortin-1 receptor locus (*MC1R*). Mol. Biol. Evol. 20, 1675–1681 (2003).

48. O. Vidal, J. Viñas, C. Pla, Variability of the melanocortin 1 receptor (*MC1R*) gene explains the segregation of the bronze locus in turkey (*Meleagris gallopavo*). Poult. Sci. 89, 1599–1602 (2010).

49. D. Schwochow, et al., The feather pattern *autosomal barring* in chicken is strongly associated with segregation at the *MC1R* locus. Pigment Cell Melanoma Res. 34, 1015–1028 (2021).

50. S. Krishnan, R. L. Cryberg, Effects of mutations in pigeon *Mc1r* implicate an expanded plumage color patterning regulatory network. [Preprint] (2019). 10.1101/792945.

51. T. Hirobe, Role of dermal factors involved in regulating the melanin and melanogenesis of mammalian melanocytes in normal and abnormal skin. Int. J. Mol. Sci. 25, 4560 (2024).

52. L. Weiner, W. Fu, W. J. Chirico, J. L. Brissette, Skin as a living coloring book: how epithelial cells create patterns of pigmentation. Pigment Cell Melanoma Res. 27, 1014–1031 (2014).

53. M. Giraldo, J. Sosa, D. Stavenga, Feather iridescence of *Coeligena* hummingbird species varies due to differently organized barbs and barbules. Biol. Lett. 17, 20210190 (2021).

54. Y. Liu, et al., Comparison of structural and chemical properties of black and red human hair melanosomes. Photochem. Photobiol. 81, 135–144 (2005).

55. J. A. Brumbaugh, Ultrastructural differences between forming eumelanin and pheomelanin as revealed by the pink-eye mutation in the fowl. Dev. Biol. 18, 375–390 (1968).

