## Supporting information for "Modulation of Avian Iridescence via Melanogenesis"

#### Method Details

##### Ethics statement

All experimental procedures involving animals were carried out in compliance with Directive 2010/63/EU on the protection of animals used for scientific purposes. These procedures were reviewed and approved by the Animal Welfare and Ethics Committee of CIBIO/BIOPOLIS (approval reference: 2018_01 and 2019_03).

##### Genome sequencing of the Indian peafowl

*Sampling and DNA isolation.* A draft reference genome sequence of the Indian peafowl (*Pavo cristatus*) was generated from a phenotypically wild-type male. Whole blood was collected using a sterile needle and transferred into a heparin-free capillary. The sample was snap-frozen in liquid nitrogen and stored at -80°C until DNA extraction. High-molecular-weight genomic DNA was isolated from 5 µl of blood using a modified salt-based protocol (1), followed by a purification step using AMPure XP magnetic beads (Beckman Coulter) at a 3× beads-to-sample ratio. DNA quantity, purity, and integrity were assessed with a NanoDrop spectrophotometer, a Qubit dsDNA BR Assay Kit (ThermoFisher Scientific), and Agilent Genomic DNA ScreenTape (Agilent).

*Nanopore long-read sequencing.* Sequencing libraries were prepared using the Oxford Nanopore Technologies (ONT) 1D Sequencing Kit (SQK-LSK109). Briefly, 1.0 µg of high-molecular-weight genomic DNA underwent DNA repair and end-repair using the NEBNext FFPE DNA Repair Mix (NEB) and the NEBNext UltraII End Repair/dA-Tailing Module (NEB). Sequencing adaptors were ligated, and the libraries were purified using 0.4× AMPure XP Beads (Beckman Coulter) and eluted in the provided buffer (SQK-LSK109). Sequencing was performed on a GridION Mk1 (ONT) using R9.4.1 FLO-MIN106D flow cells, with data collected over 72 hours. Run quality parameters were monitored in real time using MinKNOW (v4.2.5).

Data from eight flow cells were base-called using *Guppy* (v3.2.6+afc8e14 for seven flow cells; v3.2.10+aabd4ec for one flow cell). Reads were filtered to remove those with: 1) an average Phred quality score ≤7; 2) lengths <1 kb; 3) low complexity sequences exceeding 40%; or 4) matches to the control sequence (lambda phage, 3.5 kb). Additionally, contamination was assessed using *Kraken* (v0.10.5-beta) (2), and reads matching *Spirometra erinaceieuropaei*, Human gammaherpesvirus-4, cloning vector pCA-DEST2303, or *Mycoplasma hyorhinis* (the four contributing to ~0.01% of reads) were excluded. After filtering, 4,599,390 ONT reads totaling 43.2 Gb (33× coverage) with an N50 of 16.6 kb were retained for downstream analysis.

*Linked-read sequencing.* The linked-read library was prepared using a Chromium Controller (10x Genomics) and the Genome Reagent Kits v2 (10x Genomics), following the manufacturer's protocol. In brief, 10 ng of high-molecular-weight genomic DNA was partitioned in GEM (Gel Bead-in-Emulsion) reactions containing unique barcodes (Gemcode) and loaded onto a Chromium controller chip. The droplets were recovered, isothermally incubated, and fractured. The intermediate DNA library was then purified and size-selected using Silane and Solid Phase reverse immobilization (SPRI) beads. An Illumina-compatible paired-end sequencing library was prepared following 10x Genomics recommendations and validated using an Agilent 2100 BioAnalyzer with the DNA 7500 assay (Agilent). Sequencing was performed on an Illumina NovaSeq 6000 platform using paired-end reads (2 × 151 bp).

The resulting reads were preprocessed for scaffolding using *LongRanger Basic* (v2.2.2), which performed basic read and barcode processing, including read trimming (removing the first 16 base pairs of read one if they match a valid barcode), barcode error correction, barcode whitelisting, and attaching barcodes to read IDs (producing barcoded reads). Additionally, the original linked read data was processed for use as a standard paired-end library during assembly.

All reads were filtered for contaminants by mapping with *gem-mapper* (3) (allowing up to 2% mismatches) against a contamination database that included phiX, UniVec sequences, *E. coli*, the complete mitochondrial sequence of Indian peafowl (NC_024533.1), and the four contaminants detected with *Kraken* (as described above). A total of 262,153,402 paired reads, corresponding to 78.6 Gb (~60× coverage), successfully passed the quality filters and were used for genome assembly.

*Chromatin conformation capture sequencing (Hi-C)*. A Hi-C library was prepared using the Arima High Coverage Hi-C kit (Arima Genomics), following the manufacturer's instructions. Sequencing was performed on an Illumina HiSeq 4000 platform using paired-end reads (2 × 151 bp). A total of 317,826,943 pairs of reads were aligned against the draft assembly using *Juicer* (v1.6) (4). This process yielded 261,213,953 Hi-C contacts, which were used for subsequent scaffolding.

*Contig construction and scaffolding.* The Indian peafowl reference genome sequence was assembled using a combination of data sources and multiple iterative steps.

1. Hybrid assembly:
   A hybrid genome assembly was generated with *MaSuRCA* (v3.4.1) (5, 6), using the Illumina linked-read data (60× coverage, treated as paired-end reads) and the pre-processed Nanopore reads (33× coverage) to create mega-reads. These mega-reads were assembled with *Flye* (v2.5) (7). Default parameters were used in *MaSuRCA*, with adjustments to the following settings: USE_LINKING_MATES = 0, MEGA_READS_PASS=0, JF_SIZE = 26200000000, and FLYE_ASSEMBLY=1.

2. Correction and scaffolding with linked reads:
   Barcoded 10x linked reads were used to correct and scaffold the assembly. Misassemblies were identified and resolved with *Tigmint* (v1.1.2) (8), followed by scaffolding with *ARKS* (v1.0.3) (9) and *LINKS* (v1.8.5) (10), as per a published pipeline (http://protocols.faircloth-lab.org/en/latest/protocols-computer/assembly/assembly-scaffolding-with-arks-and-links.html).

3. Hi-C scaffolding:
   The genome was further scaffolded using the Hi-C data with *Juicer* (v1.6) and the *3D de novo assembly (3D-DNA)* pipeline (v180114) (11).

*Curation of the chromosome-level assembly.* The *JuiceBox Assembly Tools* (*JBAT*) (v1.11.08) (12) were used for visual inspection and revision of the contact map. The putative misassemblies were revised and corrected. The decisions were made by taking into account breaks in synteny against the chicken chromosomes (GRCg6a) based on a pairwise alignment obtained with *nucmer* (13) and inspected with *Dot* (https://github.com/dnanexus/dot). Small scaffolds potentially containing inversions or translocations were examined within the super-scaffolds. Inversions or translocations detected in the dot plot were manually corrected using JBAT, ensuring changes maintained or improved consistency with the Hi-C contact map. Despite the ~35 million years of divergence between chickens and Indian peafowl, broad synteny across the majority of the genome is expected. However, intra-scaffold rearrangements consistent with the Hi-C contact map were left unchanged, as they could represent real species-specific differences. After completing the review, the assembly was finalized by exporting the modified assembly files and generating new FASTA files using the *run-asm-pipeline-post-review.sh* script from the *3D-DNA* pipeline. Finally, this revised assembly was sealed and closed by placing the debris scaffolds into gaps by running *run-asm-pipeline-post-review.sh* with options -s seal and -i 15000. The completeness of the final genome assembly was assessed using *BUSCO* (v5.8.2) (14) and the single-copy avian ortholog genes in the *aves_odb10* database. Summary statistics for the final assembly are provided in **table S1** and **figs. S2 and S3**.

### Genome annotation of the Indian peafowl

*RNA sequencing.* RNA-seq data from five tissues of the same male individual used for the reference genome sequence were generated to support gene annotation. Total RNA was extracted from brain, lung, testes, skin, and kidney tissues and quantified using the Qubit RNA BR Assay Kit (Thermo Fisher Scientific). RNA integrity was assessed with the Agilent RNA 6000 Pico Kit on a Bioanalyzer 2100 (Agilent). RNA-seq libraries were prepared using the KAPA Stranded mRNA-Seq Kit for Illumina platforms (Roche), following the manufacturer's protocol with 500 ng of total RNA as input. Library size and quality were evaluated using the High Sensitivity DNA Bioanalyzer Assay (Agilent). The RNA-seq libraries were run on an Illumina HiSeq 4000 platform using paired-end reads (2 × 101 bp).

*Repeat identification.* Repeats in the reference genome assembly were annotated using *RepeatMasker* (v4.0.7) (http://www.repeatmasker.org) with a custom repeat library available for chicken.

*Gene model inference.* Gene annotation was performed by integrating transcript alignments, protein alignments, and *ab initio* gene predictions.

1. Transcript alignments based on RNA-seq data:
   The multi-tissue RNA-seq reads were aligned to the genome using *STAR* (v.2.7.2a) (15) and transcript models were built for each tissue separately with *Stringtie* (v2.0.1) (16). These models were combined using *TACO* (17), and curated junctions to be used during the annotation process were identified with *Portcullis* (18) after mapping with *STAR*. *PASA* (v2.3.3) (19) was used to assemble transcripts, incorporating 2,104 Indian peafowl nucleotide sequences retrieved from NCBI (July 2020). Coding regions in the transcripts were identified with *TransDecoder program* (https://github.com/TransDecoder/TransDecoder), part of the *PASA* package.

2. Protein alignments:
   Complete chicken, human, and turkey proteomes were downloaded from UniProt (July 2020) and aligned to the genome using *spaln* (v2.4.7) (20).

3. *Ab initio* predictions:
   *Ab initio* gene predictions were performed on the repeat masked assembly with three different methods: *GeneID* (v1.4), Augustus (v3.2.3) (21), and *Genemark-ES* (v2.3e) (22) with and without RNA-seq junction evidence. The gene predictors were run with trained parameters for the human genome except for *Genemark* that runs in a self-trained manner.

4. Consensus gene models:
   *EvidenceModeler* (EVM; v1.1.1) (23) integrated all transcript, protein, and *ab initio* data to produce consensus coding DNA sequence (CDS) models. Untranslated regions (UTRs) and alternative splicing forms were annotated through two rounds of *PASA* updates.

5. Functional annotation:

Functional annotation of the proteins was conducted using *Blast2GO* (24). First, a *BLASTp* (25) search was performed against the *nr* database (accessed February 2021). Second, protein domains were identified with *InterProScan* (26). The results were combined in *Blast2GO* to generate the final functional annotations.

*ncRNA annotation.* Non-coding RNAs (ncRNAs) were annotated through the following steps. First, the *cmsearch* tool (v1.1; from *Infernal*) (28, 29) was used to scan the genome against the RFAM RNA family database (v12.0) (27). Transfer RNAs (tRNAs) were identified with *tRNAscan-SE* (v1.23) (30). To detect lncRNAs, *PASA* assemblies not annotated as protein-coding genes were screened to identify expressed, non-translated sequences. Assemblies longer than 200 bp and not covered >80% by small ncRNAs were classified as lncRNAs. lncRNA transcripts were clustered into genes based on shared splice sites or significant sequence overlap. Summary statistics of the genome annotation are given in **table S1**.

### Whole-genome resequencing and genetic mapping

*Sampling of peafowl color varieties and DNA isolation.* Blood samples or growing feathers were collected from 352 captive-raised Indian peafowl for whole-genome sequencing (**table S7**). Each individual was scored for the body color and wing phenotype, since they segregate independently. Color phenotypes were scored based on visual inspection, sometimes supplemented by genotype information provided by breeders for individuals known to be heterozygous for specific recessive mutations based on crossing records and studbook data. Samples included wild-type individuals and 20 distinct color varieties. Nine of these varieties are known to result from Mendelian mutations and therefore are determined by a single locus in the genome: Bronze, Cameo, Charcoal, Midnight, Opal, Purple, Ultramarine, Violet, and White/Pied. From segregation analyses in crosses, three of these varieties are known to be sex-linked and located on the Z-chromosome (Cameo, Purple, and Violet). For the remaining varieties (Brown, Chestnut, Hazel, Indigo, Ivory, Jade, Peach, Prussian-blue, Raw-umber, Sonja Violet, and Taupe), the inheritance patterns are either unknown, poorly supported, or attributed to combinations of two or more mutations. To investigate and take into account potential introgression from green peafowl (*Pavo muticus*) that might interfere with genetic mapping, four samples from green peafowl and 12 Spalding individuals, hybrids between Indian and green peafowl, were also collected.

Whole blood was collected as described earlier, but stored in 96% ethanol until DNA extraction. For birds where blood collection was not feasible, naturally molting feathers were plucked and similarly stored in ethanol. Genomic DNA was extracted from blood samples using a modified salt-based protocol, and from feathers it was extracted using the QIAamp UCP DNA Micro Kit (Qiagen). Potential RNA contamination was removed by treating DNA extracts with RNase-A (Roche). Following DNA isolation, DNA quality and purity were assessed using spectrophotometry (Nanodrop) and fluorometric quantitation (Qubit dsDNA BR Assay Kit, ThermoFisher Scientific).

*Illumina sequencing and read mapping.* Genome-wide polymorphism data for genetic mapping was generated using whole-genome Illumina sequencing. Sequencing libraries were prepared for each of the 368 individuals (Indian peafowl, green peafowl, and their hybrids). For blood samples, which have higher DNA quality and larger quantities, libraries were constructed using the TruSeq DNA PCR-Free Library Preparation Kit (Illumina). For some feather follicle samples, which often yield lower DNA quality or quantity, libraries were prepared using the PCR-based Nextera XT Library Preparation Kit (Illumina). The quality of each library was assessed based on fragment size distribution, evaluated on an Agilent 2200 TapeStation using an HS D5000 Screen Tape (Agilent Technologies), and molarity, calculated using a KAPA qPCR library quantification kit (Roche). Libraries were sequenced using paired-end reads (2 × 150 bp) on an Illumina instrument.

Prior to data analyses, sequencing read quality was inspected with *FastQC* (v0.11.8) (https://www.bioinformatics.babraham.ac.uk/projects/fastqc/). Reads were then aligned to the Indian peafowl genome assembly generated in this study using *BWA-MEM* (v0.7.17-r1188) (31) with default parameters. Duplicate reads were flagged for downstream analysis using the *MarkDuplicates* function in *Picard* (v3.0.0) (http://broadinstitute.github.io/picard). Read group information was added to each BAM file using the *Picard* function *AddOrReplaceReadGroups*. Sequencing and mapping summary statistics were calculated using *SAMtools* (v1.11) (32). After mapping and duplicate removal, the final effective mean sequencing depth was 10.1× ± 5.3 (**table S7**).

*Variant discovery and genotype calling.* Single nucleotide polymorphisms (SNPs) and small insertions or deletions (indels) were identified using *GATK* (v4.2.6.1) (33). Briefly, *gVCF* files were generated for each individual using the *HaplotypeCaller* function requiring a minimum mapping quality of 30 and setting the heterozygosity value to 0.004. These files were subsequently combined into a single file using the function *CombineGVCFs*, and variants were called using the function *genotypeGVCFs* using default parameters except for the heterozygosity value (--heterozygosity 0.004). Prior to downstream analyses, additional filtering steps were applied at both the genotype and variant levels using *VCFtools* (v0.1.16) (34). The genotype of each individual was coded as 'missing data' if its quality was below 30 (--minGQ 30), or if its coverage was below 4× or higher than 58× (i.e., twice the average coverage of the individual with the highest coverage; --minDP 4 --maxDP 58). Following genotype filtering steps, variants with 50% or more missing genotype data across individuals (--max-missing 0.5 in *VCFtools*) were excluded.

*Variant effect prediction.* Filtered variants were annotated for their potential impact on protein-coding regions, including nonsynonymous, frameshift, splicing, and stop-gain mutations, using the *SnpEff* toolbox (v4.3t) (35). Based on these annotations, a subset of variants classified as high or moderate impact mutations was extracted for independent analysis of coding region variation. This filtering process resulted in a dataset of 228,553 variants.

*Genome-wide association analysis.* Before conducting the genome-wide association analysis, missing genotypes were imputed, and genotype data were phased using the software *Beagle* (v5.1) (36, 37). Association testing for each marker was performed with *GEMMA* (v0.98.1) (38) using a linear mixed model and coding each color morph as a binary trait (mutant vs. all others). The varieties that were known or discovered to be combinations of mutations were sometimes excluded from the control or case groups. Individuals belonging to the *white* variety ($n$ = 17) were excluded from the mapping analyses of the other varieties because the mutation is epistatic, masking all other phenotypes. To minimize the potential confounding effects of population stratification and relatedness among individuals, a kinship matrix was included in the model as a random effect. The kinship matrix was estimated using *GEMMA*'s centered relatedness matrix option (-gk 1). Additionally, the first three principal components (PCs) from a principal components analysis (PCA), calculated with *PLINK* (v1.90b6.26) (39), were included as covariates. Variants with a minor allele frequency (MAF) lower than 10% were not considered. Significance thresholds for association testing were determined using a Bonferroni correction (-$\log_{10}$ [0.05/ number of loci]) applied to the likelihood ratio test for allele frequencies from *GEMMA*. Manhattan plots summarizing the associations were generated with the *qqman R* package (40).

### Bulk RNA-sequencing

*Sampling and RNA extraction.* RNA-seq data were generated from 37 regenerating feather follicle samples of Indian peacocks, including neck feather follicles from wild-type and mutant varieties, and wing feather follicles from wild-type and Black-shoulder individuals (**table S8**). Feather follicles were collected in the pin stage of development (an early stage of feather development shortly after emerging through the skin), placed in RNAlater at room temperature for 8 hours, and subsequently stored at -80°C until RNA extraction. Total RNA was isolated using the RNeasy Plus Mini Kit (Qiagen). Following extraction, RNA integrity was assessed using a TapeStation RNA ScreenTape (Agilent), and RNA concentration was measured using a Qubit RNA BR assay kit (ThermoFisher Scientific).

*Library preparation and sequencing*. Illumina strand-specific RNA-seq libraries were prepared using the TruSeq Stranded mRNA Kit, according to the manufacturer's instructions. Sequencing of these libraries generated approximately 1,085 million paired-end reads (2 × 151 bp), with an average of ~70 million reads per individual (range: 41,911,332-97,612,226) (**table S8**).

*Differential gene expression analysis*. RNA-seq reads were aligned to the Indian peafowl reference genome using *HISAT2* (v.2.2.1) (41). Gene expression analyses were carried out using the *R* package *DESeq2* (v1.36.0) (42), with count data for each transcript generated with the *featureCounts* (43) function of the *Rsubread* package (v1.22.2) (44) against the annotation file. Differential expression analyses compared biological replicates of each mutant to wild-type birds (neck and wing analyzed separately). Transcripts with adjusted $P$-values (corrected for multiple testing) below 0.1 were considered significantly differentially expressed.

*Allelic imbalance analyses*. Allelic expression imbalance in neck and wing feather follicles was quantified in individuals heterozygous for the Black-shoulder nonsynonymous *MC1R* mutation (see main manuscript). For each sample, we estimated the relative expression of the two alleles either by directly counting RNA-seq reads overlapping the mutation site (when coverage exceeded 50×) or by targeted amplicon sequencing when RNA-seq coverage was below that threshold. Amplicon library preparation for Illumina sequencing followed Andrade *et al.* (2019) (45), with the following changes: first PCR with 5'-tailed primers (insert size: 177 bp; primer forward: 5' – TCGTCGGCAGCGTCAGATGTGTATAAGAGACAGCAATGAGCTCTTCCTGACG – 3'; primer reverse: 5' – GTCTCGTGGGCTCGGAGATGTGTATAAGAGACAGCAGCAGCATGAAGAGCATC – 3'), a touchdown PCR from 66-62°C, and PCR clean-up with 1:1 bead-to-sample volume ratio. The resulting libraries were pooled and sequenced using paired-end Illumina reads (2 × 150 bp). Reads were mapped to the reference genome using *BWA-MEM* (v0.7.17-r1188) with default parameters, and counts for the alleles were obtained by visual inspection with the *Integrative Genomics Viewer* (*IGV*) (46). For the neck, three heterozygous samples were analyzed ($n$ = 3): two using RNA-seq and one using amplicon sequencing. For the wing, allelic proportions were obtained from two samples ($n$ = 2) using amplicon sequencing only, as the RNA-seq datasets for wing follicles had <50× coverage at the mutation site.

**Macroscopic photographs of feathers**

The macroscopic photographs of the feathers from (i) the train eyespot region of wild-type and mutant peafowl varieties, (ii) the covert wing feathers of the common emerald dove (*Chalcophaps indica*), and (iii) the secondary wing feathers (or speculum) from the mallard duck (*Anas platyrhynchos*) were taken with a Sony Alpha 6400 combined with a Godox V1s flash. Both were installed in a setup with an angle of the flash of 13°.

**Optical microscope characterization of the peafowl eyespot**

Light microscope images and microscopic reflectance spectra were acquired with a correlative microscope setup consisting of a Zeiss Axio Imager Z2 optical microscope (Zeiss) combined with an MCS CCD UV-NIR spectrometer (Zeiss). The microscope was equipped with an

Axiocam 820 color camera (Zeiss). Illumination was provided by a built-in halogen lamp (Osram) at 9.76 V. Images of individual barbules were acquired using a 20× objective with a numerical aperture (NA) of 0.5. Spectral measurements were performed using a 50× objective with an NA of 0.55. A silver mirror (Thorlabs) served as a 100% reflectance reference. The measured spot size was 16×16 µm. A single spectrum was recorded per spot. The integration time was adjusted to avoid detector saturation and typically ranged between 100-200 ms for most color types, and 300-400 ms for *ultramarine* samples. Internal spectrometer smoothing was set to 5, and an averaging of 10 spectra was applied by the software. Binning was set to 2 × 2. Multiple spectra of the individual barbules were recorded, and the mean and standard deviation were calculated. For sample preparation, the individual barbules were carefully cut from the barb and adhered to a carbon pad (Plano).

**Color measurement and chroma determination of the peafowl eyespot**

Perceived color was quantified in the CIELAB (L*, a*, b*) color space, as defined by the International Commission on Illumination (CIE). CIELAB coordinates were calculated from the reflectance spectra recorded with the Zeiss Axio Imager Z2 microscope-spectrometer setup described above. The transformation followed standard procedures under the CIE 2006 convention, where L* represents lightness, a* corresponds to the green–red axis, and b* corresponds to the blue–yellow axis, under standard daylight illumination (D65) with a 2° observer angle and a step size of 5 nm. Chroma values were subsequently derived from these coordinates (47).

**Spectroscopy of the peafowl eyespot**

For the angle-dependent spectrometry, a customized setup was employed. A tungsten halogen lamp (HL-2000-HP-FHSA, Ocean Insight) served as the light source. The collected light was analyzed using a Flame-T-VIS-NIR-ES spectrometer (Ocean Insight). Both components were connected to collimators (74-UV-MP, Ocean Insight), which were mounted on mechanical arms and linked via glass fiber cables (105 µm, 0.22 NA Fiber Patch Cable, Thorlabs). These arms were controlled using two motorized rotational stages (Standa) to precisely adjust the angles of incidence and detection. Spectra were recorded using a custom *LabView*-based acquisition software. With this software, the integration time, averaging, and data export were controlled.

For sample preparation, the feathers were glued to a glass slide using carbon pads (Plano), ensuring a flat and stable surface for accurate measurements. Specular reflection measurements were conducted. The incidence angle was varied from 60° to 3° and subsequently from −3° to −60° in 1.5° steps. Measurements for incidence angles between 3° and −3° were not possible due to the obstruction of the light signal by the detector arm. Because angle-dependent spectra recorded from opposite sides exhibited symmetric (mirrored) peak behavior, only one side of the measurements was used for subsequent analysis (**fig. S17**). For reference measurements, a white standard (Labsphere) was used as the maximum reference, measured at the same angles as the sample. The minimum reference was obtained by recording the spectrometer signal without illumination. An integration time of 2800 ms and an averaging of 10 scans were used. For samples with a well-ordered photonic crystal (wild-type, Purple, Ultramarine, and Violet feathers), the spectra were smoothed over 200 points using the Adjacent-Averaging method in *OriginPro* (v2023b) (OriginLab), and the peak position was determined based on the highest intensity. For samples exhibiting either a less well-ordered photonic crystal or a highly ordered but few-layer photonic crystal structure that resulted in low reflectance (e.g., Bronze and Opal feathers), the spectra were first smoothed and subsequently baseline-corrected using a spline function in *OriginPro*, as the broadband pigment background otherwise obscured the structural peak. Peak positions were then determined by applying a Gaussian fit to the peak region, also in *OriginPro*. For samples without a photonic crystal (Cameo and Charcoal feathers), the spectra were only smoothed (200-point), and no peak could be evaluated. Because transformations did not significantly impact the peak wavelength identified for mutants across the multiangle measurements, values of the 200-point measures were used for peak selection (**fig. S18**).

### Transmission Electron Microscopy (TEM)

*TEM initial sample preparation.* Feathers used for TEM were collected from the same sources described for macroscopic photographs, including wild-type and mutant peafowl, the common emerald dove, and the mallard duck. Feathers were washed twice with 0.1 M phosphate buffer (PB, pH 7.4) containing 0.5% Tween 20 (Carl Roth) and stored overnight at 4°C in the same solution. Samples were then treated for 2 hours at room temperature with a solution of 1.5% potassium ferricyanide (Merck) and 1% osmium tetroxide (>99.95%, Emsdiasum) in 0.1 M PB (pH 7.4). Afterwards, the feathers were washed three times in the same PB-Tween solution and stored overnight at 4°C.

*Preparation of 0.1 M Phosphate Buffer Solution (Sørensen method).* To prepare 1 liter of phosphate buffer solution (PB), 14.417 g of disodium hydrogen phosphate dihydrate ($Na_2HPO_4 \cdot 2H_2O$, ≥99%, Carl Roth) and 2.622 g of sodium dihydrogen phosphate monohydrate ($NaH_2PO_4 \cdot 1H_2O$, ≥98%, Carl Roth) were dissolved in 900 ml of Milli-Q water. The pH of the solution was adjusted to 7.4 using hydrochloric acid (HCl, ≥25%, Carl Roth) or sodium hydroxide (NaOH, ≥98%, Carl Roth). Finally, Milli-Q water was added to bring the total volume to 1000 ml.

*Feather dehydration process*. Feathers were progressively dehydrated at room temperature through a graded ethanol–acetone series. Samples were first immersed in 70% ethanol (≥99.8%, denatured, Carl Roth) for 60 minutes, followed by 80%, 90%, and 100% ethanol (denatured) for 30 minutes each. The dehydration continued with 100% ethanol (undenatured, ≥99.8%, Carl Roth) for 30 minutes, then a 50/50 mixture of ethanol (≥99.8%, Carl Roth) and acetone (≥99.8%, Carl Roth) for 30 minutes, followed by 100% acetone for 30 minutes. Finally, samples were immersed in 2/3 acetone and 1/3 Epon resin, and then 1/3 acetone and 2/3 Epon resin, each for 30 minutes at room temperature.

*Preparation of Epon resin.* Epon resin was prepared by mixing Solution A and Solution B as follows:

1) Solution A: 75 mL Glycidether 100 (Carl Roth) and 120 mL Glycidether hardener DBA (Carl Roth)
2) Solution B: 120 mL Glycidether 100 (Carl Roth) and 105 mL Glycidether hardener MNA (Carl Roth)

The two solutions were combined and supplemented with 2% accelerator DMP-30 (Carl Roth). The mixture was gently stirred until homogeneous and stored at 4°C until use.

*Embedding and sectioning.* Dehydrated feathers were embedded in Epon resin with glycidether accelerator DMP-30 (Carl Roth) overnight at 4°C. Polymerization was carried out by incubating samples at 60°C overnight, followed by 80°C, again overnight. The Epon blocks were cut into 40-60 nm thick sections using a Leica Ultracut UCT. These sections were treated with chloroform (≥99%, Carl Roth) to ensure a flat morphology and transferred onto copper grids coated with Pioloform FN65 (Plano) film.

*Section preparation and image acquisition.* Sections were stained with 10% uranyl acetate (>98%, Serva) for 10 minutes, rinsed with Milli-Q water, treated with 2.8% lead citrate (Merck) for 10 minutes, and rinsed again with Milli-Q water. Imaging was performed using a Zeiss Leo 906 transmission electron microscope (Zeiss) operated in bright-field mode at 60 kV with a TRS 2048 digital camera system.

*Nanostructure measurements:* Several variables related to melanosome size and spatial distribution in the keratin matrix were measured from TEM images. These variables included: number of melanosome layers (*N*), distance between layers (*a*), distance between melanosomes within a layer (*b*), cortex thickness (*c*), diameter of melanosomes (*Dm*), diameter of air pockets (*Da*), and length of melanosomes (*Lm*). Measurements were manually taken using the *straight line* and *measure* tools from *ImageJ* (48) software on longitudinal and transversal cuts of feather barbules. For each breed–zone combination ($n$ = 42), 100 measurements were taken, and the mean and standard deviation were calculated.

*Statistical analyses.* Linear discriminant analyses (LDA) were conducted using either "phenotype" or "area" as the grouping variable. Model performance was assessed as the percentage of individuals correctly reclassified to their original group, based on a classifier that minimized each sample's multivariate Mahalanobis distance to the group mean. Pearson's correlation coefficients ($r$) were calculated to identify significant associations ($P < 0.05$) between Z-score standardized averages of the original measurements and the axes of the LDA models. Analyses were implemented using the *MASS R* package (49). To estimate the relative contribution of each variable to total dataset variance, squared loadings were weighted, summed, and normalized across LD axes to obtain contribution percentages.

**Histology of regenerating feather follicles**

Regenerating feather follicles at the pin stage of development were collected from the following sources: (i) the neck region of wild-type and mutant peafowl varieties, (ii) the covert wing feathers of the common emerald dove, and (iii) the secondary wing feathers (or speculum) from the mallard duck. The samples were fixed in 4% paraformaldehyde prepared in 0.1M PBS (pH 7.4) at 4°C for four days. After fixation, the follicles were washed in PBS and stored in 70% ethanol until further processing. The follicles were then dehydrated in an ascending ethanol series, cleared in xylene, and embedded in Paraplast Plus (Sigma). Longitudinal 10 µm thick sections were cut with a Reichert-Jung 2040 Autocut Rotary Microtome. The ribbon of sections was floated on the surface of distilled water at 37°C and mounted onto gelatin-coated glass slides. The sections were heated at 65°C for 1 hour to melt the paraffin, deparaffinized in xylene, and mounted with DPX mounting medium (Sigma). Images were captured using a Zeiss Axio Imager M2 epifluorescent microscope equipped with AxioCam MRc5 camera (Zeiss) and *AxioVision Se64* (vRel.4.9.1) software (Zeiss). A 63× Plan-apochromat oil immersion objective was used for imaging.

**Chemical characterization of melanin**

Fully grown feathers from the same sources used in the histological analysis (see previous section) were collected for the quantification of melanin content. Feathers were kept in black plastic zip-lock bags at room temperature, protected from direct light to prevent degradation. Approximately 15 mg of each feather sample was homogenized in water using a Ten-Broeck glass homogenizer at a concentration of 10 mg/ml. An aliquot of 100 µl (equivalent to 1 mg of sample) was mixed with 900 µL of Soluene-350 (Perkin-Elmer) in a 10 ml screw-capped conical test tube (50). The mixture was vortexed and heated at 100˚C in a boiling water bath for 15 minutes, then cooled. This process was repeated, followed by vortexing and centrifugation at 4,000 g for 3–5 minutes. The resulting supernatant was analyzed using a spectrophotometer at 500 nm (A500) and 650 nm (A650). A blank reference sample (100 µl water and 900 µl Soluene-350) was processed under the same conditions. To analyze pyrrole-2,3,5-tricarboxylic acid (PTCA), pyrrole-2,3-dicarboxylic acid (PDCA), and thiazole-2,4,5-tricarboxylic acid (TTCA), alkaline hydrogen peroxide oxidation (AHPO) was employed (51), while 4-amino-3-hydroxyphenylalanine (4-AHP) and 3-amino-4-hydroxyphenylalanine (3-AHP) were quantified via hydroiodic acid (HI) hydrolysis (52). Absorbances due to proteins (0.021 and 0.001 per mg of feather) were subtracted from A500 and A650 values. Total melanin, eumelanin, benzothiazine-pheomelanin (BT), and benzothiazole-pheomelanin (BZ) contents were calculated by multiplying the concentrations of A500, PTCA, 4-AHP, and TTCA by factors of 101 (49), 80 (51), 9 (53), and 34 (54), respectively. The quantification of different metabolites from these pigment analyses is available in **tables S6 and S9**.

**Single-cell RNA-sequencing**

*Tissue preparation and production of libraries for scRNA*-seq. The experiments were conducted on regenerating feathers of one male emerald dove and two male mallards, taking advantage of the fact that these species express within some feathers both iridescent and non-iridescent colors (**Fig. 6A; fig. S11**). For each species, two scRNA-seq libraries were generated per individual feather to capture the structural coloration asymmetry, with one library representing the left side and the other the right side (emerald dove: $n$ = 4 libraries; mallard: $n$ = 4 libraries). For the emerald dove, wing covert feathers belonging to a row of feathers displaying color asymmetry relative to the central axis of the feather itself (i.e., the rachis) were analyzed. The feathers were sampled approximately nine days after being plucked, at a time when color differences between the two sides had just started to be visible. For the mallard duck, feathers of the speculum, a bright iridescent patch of inner secondary flight feathers, were sampled. Mallard feathers were sampled 20 days after plucking during the formation of the iridescent patch.

For all the experiments, regenerating feathers were plucked, immediately placed on ice in Hanks' Balanced Salt Solution (HBSS, Gibco), and dissected under a stereomicroscope. The distal-most half of the feather, which is fully keratinized at the time of sampling, was removed (**fig. S11**). The remaining feather was incised longitudinally along the rachis and flattened by pinning its corners in HBSS. This exposed the inner pulp, which was then carefully removed with forceps. At this stage, in both emerald dove and mallard feathers, the color asymmetry is evident as a darkening of the tissue where structural coloration would emerge during later keratinocyte maturation. After confirming this difference, the feather was further cut at about 4 mm from its base, at a level where barb ridges are formed, and the proximal-most portion of the sample was kept. The left and right halves of the feather were then separated by cutting along an evident midline devoid of keratinocytes, and processed individually for cell dissociation.

The samples were dissociated in 2mg/ml dispase solution in DMEM (Gibco) for 60 min at 37°C at 300 rpm. After 10 minutes of incubation, 30 µl of Liberase solution (Roche) in DMEM (10 mg/ml) was added, and the mixture was passed through a pipette tip several times every 10 minutes. The partially dissociated tissues were pelleted by centrifugation at 400 × g at 4°C and incubated in 0.05% Trypsin/EDTA (Gibco) for 10 minutes at 37°C and 300 rpm mixing, and occasionally passed through a pipette tip. The digestion was arrested by adding 10% FBS (Gibco) in DMEM, the feather sheath was removed with forceps, and the mixture was treated with DNase I (Roche) at 37°C for 5 min and 300 rpm. The cells were then washed twice in 0.4% BSA (Sigma-Aldrich) in HBSS (with calcium and magnesium), filtered twice through a 40 µm-mesh filter (Falcon) and once through a Flowmi 40 µm Cell Strainer (Bel-Art SP Scienceware), resuspended in 0.4% BSA in HBSS, and counted in a hemocytometer after trypan blue staining. The resuspended cells were partitioned and barcoded separately using a 10x Genomics Chromium instrument (10x Genomics) following the manufacturer's protocol (Chromium Next GEM Single Cell 3′ Reagent Kits v3.1 Dual Index, Rev C) and targeting the recovery of 10,000 cells per reaction. Following quantification and quality control (LightCycle qPCR, Agilent TapeStation), the libraries were sequenced on an Illumina instrument and demultiplexed using the *CellRanger Fastq* pipeline (10x Genomics). Summary statistics for the single-cell libraries are given in **table S10**.

*Preprocessing of the scRNA-seq data*. The scRNA-seq libraries were pre-processed individually using the *count* pipeline in *Cell Ranger* (v7.0.1; 10x Genomics). Briefly, the reads were aligned to the reference genomes (emerald dove: this study and described below; mallard: GCF_047663525.1) using the splicing-aware aligner *STAR* (66) as implemented in *Cell Ranger*. Uniquely mapped reads within the start and end coordinates of each gene were considered for UMI counts, and cells were distinguished from empty droplets using an algorithm based on the *EmptyDrop* method (55). Filtered count matrices were imported into the *R* package *Seurat* (v5.3.0) (56) and further filtered to remove low-quality cells. For the emerald dove ($n$ = 4) and mallard ($n$ = 4) libraries, cells with UMI counts between 2,000 and 18,000 and between 2,000 and 20,000, respectively, and expressing more than 750 features were retained. After filtering, 10,213 emerald dove cells and 20,991 mallard cells remained. Gene expression matrix processing (data normalization and scaling) and dimensionality reduction by Principal Components Analysis (PCA) were performed independently for each library in *Seurat* with default settings. For each species separately, the libraries were integrated using the *HarmonyIntegration* method as implemented in the *IntegrateLayers* function in *Seurat*.

*Cluster annotation*. Unsupervised clustering of cells was performed on the integrated data using the *FindNeighbors* (*k.param* = 50, $dims_{\text{dove}}$ = 50; $dims_{\text{mallard}}$ = 30) and the *FindClusters* ($resolution_{\text{dove}}$ = 0.14; $resolution_{\text{mallard}}$ = 0.05) functions in *Seurat.* Differential gene expression analyses between each cluster and all remaining cells in the dataset, as well as between selected cluster pairs, were conducted with the Wilcoxon rank-sum test as implemented in the *FindAllMarkers* and *FindMarkers* functions in *Seurat,* respectively. Cluster annotation was manually curated based on the expression of known marker genes and the top differentially expressed genes in each cluster. An overview of the cluster annotations and representative marker genes is shown in **figs S12-S15**, and complete lists of the top differentially expressed genes for each cluster are provided in **Data S3** and S4.

Within each dataset, clusters of cells showing enriched expression of keratin genes (e.g., *Keratin-14*, *Keratin-19*, *Keratin-8*, *Keratin-5*) and known keratinocyte markers (e.g., *SHH* and *COL17A1*) (57-59) were collectively labeled as keratinocytes. In both datasets, two distinct clusters of cells showed enriched expression of melanocyte-specific markers such as *TYR*, *TYRP1*, *PMEL*, and *MLANA*. Within each dataset, pairwise differential expression analyses between these clusters identified one population of cells (labeled as melanocytes 1) showing upregulation of several melanocyte markers (e.g., *TRPM1*: $\text{Log}_2\text{FC}_{\text{dove}} = 4.6$; $P < 0.0001$; $\text{Log}_2\text{FC}_{\text{mallard}} = 3.7$, $P < 0.0001$). In both datasets, the second population (labeled as melanocytes 2) showed enriched expression of several glutathione S-transferases (e.g., Hematopoietic Prostaglandin D Synthase*, HPGDS*: $\text{Log}_2\text{FC}_{\text{dove}} = 1.76$; $P < 0.0001$; $\text{Log}_2\text{FC}_{\text{mallard}} = 1.29$, $P <$ 0.0001). A previous study in quails identified two distinct melanocyte lineages committed to the production of eumelanin or pheomelanin within embryonic skin tracts, eventually forming black or yellow feather stripes (60). These populations were negative and positive for the melanoblast/melanocyte early marker (MelEM) monoclonal antibody, which binds a glutathione S-transferase alpha class subunit and labels melanoblasts and early melanocytes in the avian embryo (61, 62). Increased glutathione S-transferase expression and decreased melanocyte-specific marker gene expression suggest that these cells are early differentiating melanocytes, possibly progressing towards pheomelanin production. Based on the common expression of many melanocyte markers, these populations of cells were collectively labeled as melanocytes and were considered together for some analyses.

Additional small but clearly distinct clusters were manually annotated. In both datasets, a small cluster of cells expressing erythrocyte markers, such as *HBAA* and *HBB*, was manually assigned and annotated as erythrocytes (emerald dove: $n$ = 30 cells; mallard: $n$ = 67 cells). In the mallard dataset, co-expression of erythrocyte markers and some keratin genes suggests the contamination of small numbers of low-quality keratinocyte cells within the erythrocyte cluster. In the emerald dove dataset, one small cluster of cells ($n$ = 29 cells) showed enriched expression of neural crest-derived cell genes (e.g. *FOXD3* and *NGFR*) and likely represents the melanoblast population at the base of the feather follicle. In the absence of canonical melanoblast markers (e.g., *DCT, EDNRB2*), this cluster was manually assigned and labeled as neural crest-derived cells. Overall, these changes represented minor manual adjustments to the clustering output, limited to clearly distinct and well-characterized cell populations, and did not affect the overall interpretation of the data. The remaining clusters in both datasets were left unchanged from the clustering provided by *Seurat* and were annotated based on the expression of marker genes for cell types previously described in regenerating feather follicles (7).

For the emerald dove, the final annotation of cells consisted in 11 clusters: (1) keratinocytes, $n$ = 4,261; (2) mesenchymal cells 1, $n$ = 1,716; (3) melanocytes 1, $n$ = 1,430; (4) endothelial cells, $n$ = 1,215; (5) melanocytes 2, $n$ = 566; (6) leukocytes 1, $n$ = 373; (7) mesenchymal cells 2, $n$ = 276; (8) leukocytes 2, $n$ = 239; (9) leukocytes 3, $n$ = 77; (10) erythrocytes, n = 31, and (11) neural crest-derived cells, $n$ = 29. For the mallard, the final annotation consisted of 8 clusters: (1) keratinocytes, $n$ = 14,312; (2) mesenchymal cells 1, $n$ = 4,376; (3) melanocytes 1, $n$ = 855; (4) leukocytes, $n$ = 491; (5) endothelial cells, $n$ = 406; (6) melanocytes 2, $n$ = 334; (7) mesenchymal cells 2, $n$ = 150; and (8) erythrocytes, $n$ = 67.

*Tissue composition analyses.* For both datasets, cell counts per cluster were extracted from each library ($n$ = 4 libraries per species) and normalized by library total cell counts. Cell-type proportions in the samples were compared between conditions (iridescent color vs. non-iridescent color) with a Welch Two Sample $t$-test in *R* ($n$ = 2 replicates per condition). Cell-type counts and summary test statistics are reported in **table S11**.

*Differential gene expression analyses.* For each species and for each cell type, differential expression analyses were performed using a pseudo-bulk approach. For all the analyses, differentially expressed genes with $Log_2FC \geq 0.5$ and $P$-value $\leq 0.05$ were considered. To test for differences between feather samples developing iridescent or non-iridescent colors (i.e., color types), summed UMI counts within each sample ($n_{iridescent}$ = 2 samples per dataset, $n_{non\text{-}iridescent}$ = 2 samples per dataset) were used as inputs for *DESeq2* (v1.48.1) with default parameters as implemented in the *FindMarkers* function in *Seurat*. Lists of differentially expressed genes per cell type for the emerald dove and the mallard are reported in **Data S1 and S2**, respectively.

*Gene ontology enrichment analyses*. For each species, gene ontology (GO) enrichment analyses were carried out for the lists of melanocyte-specific differentially expressed genes (considering the merged populations of melanocytes 1 and 2) using the *topGO* package (v.2.59.0) (https://bioconductor.org/packages/topGO) in *R*. For each dataset, enrichment was assessed by comparing the frequencies of GO term annotations in the differentially expressed gene sets against background GO term frequencies in the species' genomes, followed by statistical testing by Fisher's exact test. GO terms associated with the differentially expressed gene sets for the emerald dove and the mallard are reported in **Data S5 and S6**, respectively.

*Emerald dove genome assembly and annotation*. The individual used to produce the draft reference genome sequence of the emerald dove was one adult male obtained from a licensed breeder in Portugal. Whole blood was collected using a sterile needle into a heparin-free capillary, transferred into a 10ml Vacutainer K2 EDTA tube (BD Biosciences), and immediately snap-frozen in liquid nitrogen. The sample was subsequently shipped to the genomic services provider Macrogen (Seoul, South Korea) in dry ice. High molecular weight DNA was extracted from 140 µl of blood using the Wizard HMW DNA Extraction kit (Promega). DNA quantity and integrity were assessed using a Victor Nivo Multimode Microplate Reader (PerkinElmer) with the QuantiFluor® dsDNA System (Promega) method, the Femto Pulse System (Agilent), and a N120 NanoPhotometer (IMPLEN). The average fragment length of the extracted DNA was estimated to be approximately 44 kb.

A Pacific Biosciences (PacBio) Revio sequencing library was prepared from 7 μg of DNA input. gDNA was sheared with the Megaruptor® 3 (Diagenode) and both purified and size-selected using AMPure PB magnetic beads (Pacific Biosciences). The size distribution was determined using the Femto Pulse System (Agilent) for all size quality controls. The library insert size was within the optimal size range. A total of 10 μL of library was prepared using the PacBio SMRTbell prep kit 3.0 and SMRTbell templates were annealed using the Revio polymerase kit (Pacific Biosciences). Sequencing was performed using a Revio sequencing plate and Revio SMRT Cell tray. Each SMRT cell was captured using a 24-hour movie time using the PacBio Revio (Pacific Biosciences) sequencing platform by Macrogen (Seoul, South Korea). The subsequent steps are based on the PacBio Sample Net-Shared Protocol, available at https://www.pacb.com/.

A total of 6,372,772 HiFi long reads with an average length of 13,480 bp were generated, which corresponds to ~71× coverage of the approximate size of a bird genome (~1.2 Gb). Contig assembly was performed using *Hifiasm* (v0.20.0-r639) (63), resulting in 1,283,043,043 bp of primary assembly (contigs $n = 315$) with a contig N50 of 33 Mb and a maximum length of 121.1 Mb. These contigs were scaffolded into a pseudochromosome assembly using *RagTag* (v2.1.0) (64), with the rock pigeon *bColLiv1.pat.W.v2* genome assembly (GCF_036013475.1) serving as the reference. The final assembly comprised 219 scaffolds with an N50 of 83.8 Mb and a maximum scaffold length of 215.1 Mb. Completeness analysis of the assembly was performed using *BUSCO* (v5.8.2) and the 8,338 single-copy avian ortholog genes (*aves_odb10*), which indicates that our genome assembly was 97.1% complete. Annotation of the draft reference genome was conducted using *Liftoff* (v1.6.3) (65) by lifting over gene models from the annotation available for the rock pigeon genome. Genes of interest for the single-cell analyses were manually inspected.

### References


1. E. D. Enbody, *et al.*, A multispecies *BCO2* beak color polymorphism in the Darwin's finch radiation. *Current Biology* **31**, 5597-5604.e7 (2021).
2. D. E. Wood, S. L. Salzberg, Kraken: ultrafast metagenomic sequence classification using exact alignments. *Genome Biol.* **15**, R46 (2014).
3. S. Marco-Sola, M. Sammeth, R. Guigó, P. Ribeca, The GEM mapper: fast, accurate and versatile alignment by filtration. *Nat. Methods* **9**, 1185–1188 (2012).
4. N. C. Durand, *et al.*, Juicer provides a one-click system for analyzing loop-resolution Hi-C experiments. *Cell Syst.* **3**, 95–98 (2016).
5. A. V. Zimin, *et al.*, Hybrid assembly of the large and highly repetitive genome of *Aegilops tauschii* , a progenitor of bread wheat, with the MaSuRCA mega-reads algorithm. *Genome Res.* **27**, 787–792 (2017).
6. A. V. Zimin, *et al.*, The MaSuRCA genome assembler. *Bioinformatics* **29**, 2669–2677 (2013).
7. M. Kolmogorov, J. Yuan, Y. Lin, P. A. Pevzner, Assembly of long, error-prone reads using repeat graphs. *Nat. Biotechnol.* **37**, 540–546 (2019).
8. S. D. Jackman, *et al.*, Tigmint: correcting assembly errors using linked reads from large molecules. *BMC Bioinformatics* **19**, 393 (2018).
9. L. Coombe, *et al.*, ARKS: chromosome-scale scaffolding of human genome drafts with linked read kmers. *BMC Bioinformatics* **19**, 234 (2018).
10. R. L. Warren, *et al.*, LINKS: Scalable, alignment-free scaffolding of draft genomes with long reads. *Gigascience* **4** (2015).
11. O. Dudchenko, *et al.*, De novo assembly of the *Aedes aegypti* genome using Hi-C yields chromosome-length scaffolds. *Science (1979).* **356**, 92–95 (2017).
12. O. Dudchenko, *et al.*, The Juicebox Assembly Tools module facilitates *de novo* assembly of mammalian genomes with chromosome-length scaffolds for under $1000. [Preprint] (2018). https://doi.org/10.1101/254797.

13. G. Marçais, *et al.*, MUMmer4: A fast and versatile genome alignment system. *PLoS Comput. Biol.* **14**, e1005944 (2018).
14. F. A. Simão, R. M. Waterhouse, P. Ioannidis, E. V. Kriventseva, E. M. Zdobnov, BUSCO: Assessing genome assembly and annotation completeness with single-copy orthologs. *Bioinformatics* **31**, 3210–3212 (2015).
15. A. Dobin, *et al.*, STAR: ultrafast universal RNA-seq aligner. *Bioinformatics* **29**, 15–21 (2013).
16. M. Pertea, *et al.*, StringTie enables improved reconstruction of a transcriptome from RNA-seq reads. *Nat. Biotechnol.* **33**, 290–295 (2015).
17. Y. S. Niknafs, B. Pandian, H. K. Iyer, A. M. Chinnaiyan, M. K. Iyer, TACO produces robust multisample transcriptome assemblies from RNA-seq. *Nat. Methods* **14**, 68–70 (2017).
18. D. Mapleson, L. Venturini, G. Kaithakottil, D. Swarbreck, Efficient and accurate detection of splice junctions from RNA-seq with Portcullis. *Gigascience* **7** (2018).
19. B. J. Haas, Improving the Arabidopsis genome annotation using maximal transcript alignment assemblies. *Nucleic Acids Res.* **31**, 5654–5666 (2003).
20. O. Gotoh, Direct mapping and alignment of protein sequences onto genomic sequence. *Bioinformatics* **24**, 2438–2444 (2008).
21. M. Stanke, S. Waack, Gene prediction with a hidden Markov model and a new intron submodel. *Bioinformatics* **19**, ii215–ii225 (2003).
22. A. Lomsadze, V. Ter-Hovhannisyan, Y. O. Chernoff, M. Borodovsky, Gene identification in novel eukaryotic genomes by self-training algorithm. *Nucleic Acids Res.* **33**, 6494–6506 (2005).
23. B. J. Haas, *et al.*, Automated eukaryotic gene structure annotation using EVidenceModeler and the Program to Assemble Spliced Alignments. *Genome Biol.* **9**, R7 (2008).
24. A. Conesa, *et al.*, Blast2GO: a universal tool for annotation, visualization and analysis in functional genomics research. *Bioinformatics* **21**, 3674–3676 (2005).
25. S. F. Altschul, W. Gish, W. Miller, E. W. Myers, D. J. Lipman, Basic local alignment search tool. *J. Mol. Biol.* **215**, 403–410 (1990).
26. P. Jones, *et al.*, InterProScan 5: genome-scale protein function classification. *Bioinformatics* **30**, 1236–1240 (2014).
27. X. Cui, Z. Lu, S. Wang, J. Jing-Yan Wang, X. Gao, CMsearch: simultaneous exploration of protein sequence space and structure space improves not only protein homology detection but also protein structure prediction. *Bioinformatics* **32**, i332–i340 (2016).
28. E. P. Nawrocki, S. R. Eddy, Infernal 1.1: 100-fold faster RNA homology searches. *Bioinformatics* **29**, 2933–2935 (2013).
29. E. P. Nawrocki, *et al.*, Rfam 12.0: Updates to the RNA families database. *Nucleic Acids Res.* **43**, D130–D137 (2015).
30. T. M. Lowe, S. R. Eddy, tRNAscan-SE: A program for improved detection of transfer RNA genes in genomic sequence. *Nucleic Acids Res.* **25**, 955–964 (1997).
31. H. Li, R. Durbin, Fast and accurate short read alignment with Burrows-Wheeler transform. *Bioinformatics* **25**, 1754–1760 (2009).
32. H. Li, *et al.*, The Sequence Alignment/Map format and SAMtools. *Bioinformatics* **25**, 2078–2079 (2009).
33. A. McKenna, *et al.*, The genome analysis toolkit: A MapReduce framework for analyzing next-generation DNA sequencing data. *Genome Res.* **20**, 1297-1303 (2010).
34. P. Danecek, *et al.*, The variant call format and VCFtools. *Bioinformatics* **27**, 2156–2158 (2011).
35. P. Cingolani, *et al.*, A program for annotating and predicting the effects of single nucleotide polymorphisms, SnpEff. *Fly (Austin).* **6**, 80–92 (2012).
36. B. L. Browning, Y. Zhou, S. R. Browning, A One-Penny Imputed Genome from Next-Generation Reference Panels. *The American Journal of Human Genetics* **103**, 338–348 (2018).
37. B. L. Browning, X. Tian, Y. Zhou, S. R. Browning, Fast two-stage phasing of large-scale sequence data. *The American Journal of Human Genetics* **108**, 1880–1890 (2021).
38. X. Zhou, M. Stephens, Genome-wide efficient mixed-model analysis for association studies. *Nat. Genet.* **44**, 821–824 (2012).
39. S. Purcell, *et al.*, PLINK: A Tool Set for Whole-Genome Association and Population-Based Linkage Analyses. *The American Journal of Human Genetics* **81**, 559–575 (2007).

40. S. D. Turner, qqman: an R package for visualizing GWAS results using Q-Q and manhattan plots. *J. Open Source Softw.* **3**, 731 (2018).
41. D. Kim, J. M. Paggi, C. Park, C. Bennett, S. L. Salzberg, Graph-based genome alignment and genotyping with HISAT2 and HISAT-genotype. *Nat. Biotechnol.* **37**, 907–915 (2019).
42. M. I. Love, W. Huber, S. Anders, Moderated estimation of fold change and dispersion for RNA-seq data with DESeq2. *Genome Biol.* **15**, 550 (2014).
43. Y. Liao, G. K. Smyth, W. Shi, featureCounts: an efficient general purpose program for assigning sequence reads to genomic features. *Bioinformatics* **30**, 923–930 (2014).
44. Y. Liao, G. K. Smyth, W. Shi, The R package Rsubread is easier, faster, cheaper and better for alignment and quantification of RNA sequencing reads. *Nucleic Acids Res.* **47**, e47–e47 (2019).
45. P. Andrade, *et al.*, Regulatory changes in pterin and carotenoid genes underlie balanced color polymorphisms in the wall lizard. *PNAS* **116**, 5633–5642 (2019).
46. J. T. Robinson, *et al.*, Integrative genomics viewer. *Nat. Biotechnol.* **29**, 24-26 (2011).
47. J. Schanda, Ed., *Colorimetry* (Wiley, 2007).
48. C. A. Schneider, W. S. Rasband, K. W. Eliceiri, NIH Image to ImageJ: 25 years of image analysis. *Nat. Methods* **9**, 671–675 (2012).
49. B. Ripley, B. Venables, MASS: support functions and datasets for Venables and Ripley's MASS. *CRAN: contributed packages* [Preprint] (2009).
50. H. Ozeki, S. Ito, K. Wakamatsu, A. J. Thody, Spectrophotometric characterization of eumelanin and pheomelanin in hair. *Pigment Cell Res.* **9**, 265–270 (1996).
51. S. Ito, *et al.*, Usefulness of alkaline hydrogen peroxide oxidation to analyze eumelanin and pheomelanin in various tissue samples: application to chemical analysis of human hair melanins. *Pigment Cell Melanoma Res.* **24**, 605–613 (2011).
52. K. Wakamatsu, S. Ito, J. L. Rees, The usefulness of 4-amino-3-hydroxyphenylalanine as a specific marker of pheomelanin. *Pigment Cell Res.* **15**, 225–232 (2002).
53. M. d'Ischia, *et al.*, Melanins and melanogenesis: methods, standards, protocols. *Pigment Cell Melanoma Res.* **26**, 616–633 (2013).
54. S. Del Bino, *et al.*, Chemical analysis of constitutive pigmentation of human epidermis reveals constant eumelanin to pheomelanin ratio. *Pigment Cell Melanoma Res.* **28**, 707–717 (2015).
55. A. T. L. Lun, *et al.*, EmptyDrops: distinguishing cells from empty droplets in droplet-based single-cell RNA sequencing data. *Genome Biol.* **20**, 63 (2019).
56. Y. Hao, *et al.*, Integrated analysis of multimodal single-cell data. *Cell* **184**, 3573-3587.e29 (2021).
57. E. Cohen, C. Johnson, C. J. Redmond, R. R. Nair, P. A. Coulombe, Revisiting the significance of keratin expression in complex epithelia. *J. Cell Sci.* **135** (2022).
58. M. Nishimura, *et al.*, Extracellular cleavage of collagen XVII is essential for correct cutaneous basement membrane formation. *Hum. Mol. Genet.* **25**, 328–339 (2016).
59. M. Yu, P. Wu, R. B. Widelitz, C.-M. Chuong, The morphogenesis of feathers. *Nature* **420**, 308–312 (2002).
60. M. Inaba, C.-M. Chuong, Avian pigment pattern formation: developmental control of macro- (across the body) and micro- (within a feather) level of pigment patterns. *Front. Cell Dev. Biol.* **8** (2020).
61. V. Nataf, P. Mercier, C. Ziller, N. M. Le Douarin, Novel markers of melanocyte differentiation in the avian embryo. *Exp. Cell Res.* **207**, 171–182 (1993).
62. V. Nataf, *et al.*, Melanoblast/Melanocyte Early Marker (MelEM) is a glutathione S-Transferase subunit. *Exp. Cell Res.* **218**, 394–400 (1995).
63. H. Cheng, G. T. Concepcion, X. Feng, H. Zhang, H. Li, Haplotype-resolved de novo assembly using phased assembly graphs with hifiasm. *Nat. Methods* **18**, 170–175 (2021).
64. M. Alonge, *et al.*, Automated assembly scaffolding using RagTag elevates a new tomato system for high-throughput genome editing. *Genome Biol.* **23**, 258 (2022).
65. A. Shumate, S. L. Salzberg, Liftoff: accurate mapping of gene annotations. *Bioinformatics* **37**, 1639–1643 (2021).

### Supplemental Files

**Data S1.** Differentially expressed genes (DEGs) per cell type between iridescent ($n$ = 2 replicates) and non-iridescent barbules ($n$ = 2 replicates) in the common emerald dove (*Chalcophaps indica*).

**Data S2.** Differentially expressed genes (DEGs) per cell type between iridescent ($n$ = 2 replicates) and non-iridescent barbules ($n$ = 2 replicates) in the mallard (*Anas platyrhynchos*).

**Data S3.** Top upregulated differentially expressed genes (DEGs) for each cell type in the common emerald dove (*Chalcophaps indica*) ($n$ =10,213 cells; $n$ = 12 cell types).

**Data S4.** Top upregulated differentially expressed genes (DEGs) for each cell type in the mallard (*Anas platyrhynchos*) ($n$ = 20,991 cells; $n$ = 9 cell types).

**Data S5.** Gene ontology (GO) enrichment analyses of melanocyte differentially expressed genes between barbule color types (iridescent vs. non-iridescent barbules) for the emerald dove (*Chalcophaps indica*) (adjusted $P$-value ≤ 0.1).

**Data S6.** Gene ontology (GO) enrichment analyses of melanocyte differentially expressed genes between barbule color types (iridescent vs. non-iridescent barbules) for the mallard (*Anas platyrhynchos*) (adjusted $P$-value ≤ 0.1).

### SUPPLEMENTAL TABLES

**Table S1. Genome assembly, annotation, and BUSCO statistics of the Indian peafowl genome**

| | | |
|---|---|---|
| **Genome assembly and sequencing statistics** | | |
| **10x linked-reads** | | |
| | Number of reads | 262 153 402 |
| | Average coverage | 60.03 |
| | Assembly size (bp) | 372 |
| **Nanopore** | | |
| | Number of reads | 4 599 390 |
| | Average coverage | 32.99 |
| | Contig N50 (kb) | 16.62 |
| **Hi-C** | | |
| | Number of reads | 317 826 943 |
| | Number of contacts | 261 213 953 |
| **Final assembly** | | |
| | Assembly size (Gb) | 1.04 |
| | Contig N50 (Mb) | 3.57 |
| | Contig L50 | 86 |
| | Scaffolds | 813 |
| | Scaffold N50 (Mb) | 93.07 |
| | Scaffold L50 | 4 |
| **Genome annotation report** | | |
| | Number of protein-coding genes | 21 618 |
| | Median gene length (bp) | 9 416 |
| | Number of transcripts | 32 889 |
| | Number of exons | 224 330 |
| | Number of coding exons | 213 541 |
| | Coding GC content | 52.62% |
| | Median UTR length (bp) | 1692 |
| | Median intron length (bp) | 749 |
| | Exons per transcript | 11.07 |
| | Transcripts per gene | 1.52 |
| | Multi-exonic transcripts | 0.92 |
| | Gene density (genes/Mb) | 20.85 |
| **BUSCO analysis (8,338 avian genes; aves_odb10)** | | |
| Complete (%) | | 96.2 |
| | singleton (%) | 95.8 |
| | duplicated (%) | 0.4 |
| Fragmented (%) | | 0.8 |
| Missing (%) | | 3.0 |

**Table S2. List of genes and mutations associated with the Indian peafowl varieties.**

| Variety | Gene | Acronym | Position | Type[a] | Protein change[b] | Evolutionary conservation (*n* = species)[c] |
|---|---|---|---|---|---|---|
| **Mendelian** | | | | | | |
| Charcoal | *Lysosomal trafficking regulator* | *LYST* | 3:38,734,068 | Missense | Arg1556His | 99% (135) |
| White | *Endothelin receptor B subtype 2* | *EDNRB2* | 4:80,919,920 | Nonsense | Trp206* | |
| Black-shoulder | *Melanocortin 1 receptor* | *MC1R* | 6:53,715,975 | Frameshift (Δ1) | Ser165Ala | |
| Black-shoulder | *Melanocortin 1 receptor* | *MC1R* | 6:53,716,234 | Missense | Trp251Arg | 100% (47) |
| Bronze | *Hermansky-Pudlak syndrome type 3* | *HPS3* | 7:53,510,213 | Insertion (Δ12) | Cys524* | |
| Opal | *Melanophilin* | *MLPH* | 8:34,112,863 | Deletion (Δ483) | Ser245Gly | |
| Ultramarine | *Microphthalmia-associated transcription factor* | *MITF* | 10:15,139,593 | Missense | Glu205Lys | 100% (139) |
| Purple | *Tyrosinase-related protein 1 enzyme* | *TYRP1* | Z:31,260,949 | Missense | Phe243Ser | 100% (288) |
| Violet | *Tyrosinase-related protein 1 enzyme* | *TYRP1* | Z:31,264,368 | Nonsense | Arg373* | |
| Cameo | *Adaptor-related protein complex 3 sigma 1 subunit* | *AP3S1* | Z:74,408,869 | Missense | Leu71Phe | 100% (129) |
| **Combinations of mutations**[d] | | | | | | |
| Taupe | | Purple + *MLPH* | 8:34,104,916 | *nonsense* | Arg581* | |
| Hazel | | Bronze + Purple | | | | |
| Raw-umber | | Cameo + Opal | | | | |
| Peach | | Cameo + Purple | | | | |

[a]The delta (Δ) represents the size of the insertion or deletion

[b]The asterisk represents mutations that give rise to premature STOP codons

[c]The proportion of amino acid residues that match the wild-type allele across whole-genome alignments of proteins across birds. In parentheses, the number of species included in the alignment are given.

[d]Color varieties resulting from combinations of loci are identified by the mutants they derive from, with the exception of *taupe* where another *MLPH* mutation contributes to the phenotype.

**Table S3. Quantification of structural parameters in feather barbules across eyespot regions.**

| Phenotype | Area | *N*[a] | | *a*[b] | | *b*[c] | | *c*[d] | | *Dm*[e] | | *Da*[f] | | *Lm*[g] | | Aspect ratio[h] | Chroma[i] | HEX[j] |
|---|---|---|---|---|---|---|---|---|---|---|---|---|---|---|---|---|---|---|
| | | Mean | SD | Mean | SD | Mean | SD | Mean | SD | Mean | SD | Mean | SD | Mean | SD | | | |
| Wild-type | 1 | 7.7 | 1.8 | 147.4 | 15.0 | 225.4 | 18.4 | 97.5 | 14.8 | 105.8 | 11.8 | 39.0 | 11.6 | 692.7 | 100.6 | 6.6 | 54.7 | #007C7F |
| Wild-type | 2 | 7.5 | 1.4 | 174.0 | 12.8 | 158.3 | 17.2 | 101.4 | 9.5 | 106.1 | 9.5 | 52.3 | 14.3 | 763.4 | 115.9 | 7.3 | 70.3 | #00804D |
| Wild-type | 3 | 5.4 | 1.0 | 198.7 | 21.7 | 178.9 | 21.7 | 141.0 | 14.6 | 110.8 | 10.5 | 48.9 | 15.0 | 769.9 | 105.0 | 7.0 | 32.8 | #91643C |
| Wild-type | 4 | 5.3 | 1.2 | 200.5 | 26.3 | 166.1 | 20.4 | 130.9 | 10.2 | 116.3 | 10.9 | 64.6 | 11.2 | 723.7 | 111.9 | 6.3 | 42.7 | #A47E39 |
| Wild-type | 5 | 4.7 | 0.9 | 197.2 | 32.8 | 177.6 | 20.5 | 112.9 | 18.1 | 114.7 | 9.5 | 55.8 | 12.9 | 798.1 | 126.1 | 7.0 | 25.5 | #775071 |
| Wild-type | 6 | 4.6 | 0.9 | 191.9 | 38.2 | 162.1 | 14.1 | 130.5 | 16.1 | 119.9 | 9.6 | 65.9 | 12.9 | 778.8 | 126.1 | 6.5 | 19.9 | #8A5E6A |
| Ultramarine | 1 | 6.3 | 1.6 | 148.1 | 15.3 | 149.7 | 21.0 | 83.2 | 12.0 | 103.8 | 8.4 | 57.3 | 14.0 | 765.8 | 112.6 | 7.5 | 26.1 | #2A2273 |
| Ultramarine | 2 | 5.8 | 1.4 | 159.0 | 15.5 | 153.6 | 15.5 | 88.6 | 11.1 | 108.5 | 9.8 | 49.9 | 12.4 | 738.5 | 116.2 | 6.9 | 35.2 | #004476 |
| Ultramarine | 3 | 4.9 | 1.1 | 199.0 | 34.8 | 143.1 | 18.5 | 122.4 | 14.1 | 115.8 | 9.2 | 58.7 | 16.5 | 740.5 | 101.2 | 6.4 | 13.2 | #526554 |
| Ultramarine | 4 | 4.4 | 1.1 | 177.5 | 28.2 | 150.2 | 19.3 | 100.0 | 14.2 | 119.6 | 7.9 | 36.2 | 10.8 | 691.3 | 115.7 | 5.8 | 35.2 | #5A8853 |
| Ultramarine | 5 | 4.9 | 0.8 | 179.2 | 20.6 | 160.1 | 16.2 | 109.3 | 13.0 | 120.5 | 10.4 | 38.1 | 13.8 | 748.1 | 116.1 | 6.3 | 36.5 | #387546 |
| Ultramarine | 6 | 4.1 | 1.2 | 200.8 | 25.1 | 162.4 | 15.4 | 110.7 | 17.9 | 127.6 | 11.7 | 32.6 | 12.5 | 837.8 | 108.3 | 6.6 | 39.2 | #547738 |
| Purple | 1 | 5.8 | 2.1 | 179.9 | 30.3 | 157.6 | 19.6 | 109.0 | 18.9 | 86.7 | 14.2 | 60.6 | 15.8 | 766.4 | 134.3 | 9.1 | 26.1 | #004869 |
| Purple | 2 | 7.2 | 1.0 | 180.9 | 22.0 | 162.6 | 22.3 | 104.8 | 13.8 | 96.5 | 16.5 | 57.0 | 17.2 | 713.0 | 118.5 | 7.6 | 50.3 | #007866 |
| Purple | 3 | 4.5 | 0.6 | 196.5 | 37.0 | 145.8 | 23.4 | 141.5 | 34.7 | 78.3 | 13.3 | 55.2 | 17.3 | 707.1 | 129.6 | 9.3 | 16.5 | #7B5F4E |
| Purple | 4 | 4.5 | 0.8 | 187.2 | 31.3 | 140.0 | 23.2 | 140.4 | 19.3 | 83.0 | 12.6 | 63.8 | 18.7 | 700.5 | 149.5 | 8.7 | 27.4 | #BD8556 |
| Purple | 5 | 4.3 | 0.9 | 194.6 | 47.6 | 155.7 | 26.8 | 132.0 | 19.9 | 88.6 | 13.7 | 57.4 | 20.1 | 818.8 | 182.0 | 9.4 | 4.9 | #685D5C |
| Purple | 6 | 4.4 | 1.0 | 195.2 | 32.2 | 153.5 | 33.4 | 129.6 | 18.9 | 87.8 | 15.6 | 65.3 | 19.8 | 765.0 | 136.0 | 9.0 | 4.9 | #6A5F5F |
| Violet | 1 | 3.8 | 1.1 | 123.5 | 30.6 | 117.1 | 16.5 | 92.4 | 14.3 | 74.4 | 11.0 | 28.4 | 14.2 | 784.2 | 131.1 | 10.8 | 18.6 | #2D324D |
| Violet | 2 | 5.7 | 2.1 | 172.2 | 27.1 | 159.8 | 29.6 | 117.0 | 19.8 | 71.0 | 12.3 | 35.0 | 8.4 | 633.7 | 196.7 | 9.2 | 30.6 | #506E1E |
| Violet | 3 | 4.6 | 1.1 | 206.2 | 46.3 | 150.0 | 18.9 | 162.3 | 29.7 | 78.3 | 12.2 | 44.9 | 13.4 | 728.1 | 120.9 | 9.5 | 12.2 | #695449 |
| Violet | 4 | 4.7 | 1.0 | 171.2 | 33.2 | 154.2 | 29.7 | 140.2 | 17.7 | 81.0 | 13.6 | 54.0 | 14.3 | 790.3 | 123.3 | 10.0 | 14.4 | #667B68 |
| Violet | 5 | 3.4 | 1.2 | 181.3 | 38.0 | 143.4 | 22.5 | 165.9 | 21.6 | 84.4 | 11.3 | 44.0 | 14.4 | 869.8 | 120.3 | 10.4 | 12.5 | #5C4755 |
| Violet | 6 | 4.0 | 0.9 | 183.1 | 30.3 | 138.4 | 21.2 | 134.1 | 15.1 | 88.7 | 12.5 | 47.8 | 15.4 | 774.6 | 155.0 | 8.9 | 21.7 | #42684F |
| Opal | 1 | 1.7 | 1.1 | 196.7 | 21.2 | 168.7 | 22.5 | 114.0 | 25.5 | 107.0 | 16.5 | 29.6 | 11.0 | 654.5 | 104.7 | 6.3 | 11.8 | #1B3741 |
| Opal | 2 | 2.9 | 1.0 | 196.7 | 21.2 | 168.6 | 22.6 | 107.6 | 22.6 | 107.0 | 16.5 | 29.7 | 11.0 | 654.1 | 105.4 | 6.3 | 24.5 | #2D563A |
| Opal | 3 | 2.7 | 1.1 | 210.2 | 28.1 | 173.1 | 23.0 | 131.3 | 23.4 | 122.3 | 14.2 | 46.4 | 18.5 | 697.7 | 118.7 | 5.8 | 9.6 | #422F34 |
| Opal | 4 | 2.1 | 0.7 | 220.3 | 25.2 | 174.5 | 21.7 | 134.3 | 18.5 | 119.8 | 12.8 | 42.5 | 15.5 | 630.0 | 108.4 | 5.3 | 19.1 | #623A3A |
| Opal | 5 | 2.1 | 1.1 | 214.4 | 37.3 | 169.0 | 23.2 | 141.1 | 23.9 | 119.9 | 15.5 | 47.7 | 15.1 | 685.8 | 109.1 | 5.8 | 18.0 | #663833 |
| Opal | 6 | 0.8 | 0.8 | 216.5 | 26.3 | 199.5 | 33.3 | 166.8 | 24.1 | 132.7 | 16.9 | 51.8 | 22.8 | 653.7 | 98.7 | 5.0 | 9.2 | #493833 |
| Bronze | 1 | 1.9 | 1.1 | 232.8 | 49.1 | 237.5 | 46.5 | 82.3 | 35.2 | 156.5 | 44.9 | 46.2 | 26.3 | 187.6 | 39.9 | 1.3 | 1.5 | #373633 |

| | | | | | | | | | | | | | | | | | | |
|---|---|---|---|---|---|---|---|---|---|---|---|---|---|---|---|---|---|---|
| Bronze | 2 | 3.1 | 1.3 | 264.5 | 46.2 | 249.0 | 48.9 | 146.5 | 33.7 | 176.7 | 35.2 | 75.7 | 26.5 | 190.1 | 50.1 | 1.1 | 2.3 | #3A3639 |
| Bronze | 3 | 1.7 | 1.0 | 285.9 | 69.6 | 294.3 | 68.7 | 183.3 | 38.6 | 183.9 | 38.4 | 73.2 | 31.0 | 226.0 | 67.1 | 1.3 | 6.9 | #42493F |
| Bronze | 4 | 2.0 | 0.8 | 284.8 | 64.4 | 232.2 | 56.4 | 177.0 | 32.8 | 152.3 | 37.7 | 62.1 | 26.5 | 201.4 | 65.9 | 1.4 | 3.8 | #3B3F3A |
| Bronze | 5 | 1.9 | 0.9 | 228.5 | 41.7 | 199.1 | 42.4 | 135.7 | 34.7 | 130.8 | 34.5 | 55.9 | 20.5 | 234.7 | 88.3 | 1.9 | 5.2 | #3A4440 |
| Bronze | 6 | 1.9 | 1.0 | 303.2 | 57.8 | 246.5 | 53.2 | 175.5 | 36.2 | 180.8 | 47.5 | 51.2 | 20.7 | 174.0 | 67.7 | 1.1 | 1.7 | #3A3D3D |
| Cameo | 1 | 0.6 | 0.7 | 604.8 | 433.4 | 525.7 | 316.9 | 88.9 | 37.3 | 133.5 | 24.2 | 42.5 | 14.0 | 160.7 | 55.3 | 1.3 | 1.0 | #3F3D3E |
| Cameo | 2 | 0.8 | 0.8 | 319.5 | 193.3 | 300.2 | 205.6 | 167.1 | 38.8 | 132.9 | 30.6 | 49.3 | 14.0 | 197.2 | 80.1 | 1.6 | 1.9 | #3D3938 |
| Cameo | 3 | 0.8 | 0.8 | 313.9 | 134.9 | 264.3 | 127.5 | 160.8 | 46.8 | 127.5 | 32.2 | 41.8 | 17.8 | 186.1 | 56.4 | 1.5 | 1.5 | #413F3D |
| Cameo | 4 | 0.8 | 0.7 | 280.2 | 86.6 | 281.4 | 130.5 | 163.8 | 48.8 | 130.3 | 37.0 | 64.5 | 26.4 | 210.6 | 87.9 | 1.8 | 4.0 | #49443F |
| Cameo | 5 | 0.8 | 0.8 | 301.4 | 104.7 | 289.9 | 121.9 | 170.5 | 37.8 | 152.0 | 29.8 | 59.0 | 17.7 | 229.2 | 53.1 | 1.6 | 3.4 | #433E3A |
| Cameo | 6 | 1.0 | 0.9 | 359.4 | 169.7 | 347.9 | 150.7 | 178.3 | 47.7 | 160.7 | 37.0 | 51.1 | 17.0 | 215.0 | 59.8 | 1.4 | 3.2 | #3F3936 |
| Charcoal | 1 | 0.5 | 0.6 | 388.2 | 211.2 | 491.4 | 255.6 | 148.0 | 47.4 | 189.1 | 62.3 | 44.9 | 22.9 | 251.1 | 122.8 | 1.5 | 1.5 | #343131 |
| Charcoal | 2 | 0.3 | 0.6 | 397.2 | 248.7 | 448.6 | 318.9 | 201.4 | 51.7 | 200.2 | 86.3 | 40.6 | 21.9 | 271.6 | 115.5 | 1.7 | 2.6 | #332D2C |
| Charcoal | 3 | 0.6 | 0.6 | 354.5 | 202.8 | 457.1 | 256.4 | 148.8 | 49.6 | 215.9 | 78.5 | 35.0 | 16.3 | 274.7 | 151.0 | 1.5 | 2.7 | #322D2B |
| Charcoal | 4 | 0.6 | 0.9 | 458.7 | 401.3 | 483.5 | 326.2 | 192.3 | 70.7 | 152.5 | 72.5 | 31.4 | 20.7 | 308.3 | 143.4 | 2.5 | 1.8 | #33302E |
| Charcoal | 5 | 0.8 | 0.9 | 467.1 | 333.9 | 536.7 | 339.4 | 180.6 | 42.7 | 206.6 | 66.8 | 28.8 | 11.7 | 270.8 | 104.4 | 1.5 | 1.1 | #363433 |
| Charcoal | 6 | 0.3 | 0.6 | 663.0 | 428.8 | 387.2 | 215.6 | 162.1 | 45.5 | 227.2 | 68.6 | 38.5 | 16.5 | 315.8 | 110.4 | 1.6 | 1.9 | #373332 |

[a]Number of melanosome layers

[b]Distance between layers

[c]Distance between melanosomes within a layer

[d]Cortex thickness

[e]Melanosome diameter

[f]Air pocket diameter

[g]Melanosome length

[h]Calculated as the ratio of melanosome length (*Lm*) and diameter (*Dm*)

[i]Measure of color saturation or intensity

[j]RGB-based hexadecimal color code

**Table S4. Linear discriminant analysis (LDA) per group (all mutants, iridescent, and non-iridescent).**

| | All mutants | | | | Iridescent | | | | Non-iridescent | |
|---|---|---|---|---|---|---|---|---|---|---|
| | **LD1** | **LD2** | **LD3** | **LD4** | **LD1** | **LD2** | **LD3** | **LD4** | **LD1** | **LD2** |
| **Variance explained[a]** | | | | | | | | | | |
| | 0.899 | 0.060 | 0.024 | 0.014 | 0.628 | 0.252 | 0.095 | 0.024 | 0.851 | 0.149 |
| **Pearson's *r*[b]** | | | | | | | | | | |
| *N*[d] | 0.852 | 0.146 | **-0.395** | -0.221 | -0.341 | **0.807** | 0.198 | -0.056 | -0.746 | **0.642** |
| *a*[e] | -0.768 | -0.383 | 0.103 | -0.255 | 0.381 | -0.414 | 0.345 | -0.068 | 0.634 | -0.252 |
| *b*[f] | -0.804 | -0.489 | 0.038 | **-0.289** | 0.543 | -0.020 | 0.356 | **-0.723** | 0.836 | -0.213 |
| *c*[g] | -0.580 | -0.138 | 0.218 | -0.186 | -0.232 | -0.487 | 0.081 | -0.402 | 0.318 | -0.006 |
| *Dm*[h] | -0.823 | -0.405 | -0.330 | 0.145 | **0.941** | 0.194 | -0.019 | -0.118 | 0.663 | 0.510 |
| *Da*[i] | 0.022 | **0.594** | -0.413 | -0.291 | -0.204 | 0.311 | **0.704** | 0.104 | -0.765 | 0.172 |
| *Lm*[j] | **0.986** | -0.115 | 0.057 | 0.058 | -0.432 | 0.512 | -0.119 | 0.020 | **0.865** | 0.213 |
| ***P*-value[c]** | | | | | | | | | | |
| *N* | 1.58E-14 | 3.22E-01 | **5.50E-03** | 1.32E-01 | 6.53E-02 | **7.29E-08** | 2.94E-01 | 7.67E-01 | 3.78E-04 | **4.04E-03** |
| *a* | 1.88E-10 | 7.26E-03 | 4.87E-01 | 8.03E-02 | 3.79E-02 | 2.28E-02 | 6.21E-02 | 7.22E-01 | 4.75E-03 | 3.12E-01 |
| *b* | 6.13E-12 | 4.23E-04 | 7.98E-01 | **4.61E-02** | 1.93E-03 | 9.15E-01 | 5.37E-02 | **6.50E-06** | 1.55E-05 | 3.97E-01 |
| *c* | 1.59E-05 | 3.48E-01 | 1.37E-01 | 2.06E-01 | 2.17E-01 | 6.30E-03 | 6.69E-01 | 2.77E-02 | 1.99E-01 | 9.82E-01 |
| *Dm* | 7.06E-13 | 4.26E-03 | 2.18E-02 | 3.26E-01 | **1.18E-14** | 3.04E-01 | 9.22E-01 | 5.35E-01 | 2.72E-03 | 3.06E-02 |
| *Da* | 8.83E-01 | **8.39E-06** | 3.50E-03 | 4.50E-02 | 2.79E-01 | 9.40E-02 | **1.41E-05** | 5.86E-01 | 2.16E-04 | 4.94E-01 |
| *Lm* | **1.25E-37** | 4.35E-01 | 7.02E-01 | 6.93E-01 | 1.71E-02 | 3.82E-03 | 5.30E-01 | 9.18E-01 | **3.53E-06** | 3.95E-01 |

Numbers in bold represent the top correlated variable with the respective axis, for axes explaining above 5% of variance

[a]Proportion of the total between-group variance

[b]Correlation

[c]Significance of mean values of each photonic lattice parameter per area per mutant (Z-score standardized) with LDA axes

[d]Number of melanosome layers

[e]Distance between layers

[f]Distance between melanosomes within a layer

[g]Cortex thickness

[h]Melanosome diameter

[i]Air pocket diameter

[j]Melanosome length

**Table S5. Proportion of variance explained per variable and phenotype relative to wild-type in linear discriminant analysis (LDA).**

| | Ultramarine | | Purple | | Violet | | Opal | | Bronze | | Cameo | | Charcoal | |
|---|---|---|---|---|---|---|---|---|---|---|---|---|---|---|
| | **Pearson's *r*** | ***P*-value** | **Pearson's *r*** | ***P*-value** | **Pearson's *r*** | ***P*-value** | **Pearson's *r*** | ***P*-value** | **Pearson's *r*** | ***P*-value** | **Pearson's *r*** | ***P*-value** | **Pearson's *r*** | ***P*-value** |
| *N*[a] | -0.385 | 2.16E-01 | -0.323 | 3.06E-01 | **-0.589** | **4.41E-02** | **-0.868** | **1.13E-04** | **-0.895** | **8.46E-05** | **-0.943** | **4.42E-06** | **-0.949** | **2.61E-06** |
| *a*[b] | -0.216 | 4.99E-01 | 0.146 | 6.51E-01 | -0.265 | 4.04E-01 | **0.638** | **2.69E-02** | **0.866** | **2.70E-04** | **0.748** | **5.13E-03** | **0.882** | **1.48E-04** |
| *b*[c] | **-0.659** | **1.96E-02** | **-0.624** | **3.01E-02** | **-0.683** | **1.43E-02** | -0.083 | 8.27E-01 | **0.789** | **2.27E-03** | **0.771** | **3.29E-03** | **0.972** | **1.24E-07** |
| *c*[d] | -0.537 | 7.19E-02 | 0.235 | 4.61E-01 | 0.361 | 2.49E-01 | 0.362 | 2.54E-01 | 0.497 | 1.00E-01 | **0.598** | **4.00E-02** | **0.822** | **1.05E-03** |
| *Dm*[e] | 0.291 | 3.59E-01 | **-0.948** | **2.83E-06** | **-0.956** | **1.27E-06** | 0.319 | 2.31E-01 | **0.882** | **1.46E-04** | **0.822** | **1.03E-03** | **0.931** | **1.13E-05** |
| *Da*[f] | -0.457 | 1.36E-01 | 0.375 | 2.29E-01 | -0.571 | 5.27E-02 | **-0.581** | **4.05E-02** | 0.300 | 3.43E-01 | -0.173 | 5.90E-01 | **-0.766** | **3.70E-03** |
| *Lm*[g] | -0.011 | 9.74E-01 | -0.122 | 7.06E-01 | 0.080 | 8.04E-01 | **-0.808** | **5.29E-04** | **-0.997** | **2.18E-12** | **-0.996** | **5.50E-12** | **-0.994** | **6.05E-11** |

Significant values are represented in bold

[a]Number of melanosome layers

[b]Distance between layers

[c]Distance between melanosomes within a layer

[d]Cortex thickness

[e]Melanosome diameter

[f]Air pocket diameter

[g]Melanosome length

Table S6. Quantification of melanin in mature feathers of wild-type and mutant peacocks.

| Sample | Region | Pheno | S-350 (/mg)[a] | | AHPO (ng/mg)[b] | | | HI (ng/mg)[c] | | TM[d] (µg/mg) | EM[e] (µg/mg) | BZ[f] (µg/mg) | BT[g] (µg/mg) | PM[h] (µg/mg) |
|---|---|---|---|---|---|---|---|---|---|---|---|---|---|---|
| | | | A500 | A650 | PTCA | PDCA | TTCA | 4-AHP | 3-AHP | | | | | |
| peacock247 | Neck | Wild-type | 1.851 | 0.601 | 1985 | 277 | 465 | 39 | 96 | 187 | 159 | 15.8 | 0.4 | 16.2 |
| peacock411 | Neck | Wild-type | 2.229 | 0.749 | 2188 | 288 | 451 | 37 | 66 | 225 | 175 | 15.3 | 0.3 | 15.7 |
| peacock282 | Neck | Wild-type | 1.820 | 0.571 | 1952 | 235 | 427 | 18 | 19 | 184 | 156 | 14.5 | 0.2 | 14.7 |
| peacock479 | Neck | Wild-type | 1.593 | 0.498 | 1685 | 248 | 326 | 11 | 16 | 161 | 135 | 11.1 | 0.1 | 11.2 |
| peacock506 | Neck | Ultramarine | 2.141 | 0.745 | 2193 | 234 | 458 | 34 | 96 | 216 | 175 | 15.6 | 0.3 | 15.9 |
| peacock510 | Neck | Ultramarine | 1.963 | 0.657 | 1779 | 193 | 409 | 34 | 175 | 198 | 142 | 13.9 | 0.3 | 14.2 |
| peacock520 | Neck | Ultramarine | 1.519 | 0.491 | 1955 | 183 | 350 | 19 | 66 | 153 | 156 | 11.9 | 0.2 | 12.1 |
| peacock197 | Neck | Purple | 1.063 | 0.344 | 947 | 200 | 198 | 63 | 30 | 107 | 76 | 6.7 | 0.6 | 7.3 |
| peacock236 | Neck | Purple | 0.944 | 0.292 | 927 | 205 | 228 | 86 | 54 | 95 | 74 | 7.8 | 0.8 | 8.5 |
| peacock314 | Neck | Purple | 0.843 | 0.260 | 684 | 167 | 202 | 64 | 46 | 85 | 55 | 6.9 | 0.6 | 7.4 |
| peacock316 | Neck | Purple | 0.857 | 0.276 | 784 | 154 | 192 | 69 | 44 | 87 | 63 | 6.5 | 0.6 | 7.1 |
| peacock274 | Neck | Violet | 0.617 | 0.191 | 549 | 128 | 161 | 93 | 50 | 62 | 44 | 5.5 | 0.8 | 6.3 |
| peacock278 | Neck | Violet | 0.638 | 0.194 | 567 | 127 | 164 | 83 | 50 | 64 | 45 | 5.6 | 0.7 | 6.3 |
| peacock150 | Neck | Violet | 0.413 | 0.122 | 508 | 108 | 126 | 23 | 21 | 42 | 41 | 4.3 | 0.2 | 4.5 |
| peacock476 | Neck | Violet | 0.448 | 0.134 | 525 | 104 | 115 | 21 | 23 | 45 | 42 | 3.9 | 0.2 | 4.1 |
| peacock239 | Neck | Opal | 0.532 | 0.176 | 481 | 56 | 249 | 40 | 34 | 54 | 38 | 8.5 | 0.4 | 8.8 |
| peacock249 | Neck | Opal | 0.397 | 0.129 | 384 | 54 | 214 | 27 | 30 | 40 | 31 | 7.3 | 0.2 | 7.5 |
| peacock302 | Neck | Opal | 0.524 | 0.172 | 442 | 63 | 278 | 77 | 44 | 53 | 35 | 9.5 | 0.7 | 10.1 |
| peacock238 | Neck | Opal | 0.561 | 0.186 | 513 | 74 | 257 | 45 | 35 | 57 | 41 | 8.7 | 0.4 | 9.1 |
| peacock306 | Neck | Opal | 0.500 | 0.165 | 420 | 61 | 242 | 56 | 39 | 51 | 34 | 8.2 | 0.5 | 8.7 |
| peacock174 | Neck | Bronze | 0.579 | 0.193 | 496 | 69 | 322 | 653 | 215 | 58 | 40 | 10.9 | 5.9 | 16.8 |
| peacock192 | Neck | Bronze | 0.671 | 0.221 | 551 | 81 | 319 | 552 | 177 | 68 | 44 | 10.8 | 5.0 | 15.8 |
| peacock196 | Neck | Bronze | 0.581 | 0.187 | 514 | 72 | 292 | 601 | 185 | 59 | 41 | 9.9 | 5.4 | 15.3 |
| peacock175 | Neck | Bronze | 0.750 | 0.248 | 562 | 100 | 303 | 440 | 154 | 76 | 45 | 10.3 | 4.0 | 14.3 |
| peacock284 | Neck | Bronze | 0.642 | 0.207 | 515 | 76 | 307 | 582 | 202 | 65 | 41 | 10.4 | 5.2 | 15.7 |
| peacock286 | Neck | Bronze | 0.768 | 0.256 | 588 | 92 | 319 | 526 | 179 | 78 | 47 | 10.8 | 4.7 | 15.6 |
| peacock189 | Neck | Cameo | 0.227 | 0.072 | 195 | 28 | 188 | 745 | 228 | 23 | 16 | 6.4 | 6.7 | 13.1 |
| peacock191 | Neck | Cameo | 0.191 | 0.056 | 191 | 27 | 143 | 648 | 188 | 19 | 15 | 4.9 | 5.8 | 10.7 |
| peacock224 | Neck | Cameo | 0.178 | 0.049 | 125 | 25 | 196 | 655 | 238 | 18 | 10 | 6.7 | 5.9 | 12.6 |
| peacock297 | Neck | Cameo | 0.237 | 0.069 | 209 | 32 | 183 | 660 | 223 | 24 | 17 | 6.2 | 5.9 | 12.2 |
| peacock295 | Neck | Cameo | 0.239 | 0.070 | 206 | 32 | 203 | 762 | 249 | 24 | 16 | 6.9 | 6.9 | 13.8 |
| peacock234 | Neck | Charcoal | 0.794 | 0.256 | 804 | 102 | 461 | 85 | 38 | 80 | 64 | 15.7 | 0.8 | 16.4 |
| peacock265 | Neck | Charcoal | 0.483 | 0.156 | 479 | 65 | 275 | 43 | 19 | 49 | 38 | 9.4 | 0.4 | 9.7 |

| | | | | | | | | | | | | | | |
|---|---|---|---|---|---|---|---|---|---|---|---|---|---|---|
| peacock266 | Neck | Charcoal | 0.530 | 0.171 | 494 | 68 | 261 | 33 | 17 | 54 | 40 | 8.9 | 0.3 | 9.2 |
| peacock508 | Neck | Charcoal | 0.682 | 0.223 | 722 | 76 | 369 | 304 | 110 | 69 | 58 | 12.5 | 2.7 | 12.8 |
| peacock332 | Neck | Black-shoulder | 1.853 | 0.615 | 1673 | 248 | 425 | 37 | 94 | 187 | 134 | 14.5 | 0.3 | 14.7 |
| peacock337 | Neck | Black-shoulder | 1.889 | 0.625 | 1939 | 283 | 450 | 28 | 92 | 191 | 155 | 15.3 | 0.2 | 15.5 |
| peacock237 | Neck | Black-shoulder | 2.059 | 0.689 | 2080 | 282 | 463 | 37 | 73 | 208 | 166 | 15.7 | 0.3 | 16.0 |
| peacock247 | Wing | Wild-type | 0.672 | 0.210 | 802 | 119 | 180 | 188 | 88 | 68 | 64 | 6.1 | 1.7 | 7.8 |
| peacock427 | Wing | Wild-type | 0.595 | 0.185 | 774 | 114 | 168 | 43 | 40 | 60 | 62 | 5.7 | 0.4 | 6.1 |
| peacock411 | Wing | Wild-type | 0.594 | 0.186 | 796 | 107 | 172 | 112 | 38 | 60 | 64 | 5.8 | 1.0 | 6.9 |
| peacock282 | Wing | Wild-type | 0.607 | 0.194 | 717 | 97 | 190 | 74 | 27 | 61 | 57 | 6.5 | 0.7 | 7.1 |
| peacock332 | Wing | Black-shoulder | 1.249 | 0.392 | 1247 | 199 | 180 | 15 | 63 | 126 | 100 | 6.1 | 0.1 | 6.3 |
| peacock337 | Wing | Black-shoulder | 1.227 | 0.378 | 1376 | 206 | 205 | 16 | 55 | 124 | 110 | 7.0 | 0.1 | 7.1 |
| peacock237 | Wing | Black-shoulder | 1.223 | 0.382 | 1362 | 204 | 177 | 13 | 35 | 124 | 109 | 6.0 | 0.1 | 6.1 |

[a]Soluene-350 solubilization
[b]Alkaline hydrogen peroxide oxidation
[c]Hydroiodic acid hydrolysis
[d]Total melanin (A500 × 101)
[e]Eumelanin (PTCA × 0.080)
[f]Benzothiazole (TTCA × 0.034)
[g]Benzothiazine (4-AHP × 0.009)
[h]Pheomelanin (BZ + BT)

**Table S7. Statistics from whole-genome sequencing of peafowl individuals.**

| ID | Species | Body color phenotype | Wing phenotype | Coverage | Number of reads | Mapped reads (%) | Properly paired (%) |
|---|---|---|---|---|---|---|---|
| peacock001 | *Pavo cristatus* | bronze | | 8.3 | 71 989 677 | 99.0 | 98.8 |
| peacock002 | *Pavo cristatus* | bronze | | 9.1 | 76 720 800 | 98.8 | 96.3 |
| peacock003 | *Pavo cristatus* | bronze | black-shoulder | 9.5 | 78 832 417 | 98.9 | 95.9 |
| peacock004 | *Pavo cristatus* | bronze | black-shoulder | 9.8 | 84 976 525 | 98.9 | 96.2 |
| peacock005 | *Pavo cristatus* | bronze | black-shoulder | 11.2 | 95 514 878 | 98.5 | 96.1 |
| peacock006 | *Pavo cristatus* | bronze | | 9.7 | 82 144 664 | 98.6 | 95.1 |
| peacock007 | *Pavo cristatus* | bronze | | 10.8 | 90 931 068 | 98.6 | 95.3 |
| peacock008 | *Pavo cristatus* | bronze | | 9.8 | 82 514 008 | 98.9 | 95.3 |
| peacock009 | *Pavo cristatus* | bronze | | 9.4 | 79 772 763 | 98.9 | 96.1 |
| peacock010 | *Pavo cristatus* | bronze | | 8.7 | 74 944 196 | 98.4 | 96.3 |
| peacock011 | *Pavo cristatus* | bronze | | 8.9 | 74 850 903 | 99.1 | 95.2 |
| peacock012 | *Pavo cristatus* | bronze | | 9.1 | 77 517 989 | 99.1 | 96.7 |
| peacock013 | *Pavo cristatus* | cameo | black-shoulder | 7.8 | 68 595 974 | 98.8 | 96.6 |
| peacock014 | *Pavo cristatus* | bronze | | 9.3 | 80 469 878 | 98.8 | 95.4 |
| peacock015 | *Pavo cristatus* | cameo | black-shoulder | 10.2 | 85 935 164 | 98.7 | 95.9 |
| peacock016 | *Pavo cristatus* | cameo | black-shoulder | 8.8 | 75 138 376 | 98.0 | 95.0 |
| peacock017 | *Pavo cristatus* | cameo | | 8.1 | 71 067 052 | 98.5 | 94.2 |
| peacock018 | *Pavo cristatus* | cameo | | 10.1 | 87 030 285 | 98.8 | 95.4 |
| peacock019 | *Pavo cristatus* | cameo | | 9.1 | 79 153 902 | 98.3 | 96.0 |
| peacock020 | *Pavo cristatus* | cameo | black-shoulder | 11.3 | 94 994 710 | 98.9 | 94.9 |
| peacock021 | *Pavo cristatus* | cameo | | 8.1 | 72 299 953 | 98.8 | 96.2 |
| peacock022 | *Pavo cristatus* | cameo | | 8.9 | 73 970 015 | 98.7 | 95.9 |
| peacock023 | *Pavo cristatus* | cameo | black-shoulder | 14.1 | 119 938 068 | 99.1 | 95.8 |
| peacock024 | *Pavo cristatus* | cameo | | 9.2 | 79 105 044 | 98.8 | 96.8 |
| peacock025 | *Pavo cristatus* | cameo | | 11.8 | 104 125 601 | 99.1 | 96.1 |
| peacock026 | *Pavo cristatus* | cameo | | 8.3 | 71 172 758 | 98.9 | 96.8 |
| peacock027 | *Pavo cristatus* | cameo | | 10.5 | 93 198 763 | 98.8 | 96.2 |
| peacock028 | *Pavo cristatus* | opal | | 8.5 | 73 985 449 | 98.7 | 95.8 |
| peacock029 | *Pavo cristatus* | opal | | 9.0 | 78 284 880 | 98.9 | 95.7 |
| peacock030 | *Pavo cristatus* | opal | | 8.4 | 71 571 758 | 98.8 | 96.0 |
| peacock031 | *Pavo cristatus* | opal | | 8.8 | 73 891 382 | 98.4 | 95.8 |
| peacock032 | *Pavo cristatus* | opal | | 8.5 | 72 839 698 | 98.7 | 95.4 |
| peacock033 | *Pavo cristatus* | opal | | 6.8 | 59 094 643 | 99.1 | 95.7 |
| peacock034 | *Pavo cristatus* | opal | | 8.9 | 75 667 198 | 98.7 | 96.6 |
| peacock035 | *Pavo cristatus* | opal | | 7.8 | 71 572 780 | 98.7 | 95.8 |
| peacock036 | *Pavo cristatus* | opal | | 10.9 | 90 847 869 | 98.8 | 95.7 |
| peacock037 | *Pavo cristatus* | opal | | 10.0 | 85 219 019 | 99.0 | 96.1 |
| peacock038 | *Pavo cristatus* | opal | | 9.9 | 83 601 888 | 98.6 | 96.6 |
| peacock039 | *Pavo cristatus* | opal | | 8.0 | 69 961 722 | 98.2 | 95.1 |
| peacock040 | *Pavo cristatus* | opal | | 7.6 | 64 247 995 | 98.8 | 94.8 |
| peacock041 | *Pavo cristatus* | prussian-blue | black-shoulder | 16.9 | 148 632 494 | 99.0 | 96.3 |
| peacock042 | *Pavo cristatus* | prussian-blue | black-shoulder | 11.7 | 99 871 842 | 98.6 | 96.5 |
| peacock043 | *Pavo cristatus* | prussian-blue | black-shoulder | 7.9 | 68 903 920 | 98.8 | 95.5 |
| peacock044 | *Pavo cristatus* | prussian-blue | black-shoulder | 9.5 | 80 601 736 | 98.8 | 96.0 |
| peacock045 | *Pavo cristatus* | prussian-blue | black-shoulder | 9.7 | 82 980 871 | 98.6 | 96.2 |
| peacock046 | *Pavo cristatus* | prussian-blue | black-shoulder | 8.8 | 74 950 790 | 98.5 | 95.5 |
| peacock047 | *Pavo cristatus* | prussian-blue | black-shoulder | 8.9 | 78 423 035 | 98.7 | 95.3 |
| peacock048 | *Pavo cristatus* | prussian-blue | black-shoulder | 9.2 | 77 177 496 | 98.6 | 95.7 |
| peacock049 | *Pavo cristatus* | prussian-blue | black-shoulder | 14.2 | 128 846 524 | 97.8 | 95.8 |
| peacock050 | *Pavo cristatus* | prussian-blue | black-shoulder | 9.8 | 88 365 882 | 97.5 | 92.1 |
| peacock051 | *spalding* | purple | | 7.6 | 67 195 165 | 98.6 | 91.1 |
| peacock052 | *Pavo cristatus* | purple | | 6.6 | 58 007 239 | 98.8 | 95.8 |

| | | | | | | | |
|---|---|---|---|---|---|---|---|
| peacock053 | *Pavo cristatus* | purple | | 9.6 | 82 977 526 | 98.7 | 95.8 |
| peacock054 | *Pavo cristatus* | purple | | 5.2 | 46 716 968 | 98.6 | 95.8 |
| peacock055 | *Pavo cristatus* | purple | | 9.0 | 79 847 660 | 98.6 | 95.2 |
| peacock056 | *Pavo cristatus* | purple | | 4.9 | 42 516 372 | 98.3 | 95.2 |
| peacock057 | *Pavo cristatus* | purple | | 9.1 | 79 525 289 | 98.6 | 93.4 |
| peacock058 | *Pavo cristatus* | purple | | 7.5 | 64 188 415 | 98.7 | 94.9 |
| peacock059 | *Pavo cristatus* | purple | | 7.5 | 65 858 079 | 98.7 | 95.1 |
| peacock060 | *Pavo cristatus* | purple | | 7.4 | 65 375 482 | 98.6 | 95.2 |
| peacock061 | *Pavo cristatus* | purple | | 7.3 | 62 853 140 | 98.9 | 94.9 |
| peacock062 | *Pavo cristatus* | raw-umber | black-shoulder | 7.0 | 60 844 559 | 99.1 | 96.2 |
| peacock063 | *Pavo cristatus* | raw-umber | | 8.6 | 75 843 161 | 98.5 | 96.7 |
| peacock064 | *Pavo cristatus* | raw-umber | | 6.6 | 57 904 928 | 98.8 | 94.6 |
| peacock065 | *Pavo cristatus* | raw-umber | black-shoulder | 9.8 | 85 733 473 | 98.2 | 95.7 |
| peacock066 | *Pavo cristatus* | raw-umber | | 6.6 | 58 247 364 | 97.4 | 92.6 |
| peacock067 | *Pavo cristatus* | raw-umber | | 9.0 | 79 522 167 | 98.9 | 92.6 |
| peacock068 | *Pavo cristatus* | raw-umber | black-shoulder | 9.6 | 82 887 591 | 98.8 | 95.7 |
| peacock069 | *Pavo cristatus* | violet | | 9.7 | 86 060 745 | 98.5 | 96.2 |
| peacock070 | *Pavo cristatus* | violet | | 8.2 | 68 984 267 | 99.1 | 95.1 |
| peacock071 | *Pavo cristatus* | violet | | 6.7 | 59 488 393 | 98.9 | 97.0 |
| peacock072 | *Pavo cristatus* | violet | | 9.0 | 78 784 225 | 98.9 | 96.2 |
| peacock073 | *Pavo cristatus* | violet | | 8.4 | 73 061 934 | 98.7 | 95.9 |
| peacock074 | *Pavo cristatus* | violet | | 8.8 | 76 792 327 | 98.7 | 95.6 |
| peacock075 | *Pavo cristatus* | violet | | 7.3 | 64 151 498 | 98.8 | 95.5 |
| peacock076 | *Pavo cristatus* | violet | | 8.7 | 74 466 728 | 98.8 | 96.0 |
| peacock077 | *Pavo cristatus* | violet | | 8.1 | 71 459 385 | 98.6 | 95.8 |
| peacock078 | *Pavo cristatus* | violet | | 8.4 | 73 004 433 | 98.7 | 95.3 |
| peacock079 | *Pavo cristatus* | violet | | 10.9 | 96 074 477 | 98.7 | 95.3 |
| peacock080 | *Pavo cristatus* | violet | | 8.9 | 76 879 453 | 98.9 | 95.0 |
| peacock081 | *Pavo cristatus* | violet | | 9.3 | 80 503 388 | 98.5 | 96.2 |
| peacock082 | *Pavo cristatus* | white | | 7.2 | 61 219 168 | 98.9 | 95.0 |
| peacock083 | *Pavo cristatus* | white | | 8.3 | 70 906 180 | 98.5 | 96.2 |
| peacock084 | *Pavo cristatus* | white | | 8.9 | 78 109 981 | 98.3 | 95.3 |
| peacock085 | *Pavo cristatus* | white | | 8.5 | 76 331 840 | 98.8 | 94.5 |
| peacock086 | *Pavo cristatus* | white | | 7.9 | 67 077 443 | 98.8 | 95.8 |
| peacock087 | *Pavo cristatus* | white | | 8.1 | 69 013 029 | 98.0 | 95.8 |
| peacock088 | *Pavo cristatus* | white | | 7.0 | 61 258 164 | 98.5 | 93.5 |
| peacock089 | *Pavo cristatus* | white | | 6.0 | 52 758 580 | 98.7 | 95.1 |
| peacock090 | *Pavo cristatus* | white | | 7.3 | 64 010 239 | 98.9 | 95.8 |
| peacock091 | *Pavo cristatus* | white | | 8.0 | 69 844 607 | 98.6 | 96.4 |
| peacock092 | *Pavo cristatus* | wild-type | | 8.3 | 71 579 202 | 98.1 | 95.3 |
| peacock093 | *Pavo cristatus* | wild-type | | 8.8 | 75 518 209 | 98.9 | 93.9 |
| peacock094 | *Pavo cristatus* | wild-type | | 10.1 | 86 577 842 | 98.7 | 96.2 |
| peacock095 | *Pavo cristatus* | wild-type | | 9.1 | 77 748 705 | 98.4 | 95.1 |
| peacock096 | *Pavo cristatus* | wild-type | | 10.0 | 86 014 708 | 98.5 | 95.2 |
| peacock097 | *Pavo cristatus* | wild-type | | 10.2 | 91 156 190 | 99.1 | 94.8 |
| peacock098 | *Pavo cristatus* | wild-type | | 8.1 | 67 569 237 | 98.1 | 96.7 |
| peacock099 | *Pavo cristatus* | wild-type | | 6.9 | 58 771 766 | 98.9 | 94.5 |
| peacock100 | *Pavo cristatus* | wild-type | | 8.8 | 74 265 385 | 98.7 | 96.5 |
| peacock101 | *Pavo cristatus* | peach | | 9.4 | 79 710 570 | 98.5 | 95.9 |
| peacock102 | *Pavo cristatus* | peach | | 3.9 | 35 333 950 | 97.3 | 93.1 |
| peacock103 | *Pavo cristatus* | peach | | 7.5 | 63 475 572 | 98.2 | 85.8 |
| peacock104 | *Pavo cristatus* | brown | | 7.5 | 65 632 909 | 98.5 | 92.6 |
| peacock105 | *Pavo cristatus* | brown | | 6.0 | 51 289 165 | 97.9 | 93.0 |
| peacock106 | *Pavo cristatus* | brown | | 6.1 | 54 026 900 | 97.8 | 91.8 |
| peacock107 | *Pavo cristatus* | charcoal | | 5.8 | 51 348 493 | 98.1 | 89.7 |
| peacock108 | *Pavo cristatus* | charcoal | | 7.5 | 63 885 183 | 98.6 | 91.6 |

| | | | | | | | |
|---|---|---|---|---|---|---|---|
| peacock109 | *Pavo cristatus* | charcoal | | 9.7 | 83 648 618 | 98.1 | 93.5 |
| peacock110 | *Pavo cristatus* | charcoal | | 11.0 | 97 767 387 | 98.0 | 90.8 |
| peacock111 | *Pavo cristatus* | charcoal | | 8.3 | 73 001 564 | 98.0 | 91.6 |
| peacock112 | *Pavo cristatus* | midnight | black-shoulder | 7.1 | 61 585 754 | 98.4 | 91.4 |
| peacock113 | *Pavo cristatus* | midnight | black-shoulder | 5.4 | 47 093 664 | 98.1 | 92.5 |
| peacock114 | *Pavo cristatus* | midnight | black-shoulder | 5.5 | 48 336 350 | 97.6 | 91.7 |
| peacock115 | *Pavo cristatus* | wild-type | black-shoulder | 7.5 | 65 927 927 | 98.1 | 90.8 |
| peacock116 | *Pavo cristatus* | wild-type | black-shoulder | 10.6 | 91 047 910 | 98.5 | 91.1 |
| peacock117 | *Pavo cristatus* | wild-type | black-shoulder | 10.4 | 90 393 443 | 98.3 | 94.1 |
| peacock118 | *Pavo cristatus* | charcoal | | 2.6 | 48 042 179 | 98.9 | 92.3 |
| peacock119 | *Pavo cristatus* | hazel | | 1.9 | 44 767 165 | 99.2 | 96.8 |
| peacock120 | *Pavo cristatus* | indigo | | 3.4 | 49 989 461 | 99.3 | 97.6 |
| peacock121 | *Pavo cristatus* | ivory | | 3.2 | 60 497 633 | 99.1 | 97.9 |
| peacock122 | *Pavo cristatus* | jade | | 4.7 | 67 917 003 | 99.0 | 97.4 |
| peacock123 | *Pavo cristatus* | midnight | | 5.4 | 76 642 936 | 99.0 | 97.3 |
| peacock124 | *Pavo cristatus* | peach | | 2.0 | 28 200 742 | 99.0 | 97.1 |
| peacock125 | *Pavo cristatus* | opal | | 3.8 | 59 089 537 | 99.1 | 97.1 |
| peacock126 | *Pavo cristatus* | purple | | 3.1 | 53 963 210 | 99.1 | 97.5 |
| peacock127 | *Pavo cristatus* | cameo | | 3.4 | 53 314 729 | 99.2 | 97.6 |
| peacock128 | *Pavo cristatus* | sonja violet | | 3.4 | 54 987 361 | 99.0 | 97.6 |
| peacock129 | *Pavo cristatus* | white | | 2.8 | 47 046 279 | 99.2 | 97.4 |
| peacock130 | *Pavo cristatus* | white | | 3.5 | 48 956 062 | 99.3 | 97.9 |
| peacock132 | *Pavo cristatus* | white | | 4.8 | 86 794 650 | 99.2 | 98.0 |
| peacock133 | *Pavo cristatus* | white | | 2.9 | 49 086 529 | 98.9 | 97.8 |
| peacock134 | *Pavo cristatus* | white | | 4.9 | 68 266 639 | 99.2 | 96.9 |
| peacock135 | *Pavo cristatus* | white | | 5.5 | 101 212 335 | 98.8 | 97.7 |
| peacock136 | *Pavo cristatus* | white | | 5.6 | 188 093 363 | 99.0 | 96.7 |
| peacock137 | *Pavo cristatus* | opal | | 3.6 | 52 270 392 | 98.9 | 97.1 |
| peacock138 | *Spalding* | NA | | 2.6 | 46 333 090 | 99.0 | 96.9 |
| peacock139 | *Spalding* | NA | | 4.7 | 70 574 921 | 99.0 | 97.0 |
| peacock140 | *Spalding* | NA | | 6.9 | 108 033 826 | 99.0 | 96.9 |
| peacock141 | *Spalding* | NA | | 6.8 | 132 732 376 | 98.7 | 96.8 |
| peacock142 | *Spalding* | NA | | 6.0 | 103 739 490 | 98.7 | 96.0 |
| peacock143 | *Pavo cristatus* | charcoal | black-shoulder | 4.9 | 72 640 330 | 98.7 | 96.3 |
| peacock144 | *Spalding* | NA | | 5.0 | 67 949 451 | 99.1 | 96.5 |
| peacock145 | *Spalding* | NA | | 4.2 | 64 692 475 | 98.5 | 97.2 |
| peacock146 | *Spalding* | NA | | 1.4 | 13 994 504 | 99.3 | 95.9 |
| peacock147 | *Spalding* | NA | | 6.0 | 81 793 150 | 98.8 | 96.6 |
| peacock148 | *Pavo cristatus* | raw-umber | | 10.6 | 93 980 060 | 97.7 | 96.4 |
| peacock149 | *Pavo cristatus* | raw-umber | | 5.5 | 49 192 542 | 97.7 | 92.7 |
| peacock150 | *Pavo cristatus* | raw-umber | | 8.2 | 71 812 702 | 98.2 | 91.5 |
| peacock151 | *Pavo cristatus* | raw-umber | black-shoulder | 6.5 | 59 468 420 | 97.1 | 92.9 |
| peacock152 | *Pavo cristatus* | wild-type | | 9.1 | 83 614 418 | 97.5 | 85.6 |
| peacock153 | *Pavo cristatus* | raw-umber | black-shoulder | 17.9 | 161 977 719 | 97.9 | 90.4 |
| peacock154 | *Pavo cristatus* | raw-umber | black-shoulder | 7.7 | 67 938 857 | 98.4 | 92.9 |
| peacock155 | *Pavo cristatus* | prussian-blue | black-shoulder | 11.0 | 98 086 128 | 97.6 | 93.0 |
| peacock156 | *Pavo cristatus* | prussian-blue | black-shoulder | 8.6 | 76 077 469 | 98.1 | 89.9 |
| peacock157 | *Pavo cristatus* | prussian-blue | black-shoulder | 6.9 | 63 654 245 | 97.3 | 91.8 |
| peacock158 | *Pavo cristatus* | prussian-blue | black-shoulder | 7.8 | 70 272 783 | 97.3 | 89.6 |
| peacock159 | *Pavo cristatus* | prussian-blue | black-shoulder | 9.0 | 78 320 185 | 97.9 | 90.0 |
| peacock160 | *Pavo cristatus* | bronze | | 9.5 | 85 001 373 | 98.0 | 90.3 |
| peacock161 | *Pavo cristatus* | raw-umber/prussian-blue | black-shoulder | 10.2 | 92 448 410 | 98.0 | 92.6 |
| peacock162 | *Pavo cristatus* | raw-umber/prussian-blue | | 9.7 | 86 132 454 | 97.4 | 93.0 |
| peacock163 | *Pavo cristatus* | prussian-blue | black-shoulder | 12.8 | 115 611 222 | 98.2 | 91.3 |
| peacock164 | *Pavo cristatus* | wild-type | | 19.4 | 203 719 913 | 98.1 | 91.4 |
| peacock165 | *Pavo cristatus* | wild-type | | 7.6 | 68 939 995 | 97.8 | 93.6 |

| | | | | | | | |
|---|---|---|---|---|---|---|---|
| peacock166 | *Pavo cristatus* | wild-type | | 17.9 | 220 217 938 | 98.1 | 91.6 |
| peacock167 | *Pavo cristatus* | purple | | 22.1 | 258 065 356 | 98.2 | 93.9 |
| peacock168 | *Pavo cristatus* | wild-type | | 11.7 | 105 287 197 | 97.8 | 94.1 |
| peacock169 | *Pavo cristatus* | purple | black-shoulder | 7.7 | 138 550 110 | 98.9 | 92.3 |
| peacock170 | *Pavo cristatus* | purple | | 5.9 | 52 993 405 | 97.6 | 96.9 |
| peacock171 | *Pavo cristatus* | purple | | 18.8 | 216 601 888 | 98.3 | 91.4 |
| peacock172 | *Pavo cristatus* | purple | | 21.0 | 255 493 972 | 98.2 | 94.7 |
| peacock173 | *Pavo cristatus* | purple | | 14.6 | 136 616 219 | 96.7 | 94.2 |
| peacock174 | *Pavo cristatus* | bronze | black-shoulder | 3.4 | 29 753 754 | 98.2 | 85.4 |
| peacock175 | *Pavo cristatus* | bronze | black-shoulder | 10.3 | 89 029 840 | 97.9 | 84.0 |
| peacock178 | *Pavo cristatus* | hazel | black-shoulder | 3.3 | 29 990 348 | 98.3 | 86.6 |
| peacock179 | *Pavo cristatus* | hazel | black-shoulder | 7.3 | 62 493 554 | 98.2 | 86.1 |
| peacock180 | *Pavo cristatus* | taupe | | 4.8 | 41 541 203 | 98.4 | 86.8 |
| peacock181 | *Pavo cristatus* | taupe | | 3.9 | 34 575 680 | 98.2 | 88.8 |
| peacock182 | *Pavo cristatus* | taupe | | 4.9 | 42 864 124 | 98.4 | 84.2 |
| peacock183 | *Pavo cristatus* | taupe | | 6.8 | 57 857 064 | 98.7 | 87.6 |
| peacock184 | *Pavo cristatus* | bronze | | 4.3 | 37 787 983 | 98.4 | 89.4 |
| peacock185 | *Spalding* | NA | | 11.6 | 100 809 507 | 97.9 | 88.1 |
| peacock186 | *Spalding* | NA | | 7.2 | 63 201 159 | 98.5 | 84.4 |
| peacock188 | *Pavo cristatus* | cameo | | 6.3 | 53 966 104 | 98.1 | 87.9 |
| peacock189 | *Pavo cristatus* | cameo | | 6.8 | 59 474 996 | 98.1 | 87.0 |
| peacock190 | *Pavo cristatus* | cameo | | 2.9 | 24 800 490 | 98.2 | 85.9 |
| peacock191 | *Pavo cristatus* | cameo | | 6.1 | 53 233 914 | 98.6 | 86.8 |
| peacock194 | *Pavo cristatus* | hazel | | 1.1 | 9 818 940 | 98.4 | 88.6 |
| peacock195 | *Pavo cristatus* | taupe | | 2.0 | 17 654 757 | 98.2 | 86.6 |
| peacock196 | *Pavo cristatus* | bronze | black-shoulder | 1.2 | 10 988 511 | 98.3 | 83.6 |
| peacock197 | *Pavo cristatus* | purple | | 4.4 | 40 442 485 | 98.0 | 84.2 |
| peacock198 | *Pavo cristatus* | purple | | 7.5 | 63 134 995 | 98.4 | 79.3 |
| peacock199 | *Pavo cristatus* | purple | | 13.9 | 110 871 827 | 98.2 | 88.6 |
| peacock200 | *Pavo cristatus* | hazel | | 4.8 | 41 297 599 | 98.5 | 89.1 |
| peacock202 | *Pavo cristatus* | hazel | black-shoulder | 1.3 | 10 904 568 | 98.1 | 88.1 |
| peacock203 | *Pavo cristatus* | taupe | black-shoulder | 8.4 | 73 009 016 | 98.1 | 83.0 |
| peacock213 | *Pavo cristatus* | taupe | black-shoulder | 3.8 | 34 224 797 | 97.2 | 86.9 |
| peacock214 | *Pavo cristatus* | taupe | | 4.6 | 39 514 249 | 98.6 | 78.9 |
| peacock215 | *Pavo cristatus* | taupe | | 6.0 | 51 469 764 | 98.5 | 90.0 |
| peacock216 | *Pavo cristatus* | purple | | 8.1 | 70 110 130 | 98.3 | 88.3 |
| peacock217 | *Pavo cristatus* | purple | | 8.9 | 77 435 938 | 98.1 | 88.3 |
| peacock218 | *Pavo cristatus* | purple | | 13.4 | 113 232 769 | 98.1 | 85.9 |
| peacock219 | *Pavo cristatus* | purple | | 11.6 | 101 647 507 | 98.3 | 87.9 |
| peacock220 | *Pavo cristatus* | bronze | black-shoulder | 5.6 | 48 430 459 | 98.0 | 86.0 |
| peacock221 | *Pavo cristatus* | bronze | black-shoulder | 7.0 | 61 247 542 | 98.2 | 84.1 |
| peacock234 | *Pavo cristatus* | charcoal | black-shoulder | 2.2 | 18 964 784 | 97.5 | 86.1 |
| peacock235 | *Pavo cristatus* | charcoal | black-shoulder | 14.1 | 104 729 475 | 98.4 | 83.8 |
| peacock236 | *Pavo cristatus* | purple | | 12.9 | 106 671 473 | 97.8 | 93.6 |
| peacock237 | *Pavo cristatus* | wild-type | black-shoulder | 1.8 | 16 252 693 | 97.7 | 84.7 |
| peacock238 | *Pavo cristatus* | opal | black-shoulder | 1.3 | 11 967 643 | 97.5 | 82.3 |
| peacock239 | *Pavo cristatus* | opal | black-shoulder | 0.9 | 8 454 179 | 98.2 | 76.9 |
| peacock240 | *Pavo cristatus* | opal | | 3.0 | 27 155 082 | 97.1 | 83.3 |
| peacock241 | *Pavo cristatus* | opal | black-shoulder | 14.2 | 104 252 581 | 98.8 | 76.7 |
| peacock242 | *Pavo cristatus* | opal | black-shoulder | 14.2 | 105 820 728 | 98.4 | 95.6 |
| peacock243 | *Pavo cristatus* | opal | | 13.4 | 112 016 486 | 98.1 | 94.3 |
| peacock244 | *Pavo cristatus* | opal | | 12.3 | 108 830 935 | 98.2 | 88.4 |
| peacock245 | *Pavo cristatus* | cameo | | 7.8 | 67 308 883 | 98.2 | 87.6 |
| peacock246 | *Pavo cristatus* | bronze | black-shoulder | 8.6 | 76 081 418 | 98.1 | 88.3 |
| peacock247 | *Pavo cristatus* | wild-type | | 4.8 | 42 582 597 | 98.0 | 84.3 |
| peacock248 | *Pavo cristatus* | wild-type | | 2.4 | 21 660 929 | 98.0 | 82.8 |

| | | | | | | | |
|---|---|---|---|---|---|---|---|
| peacock249 | *Pavo cristatus* | opal | | 14.1 | 111 261 792 | 98.2 | 81.2 |
| peacock251 | *Pavo cristatus* | opal | | 11.2 | 97 602 424 | 97.7 | 89.2 |
| peacock252 | *Pavo cristatus* | purple | | 12.3 | 102 523 574 | 98.2 | 85.9 |
| peacock253 | *Pavo cristatus* | purple | | 12.6 | 109 400 811 | 98.2 | 89.5 |
| peacock255 | *Pavo cristatus* | midnight | black-shoulder | 13.5 | 110 539 650 | 98.2 | 87.7 |
| peacock265 | *Pavo cristatus* | charcoal | black-shoulder | 11.9 | 91 296 018 | 98.9 | 89.4 |
| peacock266 | *Pavo cristatus* | charcoal | black-shoulder | 6.1 | 45 863 079 | 99.1 | 96.1 |
| peacock267 | *Pavo cristatus* | charcoal | | 7.1 | 53 609 721 | 98.9 | 96.4 |
| peacock268 | *Pavo cristatus* | charcoal | | 7.7 | 59 159 845 | 99.0 | 96.0 |
| peacock269 | *Pavo cristatus* | charcoal | | 7.3 | 55 401 774 | 98.6 | 95.9 |
| peacock270 | *Pavo cristatus* | charcoal | | 3.2 | 24 650 448 | 98.4 | 94.7 |
| peacock271 | *Pavo cristatus* | charcoal | | 6.3 | 48 379 662 | 98.5 | 94.2 |
| peacock272 | *Pavo cristatus* | violet | | 3.4 | 25 635 955 | 98.7 | 94.1 |
| peacock273 | *Pavo cristatus* | violet | | 7.9 | 59 660 304 | 99.1 | 95.1 |
| peacock274 | *Pavo cristatus* | violet | | 3.9 | 29 666 064 | 98.5 | 96.2 |
| peacock275 | *Pavo cristatus* | violet | | 9.8 | 73 873 385 | 98.7 | 94.7 |
| peacock276 | *Pavo cristatus* | violet | | 10.2 | 77 304 191 | 98.9 | 95.5 |
| peacock277 | *Pavo cristatus* | violet | | 10.9 | 83 026 124 | 98.7 | 95.7 |
| peacock278 | *Pavo cristatus* | violet | | 12.5 | 94 625 445 | 99.0 | 95.3 |
| peacock279 | *Pavo cristatus* | violet | | 6.1 | 46 833 204 | 98.0 | 96.4 |
| peacock280 | *Pavo cristatus* | violet | | 8.6 | 65 470 491 | 98.2 | 93.1 |
| peacock281 | *Pavo cristatus* | violet | | 3.9 | 29 509 013 | 99.0 | 94.3 |
| peacock282 | *Pavo cristatus* | bronze | black-shoulder | 14.4 | 109 037 981 | 99.0 | 96.2 |
| peacock283 | *Pavo cristatus* | bronze | black-shoulder | 12.1 | 92 583 156 | 98.7 | 96.1 |
| peacock284 | *Pavo cristatus* | bronze | black-shoulder | 10.0 | 75 618 516 | 99.1 | 95.4 |
| peacock285 | *Pavo cristatus* | bronze | | 15.7 | 123 907 871 | 98.6 | 96.4 |
| peacock286 | *Pavo cristatus* | bronze | black-shoulder | 7.3 | 54 664 769 | 98.9 | 94.2 |
| peacock287 | *Pavo cristatus* | bronze | | 9.6 | 73 826 811 | 98.8 | 95.6 |
| peacock288 | *Pavo cristatus* | bronze | | 9.8 | 74 668 646 | 99.1 | 93.9 |
| peacock289 | *Pavo cristatus* | bronze | black-shoulder | 11.6 | 86 674 171 | 99.1 | 96.6 |
| peacock290 | *Pavo cristatus* | bronze | black-shoulder | 20.8 | 160 060 747 | 98.7 | 96.6 |
| peacock291 | *Pavo cristatus* | bronze | black-shoulder | 12.6 | 95 918 318 | 98.7 | 95.5 |
| peacock292 | *Pavo cristatus* | cameo | | 22.1 | 168 853 561 | 98.7 | 94.8 |
| peacock293 | *Pavo cristatus* | cameo | black-shoulder | 10.6 | 80 177 762 | 98.7 | 94.5 |
| peacock294 | *Pavo cristatus* | cameo | | 16.0 | 121 773 622 | 98.6 | 94.9 |
| peacock295 | *Pavo cristatus* | cameo | | 23.3 | 175 887 380 | 99.0 | 95.7 |
| peacock296 | *Pavo cristatus* | cameo | black-shoulder | 22.8 | 172 769 031 | 98.9 | 96.3 |
| peacock297 | *Pavo cristatus* | cameo | | 25.5 | 192 948 430 | 98.9 | 96.3 |
| peacock298 | *Pavo cristatus* | cameo | | 26.7 | 209 519 127 | 98.7 | 96.1 |
| peacock299 | *Pavo cristatus* | cameo | | 12.0 | 91 033 581 | 98.8 | 95.6 |
| peacock300 | *Pavo cristatus* | cameo | black-shoulder | 8.0 | 60 973 876 | 98.5 | 95.4 |
| peacock301 | *Pavo cristatus* | cameo | black-shoulder | 18.3 | 139 940 589 | 98.4 | 93.5 |
| peacock302 | *Pavo cristatus* | opal | black-shoulder | 19.9 | 152 210 258 | 98.7 | 94.2 |
| peacock303 | *Pavo cristatus* | opal | black-shoulder | 23.4 | 177 286 695 | 98.8 | 95.4 |
| peacock304 | *Pavo cristatus* | opal | | 26.9 | 203 808 073 | 99.1 | 95.7 |
| peacock305 | *Pavo cristatus* | opal | | 7.7 | 57 904 819 | 98.8 | 96.9 |
| peacock306 | *Pavo cristatus* | opal | | 21.7 | 164 997 536 | 99.0 | 96.0 |
| peacock307 | *Pavo cristatus* | opal | black-shoulder | 14.0 | 106 104 077 | 98.7 | 96.4 |
| peacock308 | *Pavo cristatus* | opal | | 21.1 | 160 762 662 | 98.7 | 95.7 |
| peacock309 | *Pavo cristatus* | opal | black-shoulder | 12.7 | 97 318 344 | 98.4 | 95.9 |
| peacock310 | *Pavo cristatus* | opal | | 11.7 | 89 630 025 | 98.7 | 94.4 |
| peacock311 | *Pavo cristatus* | opal | | 18.9 | 142 182 607 | 98.9 | 94.7 |
| peacock312 | *Pavo cristatus* | purple | black-shoulder | 5.5 | 40 748 769 | 99.1 | 96.2 |
| peacock313 | *Pavo cristatus* | purple | | 7.9 | 58 348 392 | 99.1 | 96.8 |
| peacock314 | *Pavo cristatus* | purple | | 14.0 | 105 088 511 | 99.2 | 96.9 |
| peacock315 | *Pavo cristatus* | purple | | 8.7 | 64 418 410 | 99.0 | 96.6 |

| | | | | | | | |
|---|---|---|---|---|---|---|---|
| peacock316 | *Pavo cristatus* | purple | | 5.2 | 38 640 189 | 99.0 | 96.3 |
| peacock317 | *Pavo cristatus* | purple | black-shoulder | 12.9 | 95 704 636 | 99.1 | 96.2 |
| peacock318 | *Pavo cristatus* | purple | | 5.9 | 43 641 533 | 98.8 | 96.9 |
| peacock319 | *Pavo cristatus* | purple | black-shoulder | 13.4 | 99 874 696 | 98.9 | 96.0 |
| peacock320 | *Pavo cristatus* | purple | | 21.8 | 163 038 760 | 99.2 | 96.3 |
| peacock321 | *Pavo cristatus* | purple | black-shoulder | 9.6 | 71 264 913 | 98.8 | 96.8 |
| peacock322 | *Pavo cristatus* | hazel | black-shoulder | 9.8 | 73 048 388 | 99.1 | 96.3 |
| peacock323 | *Pavo cristatus* | hazel | black-shoulder | 11.8 | 87 771 167 | 99.2 | 96.6 |
| peacock324 | *Pavo cristatus* | hazel | black-shoulder | 9.2 | 68 605 609 | 99.1 | 97.2 |
| peacock332 | *Pavo cristatus* | wild-type | black-shoulder | 14.1 | 105 321 345 | 99.0 | 96.6 |
| peacock333 | *Pavo cristatus* | wild-type | | 9.3 | 69 341 044 | 99.1 | 96.5 |
| peacock334 | *Pavo cristatus* | wild-type | black-shoulder | 9.5 | 71 184 976 | 99.0 | 96.9 |
| peacock335 | *Pavo cristatus* | wild-type | | 12.1 | 89 757 663 | 99.2 | 96.2 |
| peacock336 | *Pavo cristatus* | wild-type | | 9.1 | 67 797 867 | 99.0 | 97.0 |
| peacock337 | *Pavo cristatus* | wild-type | black-shoulder | 10.1 | 75 346 911 | 99.1 | 96.7 |
| peacock338 | *Pavo cristatus* | wild-type | black-shoulder | 5.9 | 43 460 812 | 99.0 | 96.7 |
| peacock339 | *Pavo cristatus* | hazel | | 6.7 | 49 856 723 | 99.1 | 96.4 |
| peacock340 | *Pavo cristatus* | chestnut | | 10.6 | 78 974 502 | 99.0 | 96.7 |
| peacock341 | *Pavo cristatus* | charcoal | | 16.9 | 126 937 880 | 98.9 | 96.3 |
| peacock342 | *Pavo muticus* | wild-type | | 9.2 | 68 532 071 | 98.8 | 96.4 |
| peacock343 | *Pavo muticus* | wild-type | | 22.2 | 166 010 149 | 98.9 | 95.6 |
| peacock350 | *Pavo cristatus* | wild-type | black-shoulder | 24.2 | 180 618 850 | 99.2 | 95.9 |
| peacock352 | *Pavo cristatus* | wild-type | | 10.6 | 78 836 727 | 98.9 | 97.1 |
| peacock358 | *Pavo muticus* | wild-type | | 29.0 | 221 634 574 | 98.9 | 96.3 |
| peacock359 | *Pavo muticus* | wild-type | | 17.0 | 127 349 189 | 98.7 | 95.9 |
| peacock364 | *Pavo cristatus* | violet | | 21.6 | 162 541 012 | 98.8 | 95.4 |
| peacock365 | *Pavo cristatus* | violet | | 12.6 | 94 525 743 | 98.8 | 95.7 |
| peacock366 | *Pavo cristatus* | violet | | 22.2 | 168 026 281 | 98.8 | 94.8 |
| peacock367 | *Pavo cristatus* | violet | | 21.6 | 161 679 458 | 99.2 | 94.7 |
| peacock368 | *Pavo cristatus* | purple | | 10.8 | 80 852 232 | 99.0 | 96.9 |
| peacock369 | *Pavo cristatus* | purple | black-shoulder | 16.1 | 119 825 424 | 99.2 | 96.6 |
| peacock370 | *Pavo cristatus* | purple | | 17.4 | 131 065 426 | 99.2 | 96.8 |
| peacock371 | *Pavo cristatus* | purple | | 8.6 | 64 012 797 | 99.0 | 96.7 |
| peacock376 | *Pavo cristatus* | opal | | 8.9 | 66 994 698 | 98.8 | 96.4 |
| peacock377 | *Pavo cristatus* | opal | black-shoulder | 7.5 | 56 583 466 | 98.8 | 94.5 |
| peacock378 | *Pavo cristatus* | opal | | 17.0 | 127 536 413 | 98.8 | 95.2 |
| peacock379 | *Pavo cristatus* | wild-type | | 22.8 | 170 104 871 | 99.1 | 93.8 |
| peacock380 | *Pavo cristatus* | wild-type | | 8.3 | 62 501 411 | 98.8 | 96.8 |
| peacock381 | *Pavo cristatus* | charcoal | | 6.0 | 44 327 572 | 99.2 | 95.9 |
| peacock383 | *Pavo cristatus* | cameo | black-shoulder | 10.3 | 77 230 425 | 98.9 | 96.5 |
| peacock384 | *Pavo cristatus* | cameo | | 5.3 | 40 070 238 | 98.4 | 96.0 |
| peacock385 | *Pavo cristatus* | cameo | | 11.9 | 88 952 118 | 98.8 | 93.2 |
| peacock386 | *Pavo cristatus* | bronze | | 9.9 | 74 748 054 | 98.9 | 94.3 |
| peacock387 | *Pavo cristatus* | bronze | | 12.5 | 93 672 049 | 98.9 | 95.2 |
| peacock388 | *Pavo cristatus* | bronze | black-shoulder | 13.6 | 105 025 164 | 98.6 | 95.4 |
| peacock389 | *Pavo cristatus* | midnight | black-shoulder | 10.3 | 77 968 699 | 98.9 | 95.2 |
| peacock390 | *Pavo cristatus* | midnight | black-shoulder | 7.4 | 56 142 606 | 98.6 | 96.3 |
| peacock391 | *Pavo cristatus* | midnight | | 12.5 | 95 478 844 | 98.9 | 95.4 |
| peacock395 | *Pavo cristatus* | wild-type | black-shoulder | 12.7 | 97 124 037 | 98.5 | 96.0 |
| peacock398 | *Pavo cristatus* | opal | black-shoulder | 8.6 | 64 820 694 | 98.9 | 95.3 |
| peacock399 | *Pavo cristatus* | opal | black-shoulder | 8.8 | 67 110 737 | 98.8 | 96.1 |
| peacock400 | *Pavo cristatus* | opal | | 14.8 | 111 879 169 | 98.7 | 95.9 |
| peacock401 | *Pavo cristatus* | taupe | | 11.7 | 88 998 685 | 98.7 | 95.6 |
| peacock402 | *Pavo cristatus* | taupe | | 14.5 | 109 898 461 | 98.9 | 95.8 |
| peacock403 | *Pavo cristatus* | charcoal | | 5.7 | 42 462 794 | 98.9 | 96.1 |
| peacock408 | *Pavo cristatus* | midnight | black-shoulder | 9.4 | 70 858 951 | 98.9 | 96.0 |

| | | | | | | | |
|---|---|---|---|---|---|---|---|
| peacock409 | *Pavo cristatus* | midnight | black-shoulder | 13.4 | 102 506 553 | 98.7 | 95.8 |
| peacock410 | *Pavo cristatus* | wild-type | black-shoulder | 18.6 | 141 930 675 | 98.8 | 95.7 |
| peacock411 | *Pavo cristatus* | wild-type | | 23.0 | 176 741 713 | 98.4 | 95.6 |
| peacock412 | *Pavo cristatus* | wild-type | | 13.1 | 99 678 354 | 98.9 | 93.9 |
| peacock413 | *Pavo cristatus* | wild-type | | 9.2 | 70 188 099 | 98.6 | 95.9 |
| peacock414 | *Pavo cristatus* | wild-type | | 17.8 | 134 808 622 | 98.7 | 95.0 |
| peacock415 | *Pavo cristatus* | wild-type | | 10.5 | 79 961 235 | 98.8 | 95.4 |
| peacock416 | *Pavo cristatus* | wild-type | | 10.0 | 78 318 965 | 98.7 | 95.8 |
| peacock417 | *Pavo cristatus* | wild-type | | 6.9 | 52 279 229 | 98.6 | 95.2 |
| peacock418 | *Pavo cristatus* | wild-type | | 9.2 | 71 429 623 | 98.4 | 94.9 |
| peacock421 | *Pavo cristatus* | wild-type | | 20.6 | 168 551 380 | 98.4 | 94.8 |
| peacock422 | *Pavo cristatus* | wild-type | | 18.6 | 153 383 463 | 98.7 | 96.0 |
| peacock423 | *Pavo cristatus* | wild-type | | 18.5 | 150 156 609 | 98.6 | 96.3 |
| peacock424 | *Pavo cristatus* | wild-type | | 22.9 | 189 274 044 | 98.5 | 96.4 |
| peacock425 | *Pavo cristatus* | wild-type | | 21.8 | 177 563 064 | 98.7 | 96.5 |
| peacock426 | *Pavo cristatus* | wild-type | | 14.5 | 118 026 139 | 98.7 | 96.8 |
| peacock428 | *Pavo cristatus* | charcoal | | 9.0 | 117 628 788 | 99.6 | 96.7 |
| peacock429 | *Pavo cristatus* | ultramarine | | 9.9 | 149 783 723 | 99.7 | 98.5 |
| peacock430 | *Pavo cristatus* | ultramarine | | 13.0 | 179 668 317 | 99.7 | 98.8 |
| peacock431 | *Pavo cristatus* | ultramarine | | 5.9 | 68 325 260 | 99.6 | 98.9 |
| peacock432 | *Pavo cristatus* | ultramarine | | 15.9 | 274 466 223 | 99.3 | 98.7 |
| peacock433 | *Pavo cristatus* | ultramarine | | 14.8 | 210 587 599 | 99.7 | 98.2 |
| peacock434 | *Pavo cristatus* | ultramarine | | 12.5 | 175 168 642 | 98.0 | 97.5 |
| peacock435 | *Pavo cristatus* | ultramarine | | 18.1 | 301 514 174 | 99.3 | 97.3 |
| peacock436 | *Pavo cristatus* | ultramarine | | 17.7 | 222 126 642 | 99.7 | 98.0 |
| peacock437 | *Pavo cristatus* | ultramarine | | 18.2 | 248 213 883 | 99.8 | 97.3 |
| peacock438 | *Pavo cristatus* | ultramarine | | 4.3 | 61 006 029 | 99.6 | 96.5 |
| peacock439 | *Pavo cristatus* | ultramarine | | 5.6 | 69 465 446 | 98.3 | 98.5 |
| peacock448 | *Pavo cristatus* | raw-umber | black-shoulder | 12.0 | 99 918 843 | 98.5 | 96.1 |
| peacock449 | *Pavo cristatus* | raw-umber | black-shoulder | 14.2 | 117 310 064 | 98.5 | 96.2 |
| peacock452 | *Pavo cristatus* | raw-umber/ prussian-blue | black-shoulder | 18.8 | 153 387 245 | 98.7 | 96.1 |
| peacock453 | *Pavo cristatus* | prussian-blue | black-shoulder | 25.6 | 211 854 433 | 98.4 | 96.7 |
| peacock454 | *Pavo cristatus* | prussian-blue | black-shoulder | 19.2 | 157 758 481 | 98.7 | 96.4 |
| peacock455 | *Pavo cristatus* | prussian-blue | black-shoulder | 20.3 | 168 577 890 | 98.3 | 97.0 |
| peacock456 | *Pavo cristatus* | prussian-blue | black-shoulder | 18.9 | 153 790 587 | 98.9 | 95.8 |
| peacock457 | *Pavo cristatus* | prussian-blue | black-shoulder | 23.0 | 190 959 524 | 98.7 | 97.0 |

**Table S8. RNA-sequencing statistics for regenerating feather follicles from peafowl neck and wing.**

| ID | Body phenotype | Wing phenotype | Feather follicle | | Number of reads |
|---|---|---|---|---|---|
| | | | Neck | Wing | |
| peacock460 | Opal | | X | | 84 663 672 |
| peacock469 | Cameo | | X | | 97 612 226 |
| peacock470 | Cameo | | X | | 97 612 226 |
| peacock471 | Bronze | | X | | 84 853 934 |
| peacock473 | Wild-type | | X | | 67 658 498 |
| peacock474 | Bronze | | X | | 54 789 354 |
| peacock475 | Bronze | | X | | 82 021 652 |
| peacock476 | Violet | | X | | 82 765 102 |
| peacock477 | Violet | | X | | 83 352 310 |
| peacock479 | Wild-type | | X | | 82 127 766 |
| peacock480 | White | | X | | 47 051 030 |
| peacock487 | Purple | | X | | 68 502 820 |
| peacock488 | Purple | | X | | 83 074 342 |
| peacock489 | Wild-type | | X | | 85 734 468 |
| peacock490 | Wild-type | | X | | 71 770 966 |
| peacock491 | Opal | | X | | 77 832 716 |
| peacock492 | Cameo | | X | | 62 452 472 |
| peacock493 | Purple | | X | | 56 326 838 |
| peacock494 | Wild-type | | X | | 63 742 562 |
| peacock496 | White | | X | | 88 610 120 |
| peacock497 | Wild-type | | X | | 65 887 202 |
| peacock506 | Ultramarine | | X | | 145 574 546 |
| peacock507 | *Ultramarine* | | X | | 194 918 934 |
| peacock508 | *Charcoal* | | X | | 202 059 768 |
| peacock510 | *Ultramarine* | | X | | 206 821 926 |
| peacock521 | *Charcoal* | | X | | 210 442 370 |
| peacock522 | *Charcoal* | | X | | 211 975 220 |
| peacock473 | | Wild-type | | X | 66 416 662 |
| peacock476 | | Wild-type | | X | 51 536 456 |
| peacock489 | | Black-shoulder | | X | 63 387 960 |
| peacock497 | | Wild-type | | X | 62 204 698 |
| peacock500 | | Wild-type | | X | 41 911 332 |
| peacock501 | | Black-shoulder | | X | 56 572 500 |
| peacock502 | | Black-shoulder | | X | 53 733 978 |
| peacock503 | | Black-shoulder | | X | 44 944 076 |
| peacock504 | | Wild-type | | X | 73 628 562 |
| peacock505 | | Wild-type | | X | 66 995 070 |

**Table S9. Quantitative analysis of melanin content in feathers of the common emerald dove and mallard**

| | | | S-350 (/mg)[b] | | AHPO (ng/mg)[c] | | | HI (ng/mg)[d] | | TM[e] | EM[f] | BZ[g] | BT[h] | PM[i] |
|---|---|---|---|---|---|---|---|---|---|---|---|---|---|---|
| **Species** | **Sample ID** | **Area[a]** | **A500** | **A650** | **PTCA** | **PDCA** | **TTCA** | **4-AHP** | **3-AHP** | **(μg/mg)** | **(μg/mg)** | **(μg/mg)** | **(μg/mg)** | **(μg/mg)** |
| Mallard | 1 | i | 1.324 | 0.452 | 1337 | 127 | 578 | 51 | 36 | 134 | 107 | 19.7 | 0.5 | 20.1 |
| Mallard | 1 | ii | 0.297 | 0.099 | 398 | 38 | 120 | 7 | 11 | 30 | 32 | 4.1 | 0.1 | 4.1 |
| Mallard | 2 | i | 1.246 | 0.429 | 1274 | 122 | 577 | 59 | 8 | 126 | 102 | 19.6 | 0.5 | 20.1 |
| Mallard | 2 | ii | 0.271 | 0.091 | 341 | 28 | 123 | 6 | 10 | 27 | 27 | 4.2 | 0.1 | 4.2 |
| Mallard | 3 | i | 1.360 | 0.462 | 1162 | 80 | 551 | 81 | 37 | 137 | 93 | 18.7 | 0.7 | 19.5 |
| Mallard | 3 | ii | 0.336 | 0.116 | 456 | 39 | 116 | 8 | 7 | 34 | 36 | 3.9 | 0.1 | 4.0 |
| Mallard | 4 | i | 1.430 | 0.488 | 1182 | 80 | 565 | 71 | 44 | 144 | 95 | 19.2 | 0.6 | 19.8 |
| Mallard | 4 | ii | 0.424 | 0.136 | 513 | 44 | 179 | 8 | 9 | 43 | 41 | 6.1 | 0.1 | 6.2 |
| Emerald dove | 1 | i | 2.377 | 0.839 | 2409 | 204 | 660 | 47 | 33 | 240 | 193 | 22.4 | 0.4 | 22.9 |
| Emerald dove | 1 | ii | 1.276 | 0.433 | 1633 | 143 | 184 | 13 | 31 | 129 | 131 | 6.3 | 0.1 | 6.4 |
| Emerald dove | 2 | i | 2.405 | 0.841 | 2429 | 224 | 639 | 40 | 51 | 243 | 194 | 21.7 | 0.4 | 22.1 |
| Emerald dove | 2 | ii | 1.405 | 0.464 | 1691 | 151 | 188 | 17 | 47 | 142 | 135 | 6.4 | 0.1 | 6.5 |

[a]The number of the area refers to **fig. S11**.
[b]Soluene-350 solubilization
[c]Alkaline hydrogen peroxide oxidation
[d]Hydroiodic acid hydrolysis
[e]Total melanin (A500 × 101)
[f]Eumelanin (PTCA × 0.080)
[g]Benzothiazole (TTCA × 0.034)
[h]Benzothiazine (4-AHP × 0.009)
[i]Pheomelanin (BZ + BT)

**Table S10. Single-cell RNA-sequencing and read mapping statistics.**

| Species | Individual | Experiment | Phenotype | Number of read pairs | Valid barcodes (%)[a] | Mapped reads (%)[b] | Reads mapped confidently to genome (%)[c] | Reads mapped confidently to transcriptome (%)[d] | Reads mapped confidently to intronic regions (%)[e] | Reads mapped confidently to exonic regions (%)[f] | Reads mapped antisense to gene (%)[g] | Valid UMI (%)[h] |
|---|---|---|---|---|---|---|---|---|---|---|---|---|
| Emerald dove | 1 | 1 | Iridescent | 77,432,152 | 96.6 | 89.9 | 81.3 | 68.3 | 12.4 | 60.0 | 4.0 | 99.9 |
| Emerald dove | 1 | 1 | Non-iridescent | 119,961,948 | 94.4 | 72.1 | 63.5 | 50.8 | 12.4 | 43.6 | 5.2 | 100.0 |
| Emerald dove | 1 | 2 | Iridescent | 212,496,461 | 96.0 | 94.1 | 86.0 | 72.7 | 13.2 | 63.2 | 3.6 | 100.0 |
| Emerald dove | 1 | 2 | Non-iridescent | 87,166,635 | 96.1 | 93.8 | 82.4 | 69.2 | 15.4 | 58.2 | 4.2 | 99.6 |
| Mallard | 1 | 3 | Iridescent | 132,157,734 | 97.4 | 89.9 | 88.6 | 77.7 | 18.1 | 64.9 | 5.2 | 99.7 |
| Mallard | 1 | 3 | Non-iridescent | 119,568,782 | 97.2 | 90.5 | 88.8 | 76.8 | 16.1 | 67.2 | 6.4 | 99.7 |
| Mallard | 2 | 4 | Iridescent | 96,545,734 | 96.3 | 90.7 | 89.7 | 78.5 | 21.8 | 62.6 | 5.8 | 99.8 |
| Mallard | 2 | 4 | Non-iridescent | 104,256,954 | 95.8 | 90.5 | 89.6 | 78.2 | 23.0 | 61.1 | 5.7 | 99.8 |

[a]Fraction of reads with barcodes that match the whitelist after barcode correction
[b]Fraction of reads that mapped to the genome
[c]Fraction of reads that mapped uniquely to the genome
[d]Fraction of reads that mapped to a unique gene in the transcriptome
[e]Fraction of reads that mapped uniquely to an intronic region of the genome
[f]Fraction of reads that mapped uniquely to an exonic region of the genome
[g]Fraction of reads confidently mapped to the transcriptome on the opposite strand of their annotated gene
[h]Fraction of reads with UMI sequences that do not contain Ns and that are not homopolymers

**Table S11. Single-cell RNA-sequencing tissue composition and cell proportion differences between iridescent and non-iridescent feather halves in the common emerald dove and mallard.**

| | **Emerald dove** | | | | | | |
|---|---|---|---|---|---|---|---|
| | **Iridescent color** | | **Non-iridescent color** | | **Two Sample t-test** | | |
| | Replicate 1 | Replicate 2 | Replicate 1 | Replicate 2 | *t* | *df* | *P* |
| Keratinocytes | 1311 | 1247 | 881 | 822 | -0.159 | 1.849 | 0.890 |
| Melanocytes 1 | 545 | 641 | 125 | 119 | 12.955 | 1.288 | **0.025** |
| Melanocytes 2 | 227 | 236 | 19 | 84 | 3.566 | 1.408 | 0.115 |
| Endothelial cells | 236 | 413 | 228 | 338 | -1.762 | 1.506 | 0.259 |
| Mesenchymal 1 | 258 | 545 | 329 | 584 | -1.975 | 1.998 | 0.187 |
| Mesenchymal 2 | 35 | 68 | 86 | 87 | -1.337 | 1.343 | 0.364 |
| Leukocytes 1 | 94 | 130 | 33 | 116 | 0.123 | 1.025 | 0.922 |
| Leukocytes 2 | 54 | 84 | 20 | 81 | -0.123 | 1.095 | 0.921 |
| Leukocytes 3 | 15 | 23 | 16 | 23 | -4.148 | 1.848 | 0.061 |
| Erythrocytes | 7 | 6 | 15 | 3 | -0.770 | 1.022 | 0.580 |
| Neural-crest derived cells | 5 | 10 | 3 | 11 | -0.547 | 1.261 | 0.666 |
| All | 2787 | 3403 | 1755 | 2268 | | | |

| | **Mallard** | | | | | | |
|---|---|---|---|---|---|---|---|
| | **Iridescent color** | | **Non-iridescent color** | | **Two Sample t-test** | | |
| | Replicate 1 | Replicate 2 | Replicate 1 | Replicate 2 | *t* | *df* | *P* |
| Keratinocytes | 2797 | 4047 | 4228 | 3240 | -3.262 | 1.784 | 0.096 |
| Mesenchymal 1 | 1899 | 1425 | 413 | 639 | 2.182 | 1.339 | 0.220 |
| Melanocytes 1 | 74 | 601 | 94 | 86 | 0.874 | 1.000 | 0.543 |
| Melanocytes 2 | 74 | 176 | 38 | 46 | 1.802 | 1.107 | 0.304 |
| Leukocytes | 147 | 143 | 43 | 158 | 0.200 | 1.119 | 0.871 |
| Endothelial cells | 86 | 80 | 105 | 135 | -2.133 | 1.402 | 0.218 |
| Mesenchymal 2 | 42 | 29 | 33 | 46 | -0.842 | 1.996 | 0.489 |
| Erythrocytes | 12 | 26 | 17 | 12 | 0.085 | 1.322 | 0.943 |
| All | 5131 | 6527 | 4971 | 4362 | | | |

### SUPPLEMENTAL FIGURES

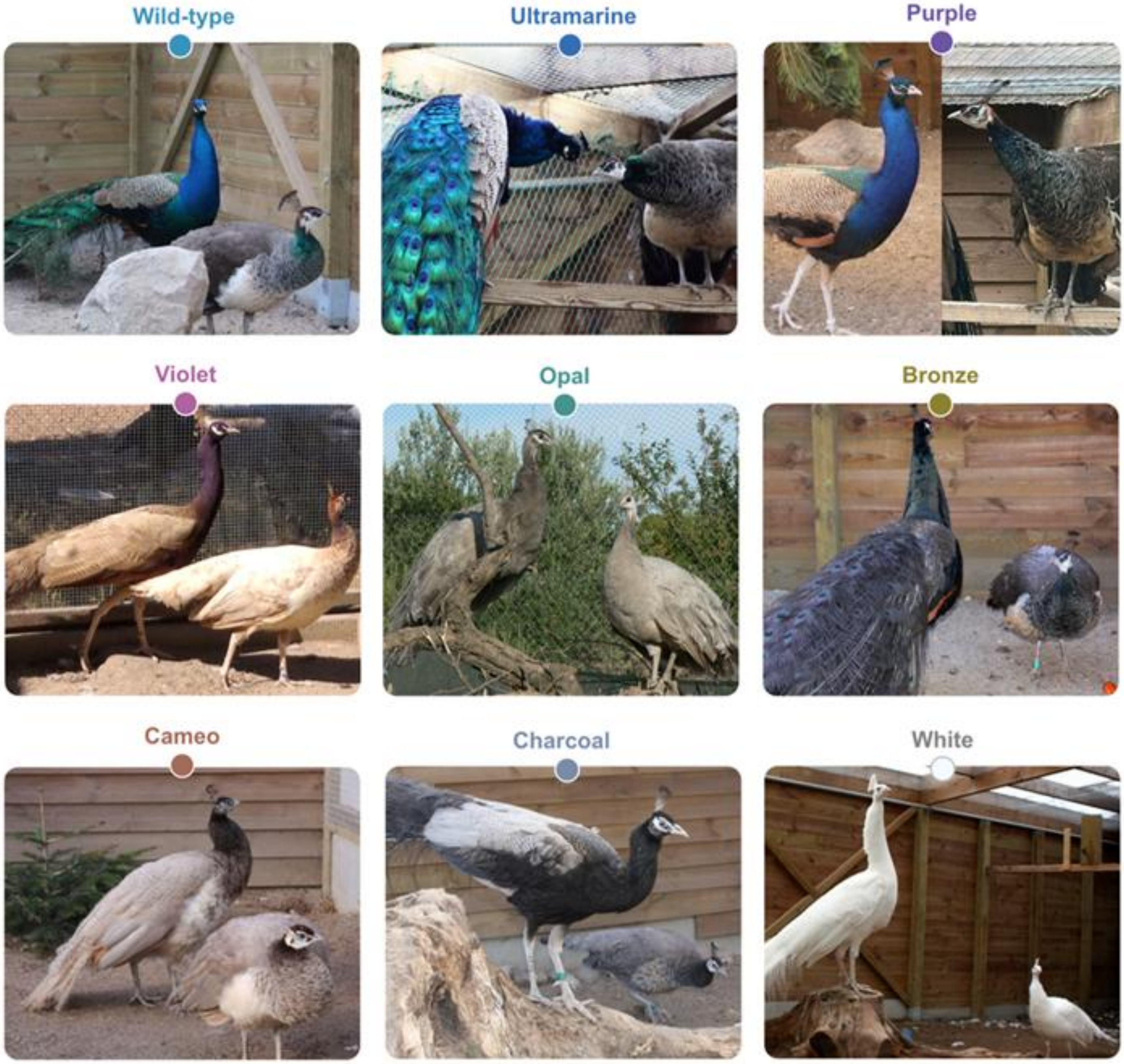


**Figure S1.** Phenotypic variation in male peafowl (peacocks, left) and female peafowl (peahens, right) across the mutations examined in this study. Each mutant phenotype differs by a single mutational step from the wild-type.

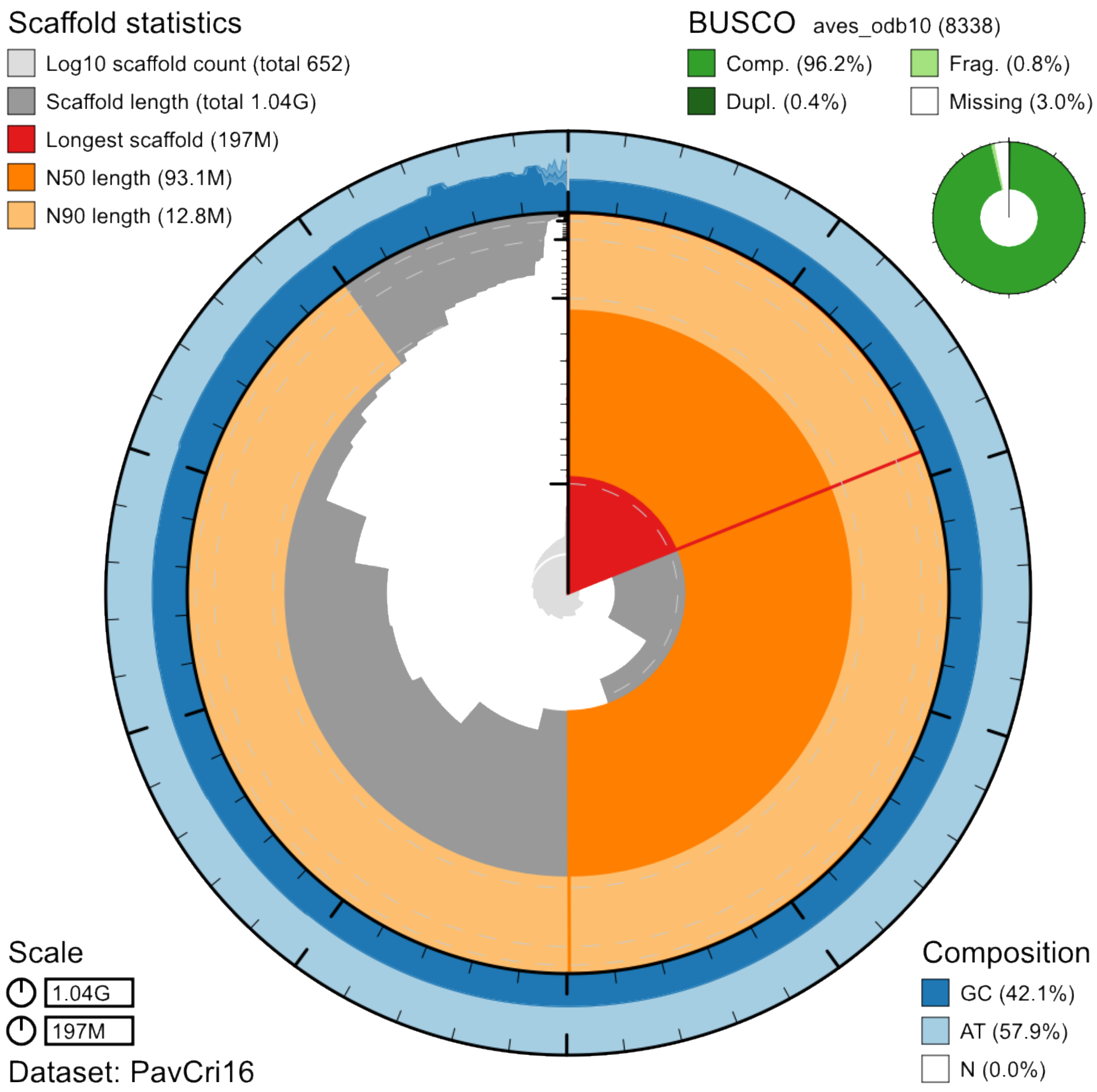


**Figure S2. Snail plot summary of assembly statistics for the Indian peafowl (*Pavo cristatus*) reference genome (*PavCri17*).** This plot provides an overview of the *PavCri17* genome assembly statistics (see also **table S1**). The main circular plot is divided into 1,000 size-ordered bins, each representing 0.1% of the 1,036,683,990 bp assembly. The distribution of sequence lengths is depicted in dark gray, with the plot radius scaled to the longest scaffold (196,737,001 bp, shown in red). Orange and pale-orange arcs indicate the N50 (93,074,296 bp) and N90 (12,795,000 bp) sequence lengths, respectively. The pale gray spiral represents the cumulative sequence count on a logarithmic scale, with white dashed lines marking increasing orders of magnitude. The outer blue and pale-blue regions show GC, AT, and N content distributions in the same bins as the inner plot. A summary of complete, fragmented, duplicated, and missing BUSCO genes from the *aves_odb10* set is provided in the top right corner.

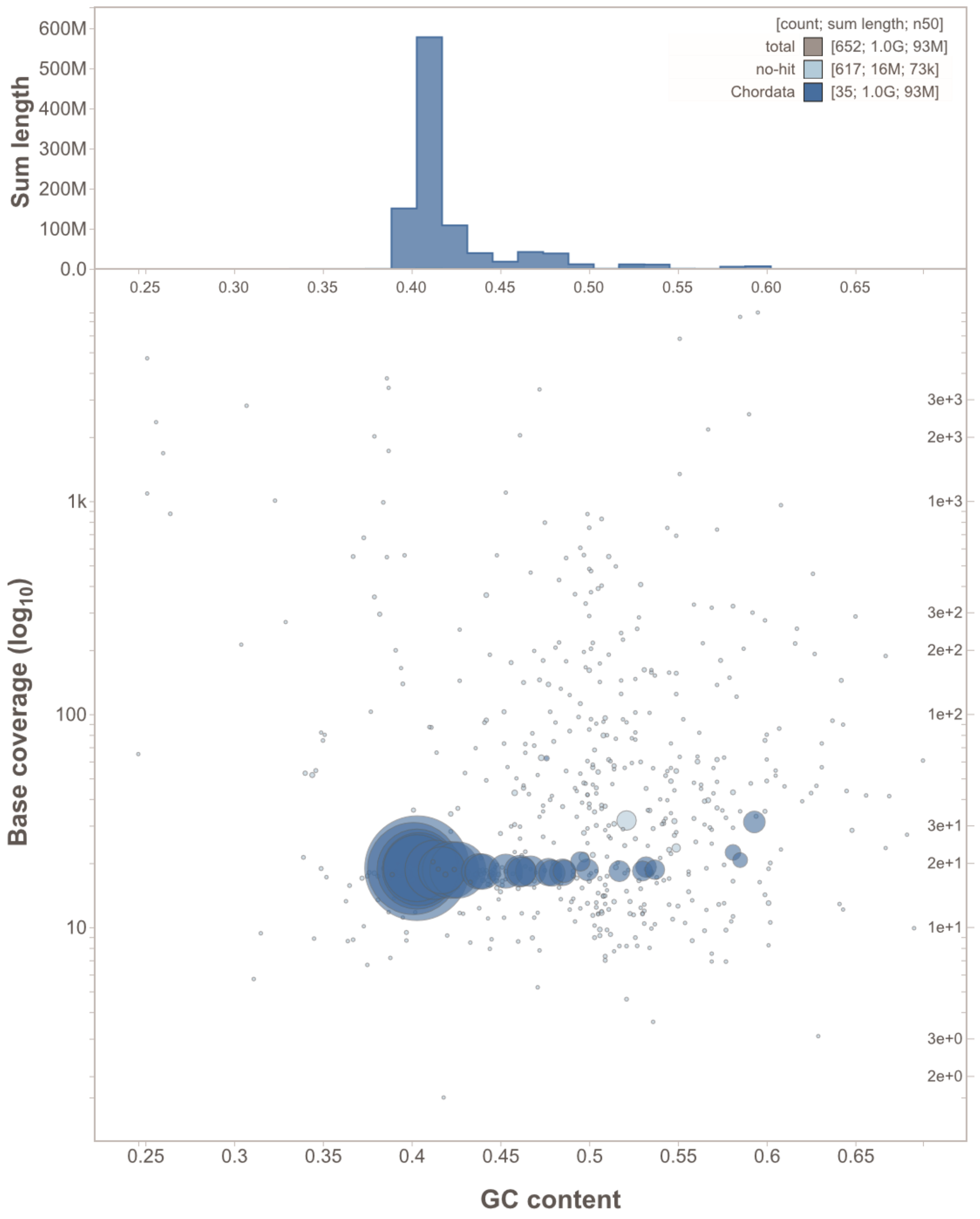


**Figure S3. Blob Plot**. The plot displays base coverage (Illumina reads; *y*-axis) versus GC content (*x*-axis) for scaffolds in the Indian peafowl (*Pavo cristatus*) reference genome (*PavCri17*). Circle sizes correspond to scaffold lengths. Marginal histograms illustrate the cumulative sequence length distribution along each axis. All scaffolds were assigned to Chordata (top-right corner), and coverage remained consistent across major scaffolds, providing no evidence of significant contamination from other taxa.

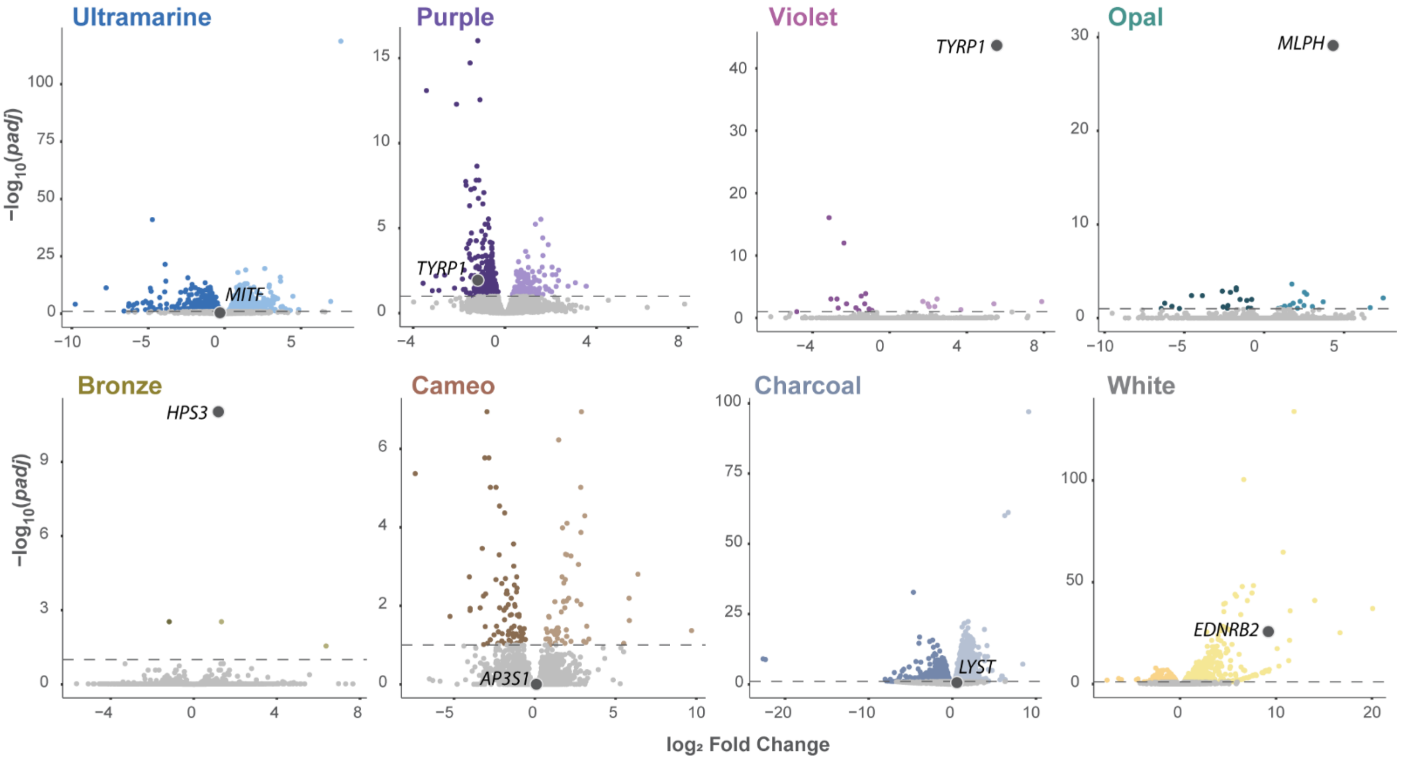


**Figure S4. Volcano plots of RNA sequencing results**. Each circle represents one gene, and light and dark colors indicate significant under- and over-expression in the mutant, respectively. Genes highlighted and labeled in dark gray circles represent those for which a genomic association was found.

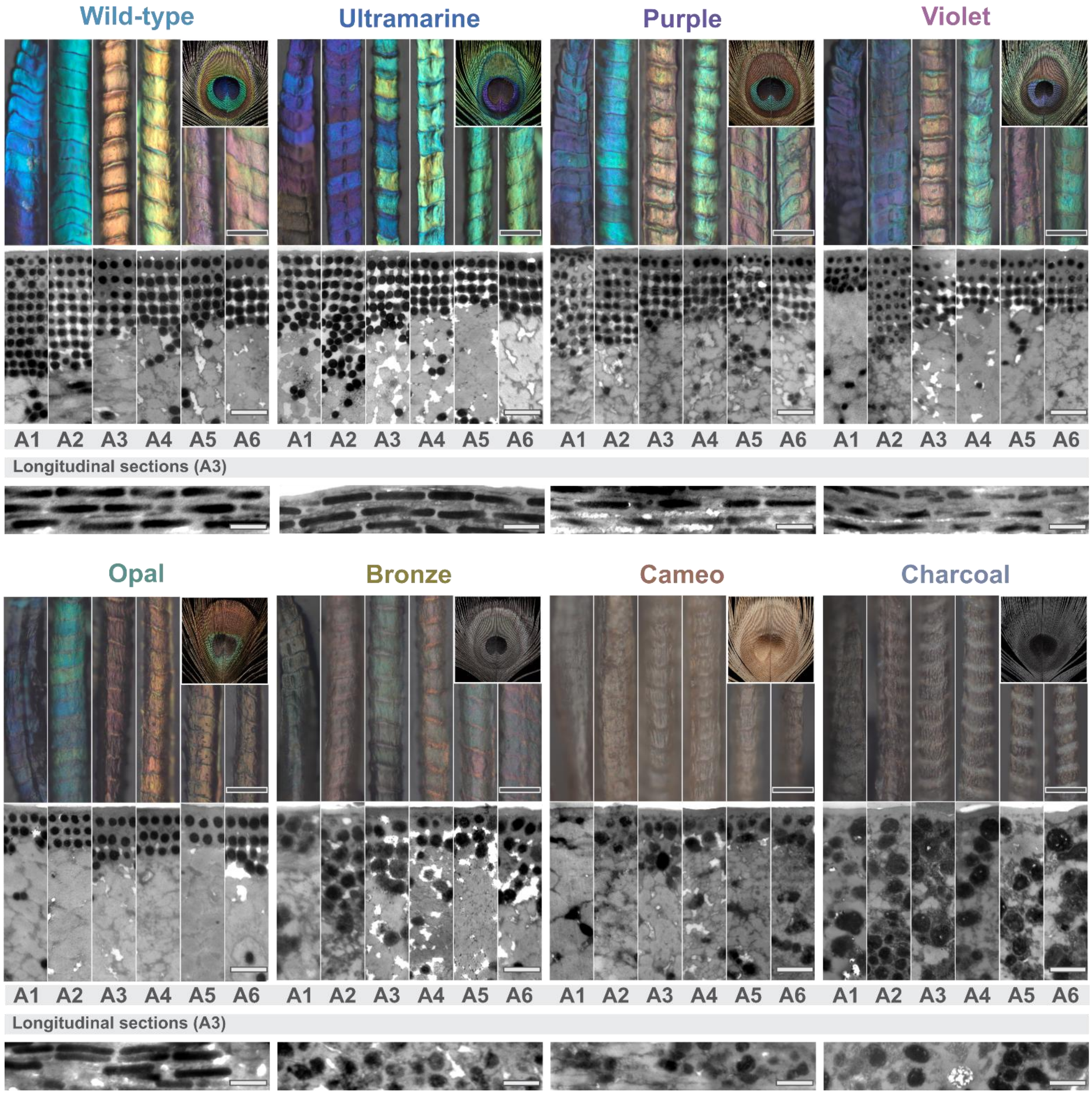


**Figure S5. Optical and transmission electron microscopy (TEM) images of feather barbules.** Transversal and longitudinal cuts are shown for each of the six areas of the eyespot (A1–A6) in wild-type and seven mutants. For each mutant and area, we show a picture of the feather barbule on top and TEM below, with an inset of the macroscopic appearance of the eyespot in the top right corner. Across the bottom of each set, we show a TEM longitudinal section of A3 to exemplify melanosome shape as rod-like versus sphere-like. Gray and white scales represent 50 µm (full barbule) and 500 nm (barbule sections), respectively.

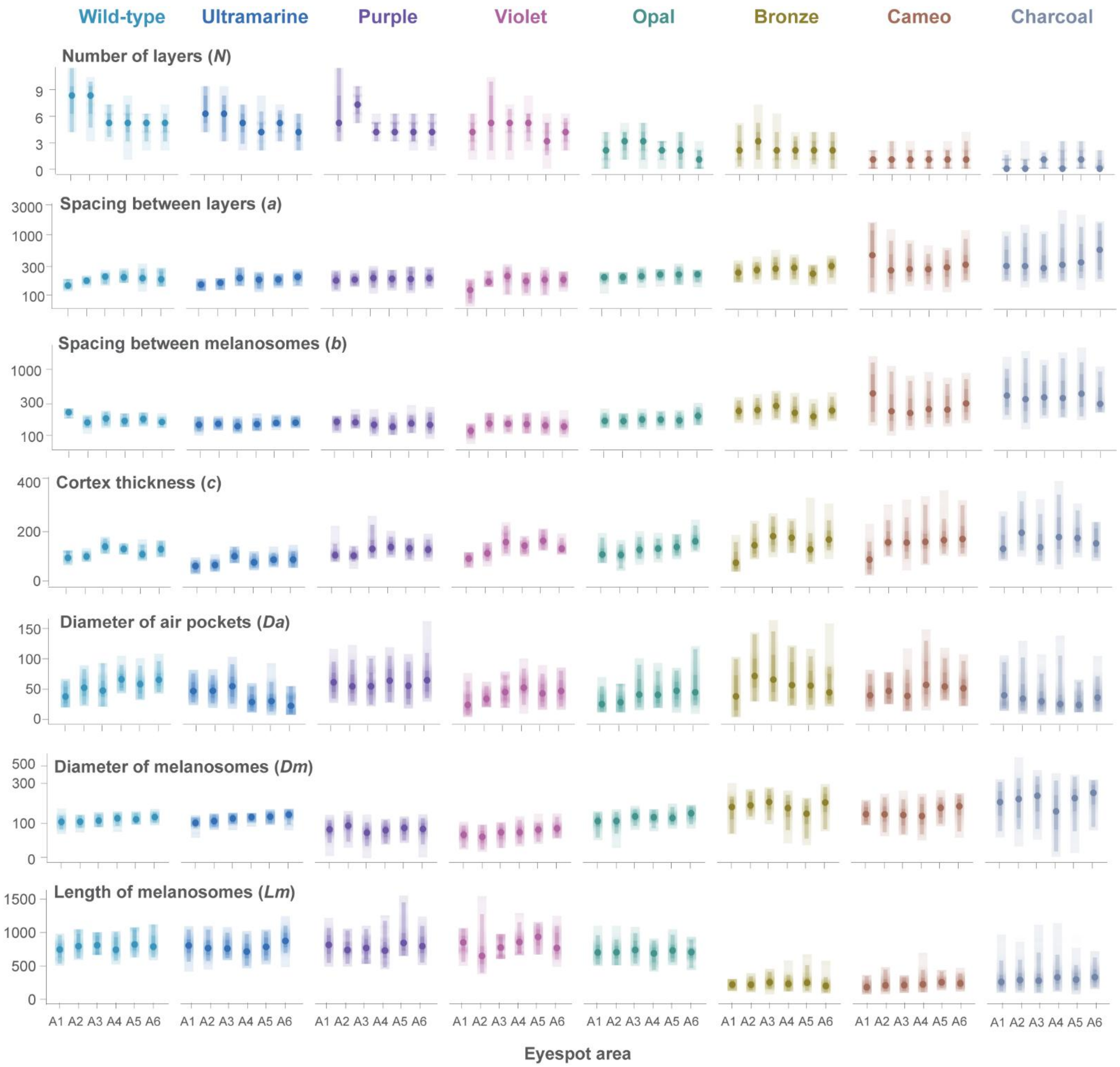


**Figure S6. TEM measurements of photonic lattice parameters.** Gradient plots of variation in lattice parameters across A1–A6 regions for wild-type and peafowl mutants: number of melanosome layers (*N*), spacing between layers (*a*), spacing between melanosomes within layer (*b*), cortex thickness (*c*), diameter of air pockets (*Da*), diameter of melanosomes (*Dm*), and length of melanosomes (*Lm*). Each circle represents an average measurement for a specific region, narrow bars represent quantile intervals at the 80% and 95% levels, and the wider bar represents the minimum–maximum range for each area. Spacing measures are in nm, and a log scale is used for clearer visualization of the variation in *a*, *b,* and *Dm*.

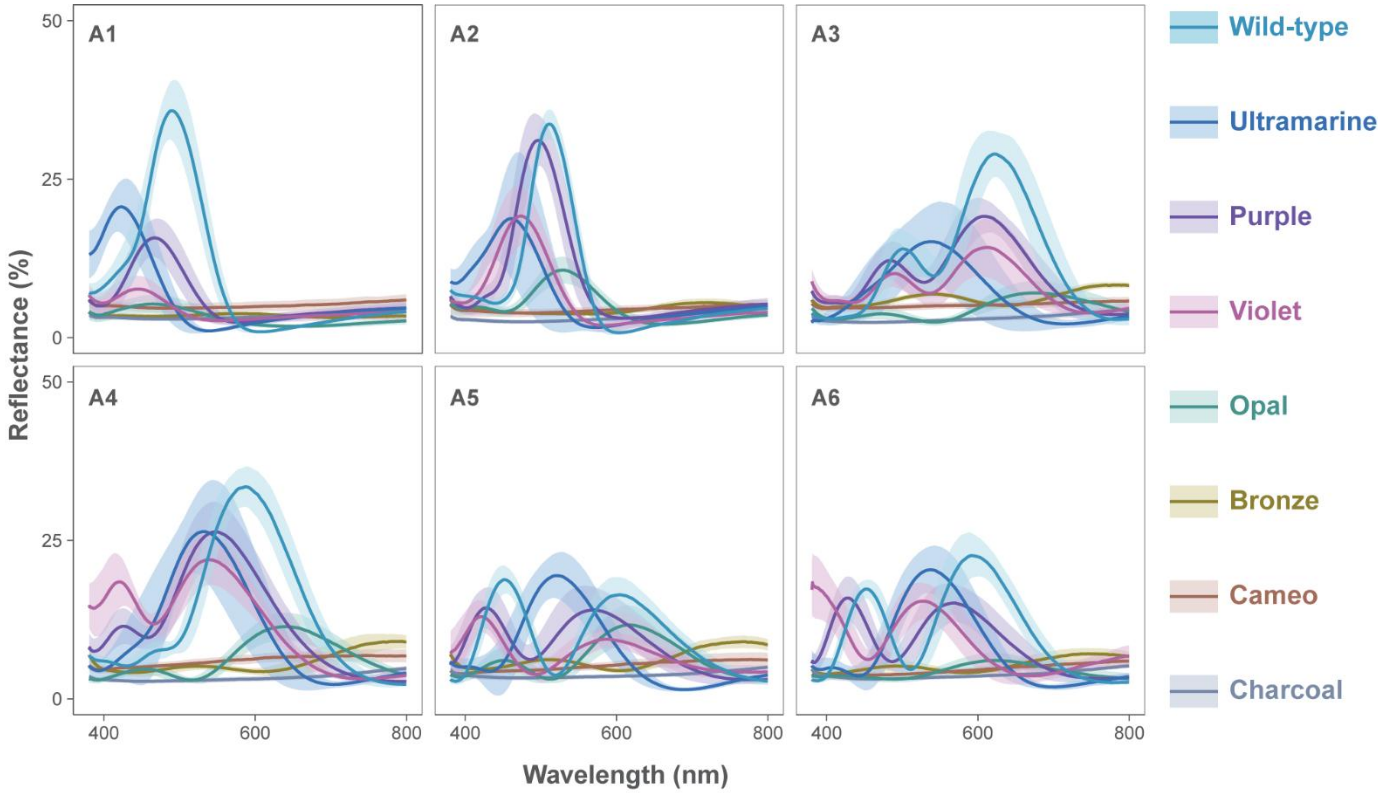


**Figure S7. Microscopic reflectance spectra at 90º angle across all eyespot areas (A1–A6) for all mutants.** For each mutant in each area, the solid line indicates the average measured reflectance across wavelengths, and the shaded area indicates the corresponding standard deviation.

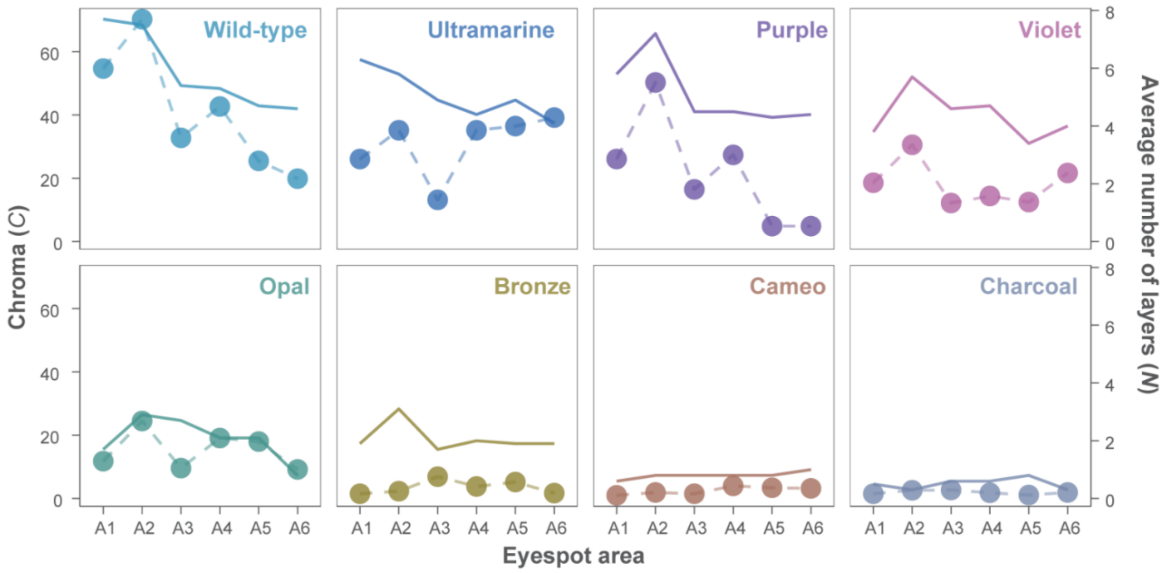


**Figure S8. Chroma intensity (*C*) across mutants and eyespot areas in relation to the average number of melanosome layers (*N*).** Chroma values (measured in Munsell chroma units) are shown as circles, while the average number of layers per area is represented by solid lines (scale on the right axis). This figure shows the high correlation of chroma intensity with the number of layers (83.9%, $t = 5.7$, $df = 94$, $P = 1.3 \times 10^{-7}$).

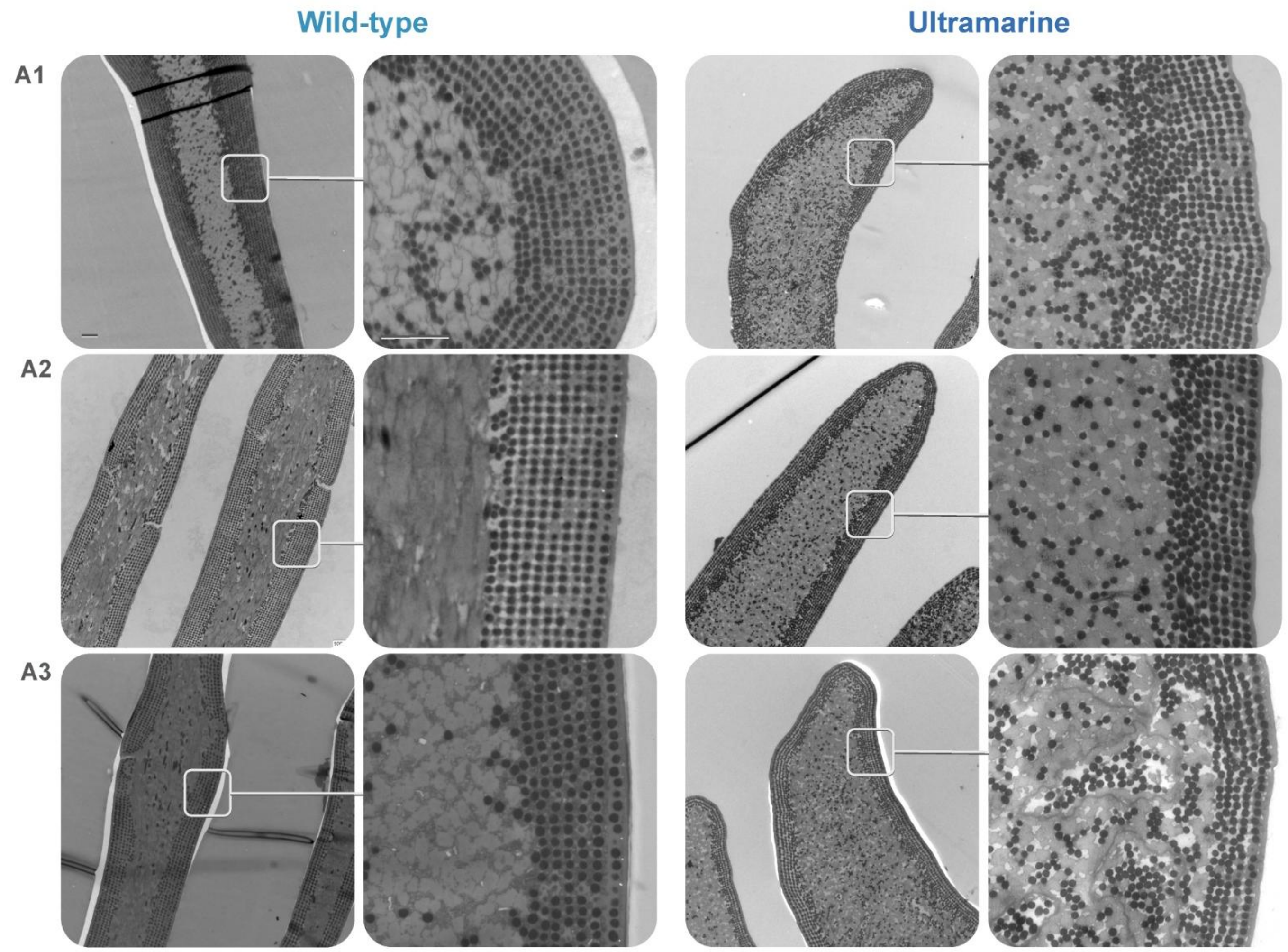


**Figure S9. Barbule ultrastructure comparison of wild-type and *ultramarine* using transmission electron microscopy** (**TEM).** Images in two scales are presented for each phenotype across the most distinct eyespot areas (A1–A3): a zoomed-out on the left (scale bar 5 µm) and a zoomed-in on the right (scale bar 1 µm).

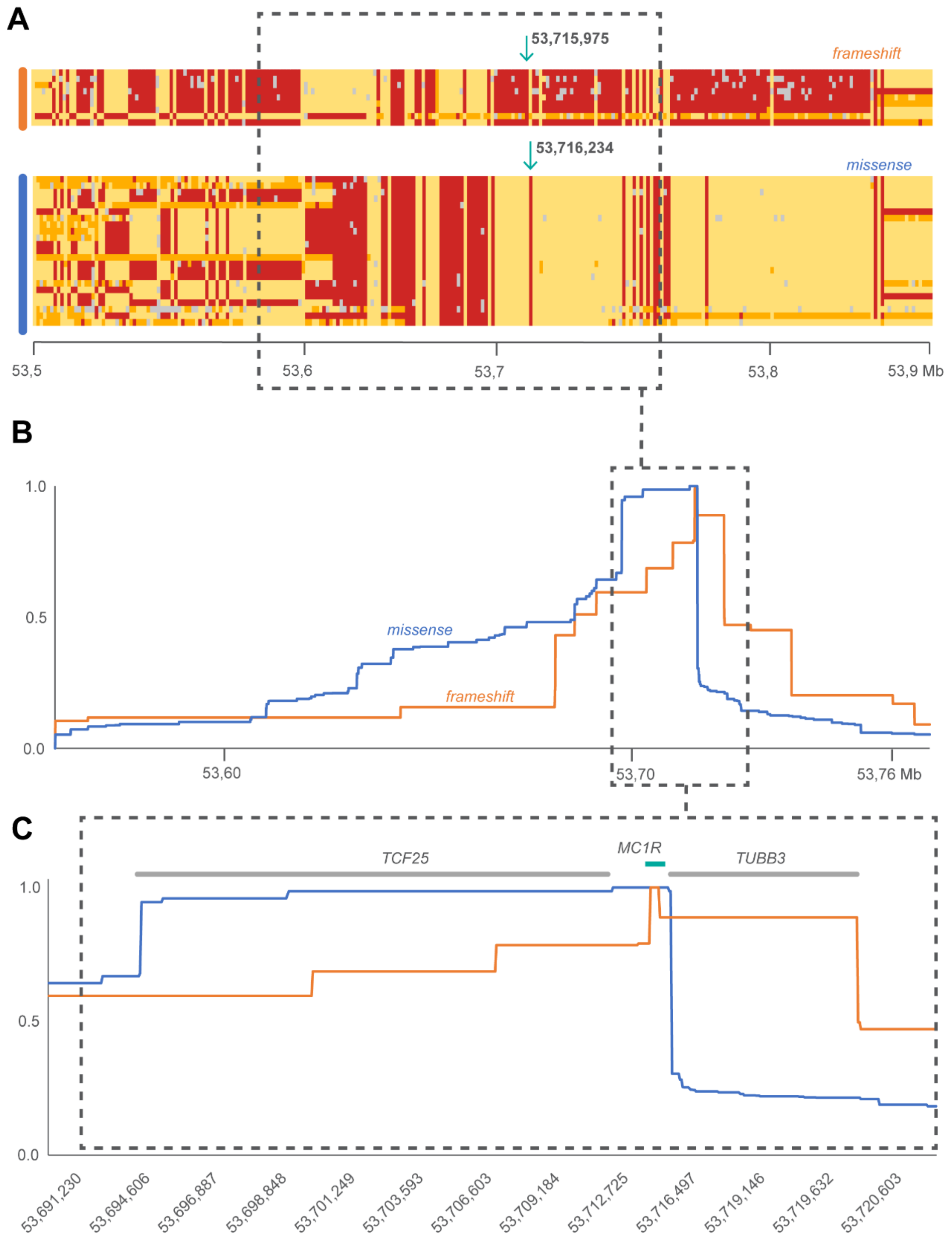


**Figure S10. Identical-by-descent (IBD) analysis of the Black-shoulder mutations. (A)** Black-shoulder haplotype structure over ~500 kb around the topmost significant variants of the genome-wide association analyses, with the relative gene position and arrows indicating the position of the two mutations [frameshift (orange): chr6:53,715,975 CA>C; missense (blue): chr6:53,716,234 T>C] (**table S2**). Each line represents a haplotype and each column a variable position with a thinning of 1,000 bp between positions. Positions containing the reference allele are colored in yellow, those containing the alternative allele are colored in red, heterozygous positions are colored in orange, and missing genotypes are colored in gray. **(B)** Extended haplotype homozygosity (EHH) within a genomic interval centered on the most significantly associated variants with the Black-shoulder color phenotype. **(C)** Zoom into the ~40kb region highlighted in **B** with a dashed black box, indicating the relative position of the *MC1R* gene (green) and the two neighboring genes (gray) in the interval. The IBD region for both protein-coding mutations is small and overlaps the *MC1R* gene.

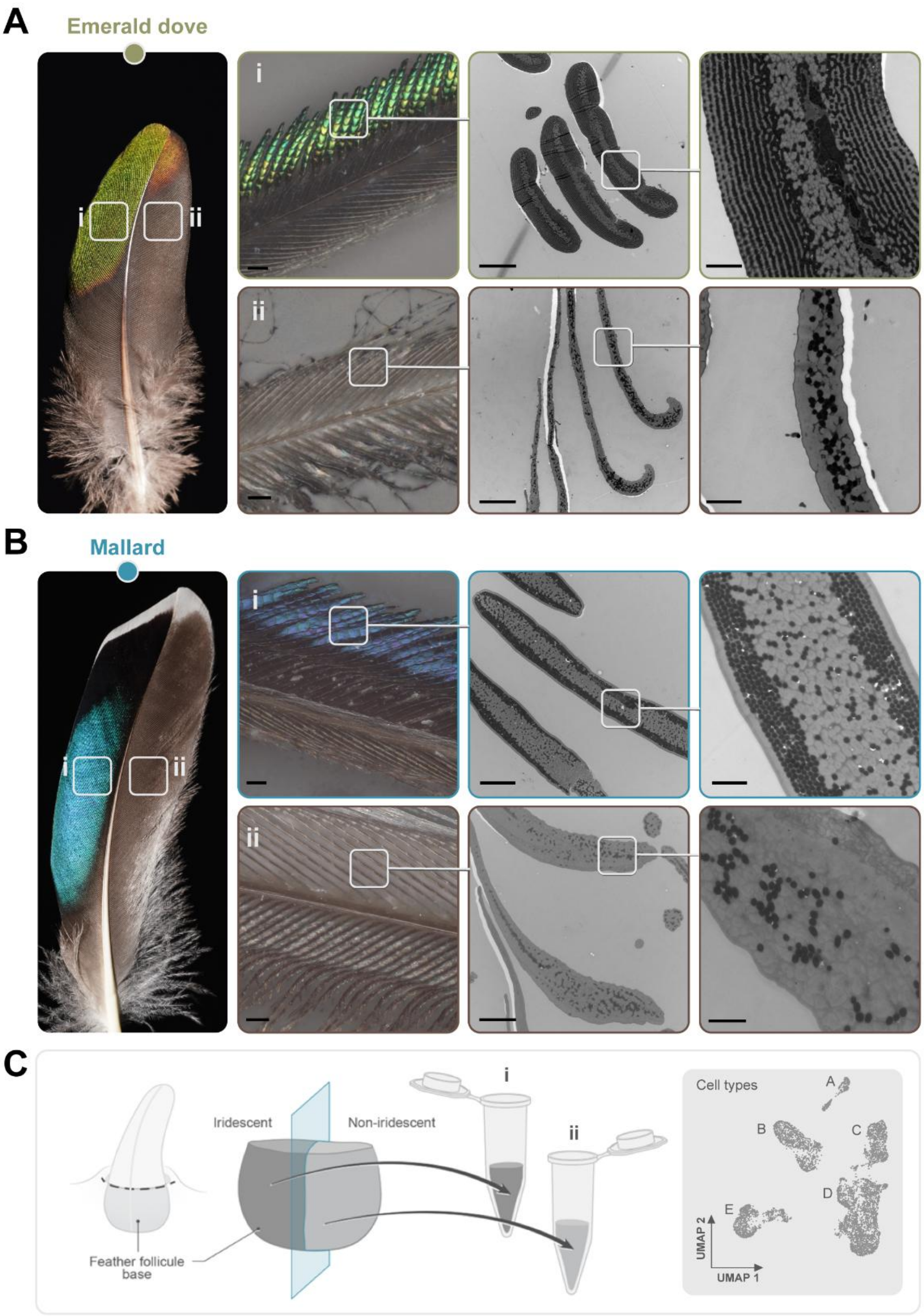


**Figure S11. Transmission electron microscopy (TEM) of iridescent and non-iridescent barbules in asymmetric feathers.** In both the emerald dove (*Chalcophaps indica*) (**A**) and the mallard duck (*Anas platyrhynchos*) (**B**), some feathers show marked coloration asymmetry, with iridescent barbules (i) developing on one side and non-iridescent barbules (ii) on the other. Left to right: optical microscopy images of individual barbs (scale bars: 50 µm) and TEM images of barbules (scale bars: 5 µm and 1 µm). Cross-sections at multiple magnifications reveal distinct barbule morphologies and melanosome organization between color types. **(C)** Schematic of the sampling strategy for the single-cell RNA-seq analyses: individual asymmetric feathers were plucked during growth at the time of iridescent color formation, the basal portion of the follicle was isolated, and each half of the feather producing iridescent barbules (i) or non-iridescent barbules (ii) was processed separately for cell dissociation and single-cell RNA-seq analyses (see Methods). The same methodology was applied for both species on two individual feathers ($n$ = 4 libraries per species).

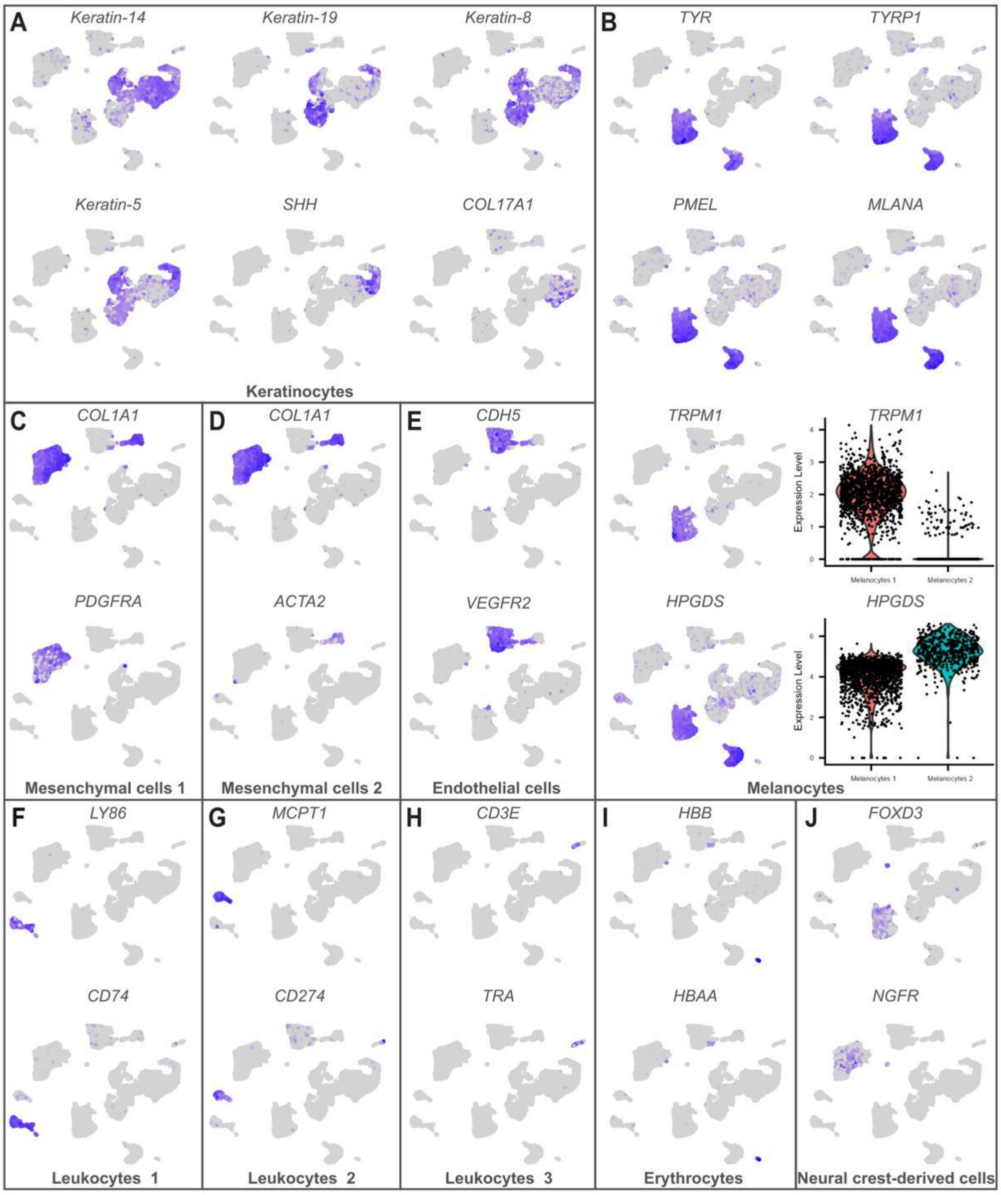


**Figure S12**. **Expression heat maps of selected genes supporting the single-cell RNA sequencing (scRNA-seq) cluster annotation for the common emerald dove**. UMAP projections based on 10,213 cells (i.e., four scRNA-seq libraries merged). **(A)** Keratinocytes: *Keratin-14* and *Keratin*-19 (type I), *Keratin-8* and *Keratin*-5 (type II), *SHH* (marker for marginal plate cells), *COL17A1* (marker for basal keratinocytes); **(B)** Melanocytes: *TYR* (required for the conversion of tyrosine to melanin), *TYRP1* (involved in eumelanin synthesis and pigment stabilization), *PMEL* (essential for melanosome formation and melanin deposition), *MLANA* (involved in melanosome biogenesis), *TRPM1* (involved in melanocyte signaling; marker for Melanocytes 1), *HPGDS* (glutathione S-transferase; enriched in Melanocytes 2; see Methods). The violin plots reports log-normalized expression for Melanocytes 1 and 2; **(C)** Mesenchymal cells 1: *COL1A1* (fibroblast marker), *PDGFRA* (fibroblast surface receptor for platelet-derived growth factors); **(D)** Mesenchymal cells 2: *COL1A1* (fibroblast marker), *ACTA2* (myofibroblast marker); **(E)** Endothelial cells: *CDH5* (plays a role in endothelial adherens junctions), *VEGFR2* (receptors for endothelial cell growth factor); **(F)** Leukocytes 1: *LY86* (lymphocyte antigen), *CD74* (regulates antigen presentation for immune response); **(G)** Leukocytes 2: *MCPT1* (mast cell–specific protease), *CD274* (ligand inhibiting T-cell activation, expressed in T cells and B cells); **(H)** Leukocytes 2: *CD3E* (part of the TCR-CD3 complex present on T-lymphocytes), *TRA* (involved in T cell activation); **(I)** Erythrocytes: *HBB* (hemoglobin subunit beta), *HBAA* (hemoglobin subunit alpha 1); **(J)** Neural crest-derived cells: *FOX3D* (promotes the development of neural crest cells from neural tube progenitors), *NGFR* (regulates neural crest-derived cell proliferation, survival, and differentiation). Log-normalized expression scale ranges from gray (no expression) to dark blue (maximum expression).

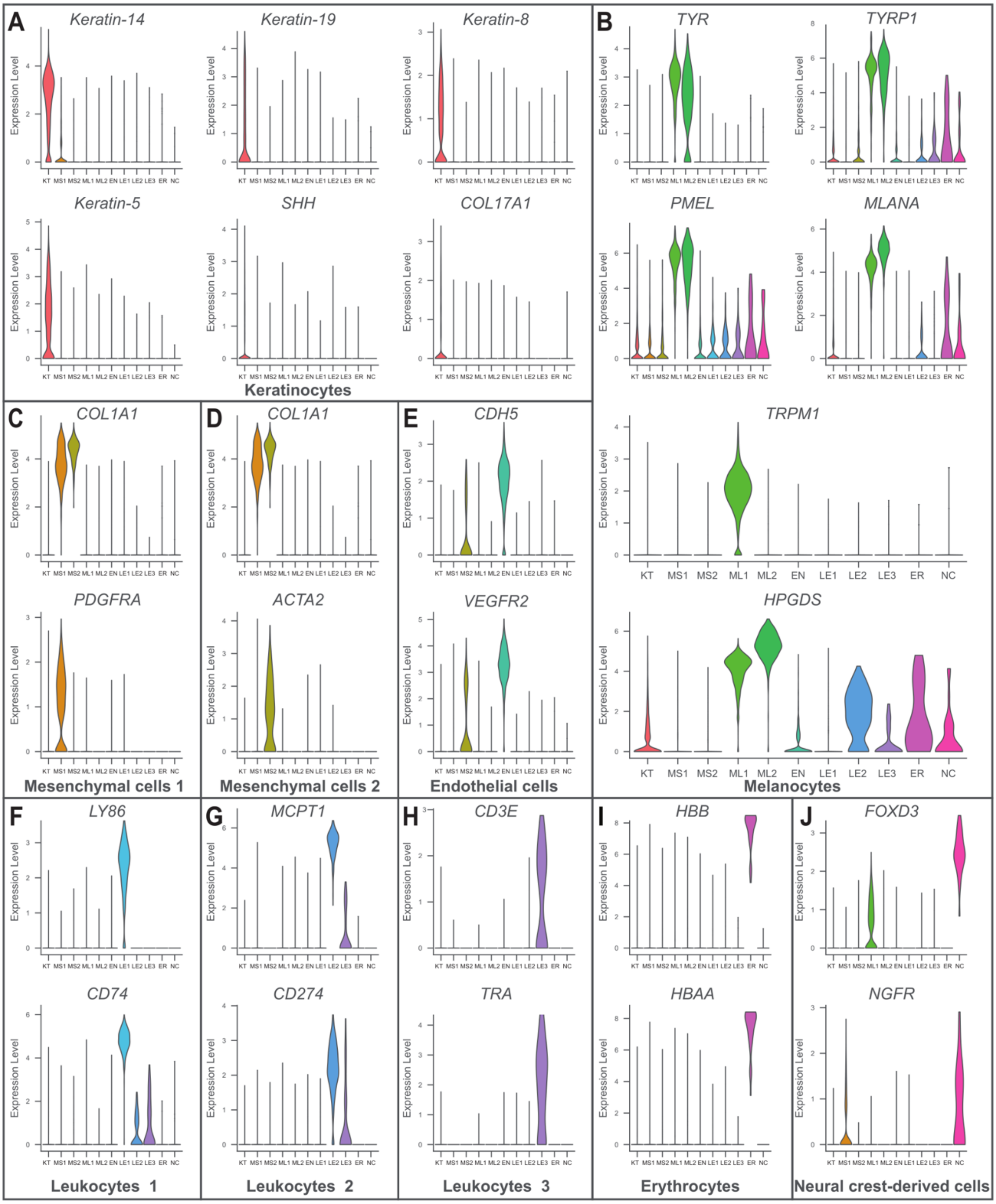


**Figure S13**. **Expression violin plots of selected genes supporting the single-cell RNA sequencing (scRNA-seq) cluster annotation for the common emerald dove**. Plots based on 10,213 cells (i.e., four scRNA-seq libraries merged). **(A-J)** The marker genes are the same as in **fig. S12**. KT: Keratinocytes; MS1: Mesenchymal cells 1; MS2: Mesenchymal cells 2; ML1: Melanocytes 1; ML2: Melanocytes 2; EN: Endothelial cells; LE1: Leukocytes 1; LE2: Leukocytes 2; LE3: Leukocytes; ER: Erythrocytes; NC: Neural crest-derived cells. The distribution of the expression levels for each gene (log-normalized) across clusters is reported.

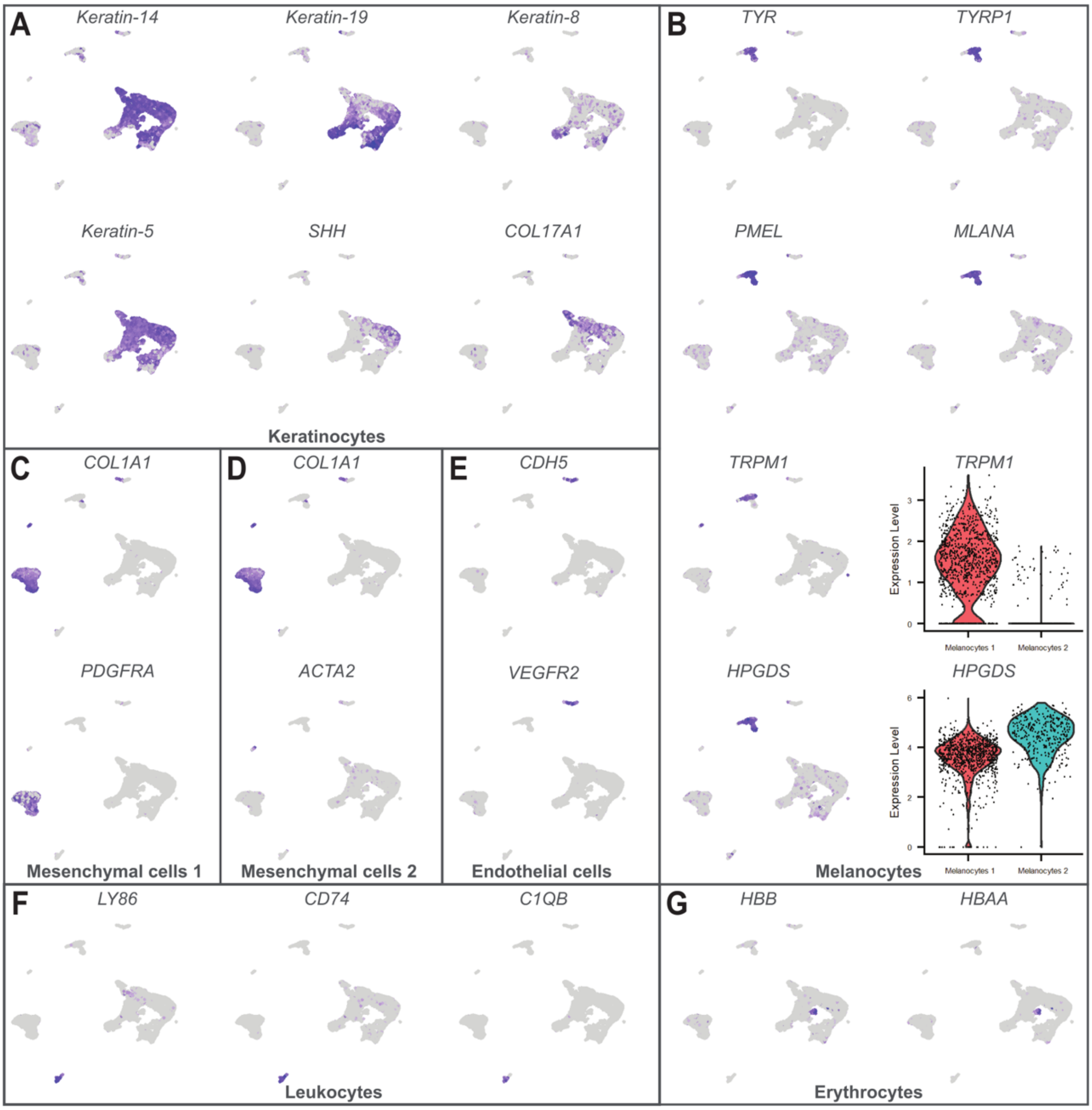


**Figure S14**. **Expression heat maps of selected genes supporting the single-cell RNA sequencing (scRNA-seq) cluster annotation for the mallard duck**. UMAP projections based on 20,991 cells (i.e., four scRNA-seq libraries merged). **(A)** Keratinocytes: *Keratin-14* and *Keratin*-19 (type I), *Keratin-8* and *Keratin*-5 (type II), *SHH* (marker for marginal plate cells), *COL17A1* (marker for basal keratinocytes); **(B)** Melanocytes: *TYR* (required for the conversion of tyrosine to melanin), *TYRP1* (involved in eumelanin synthesis and pigment stabilization), *PMEL* (essential for melanosome formation and melanin deposition), *MLANA* (involved in melanosome biogenesis), *TRPM1* (involved in melanocyte signaling; marker for Melanocytes 1), *HPGDS* (glutathione S-transferase; enriched in Melanocytes 2; see Methods). The violin plots reports log-normalized expression for Melanocytes 1 and 2; **(C)** Mesenchymal cells 1: *COL1A1* (fibroblast marker), *PDGFRA* (fibroblast surface receptor for platelet-derived growth factors); **(D)** Mesenchymal cells 2: *COL1A1* (fibroblast marker), *ACTA2* (myofibroblast marker); **(E)** Endothelial cells: *CDH5* (plays a role in endothelial adherens junctions), *VEGFR2* (receptors for endothelial cell growth factor); **(F)** Leukocytes 1: *LY86* (lymphocyte antigen), *CD74* (regulates antigen presentation for immune response), *C1QB* (component of the serum complement system); **(G)** Erythrocytes: *HBB* (hemoglobin subunit beta), *HBAA* (hemoglobin subunit alpha 1). Log-normalized expression scale ranges from gray (no expression) to dark blue (maximum expression).

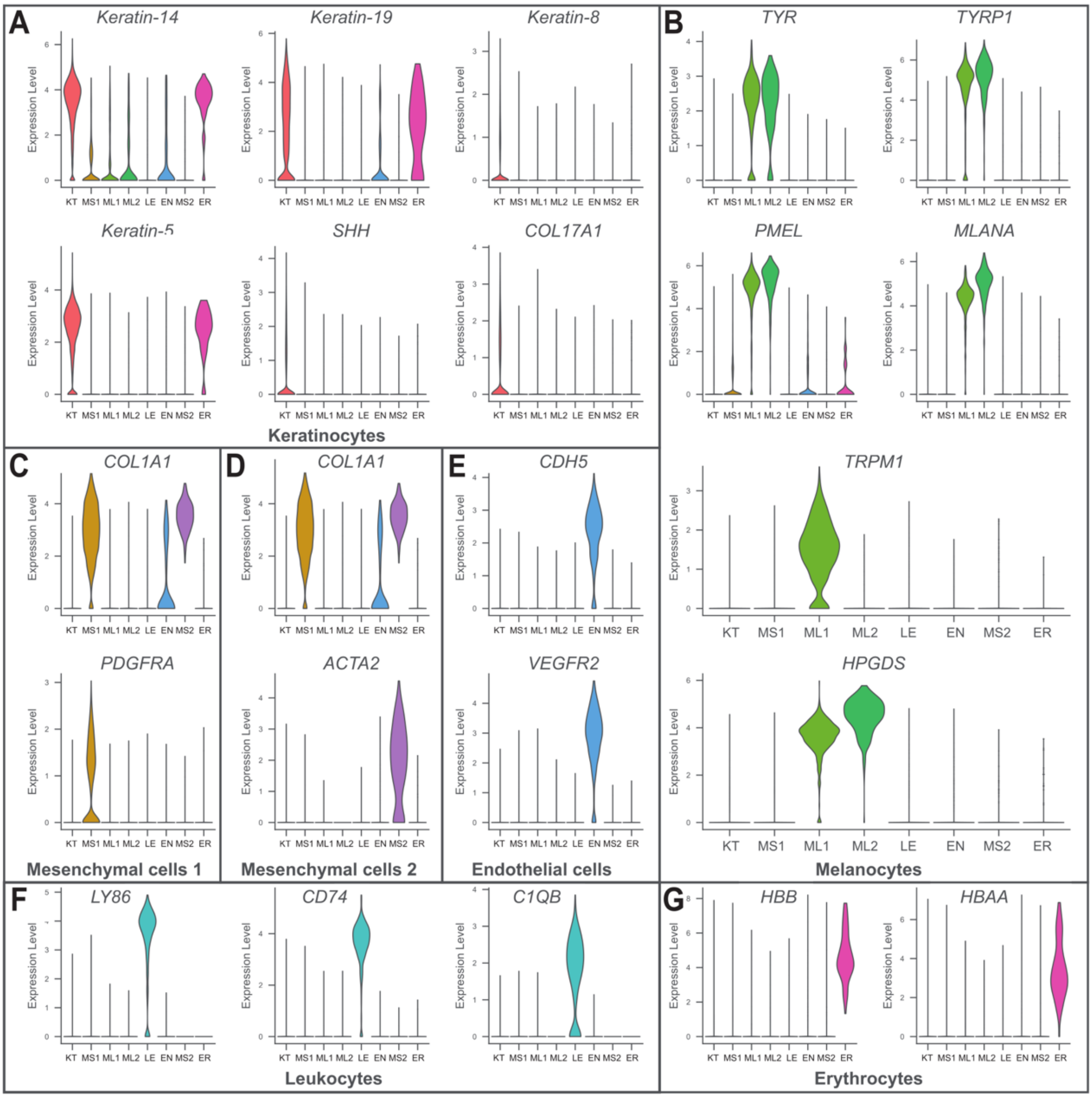


**Figure S15**. **Expression violin plots of selected genes supporting the single-cell RNA sequencing (scRNA-seq) cluster annotation for the mallard duck**. Plots based on 20,991 cells (i.e., four scRNA-seq libraries merged). **(A-G)** The marker genes are the same as in **fig. S14**. KT: Keratinocytes; MS1: Mesenchymal cells 1; MS2: Mesenchymal cells 2; ML1: Melanocytes 1; ML2: Melanocytes 2; EN: Endothelial cells; LE: Leukocytes; ER: Erythrocytes. The distribution of the expression levels for each gene (log-normalized) across clusters is reported.

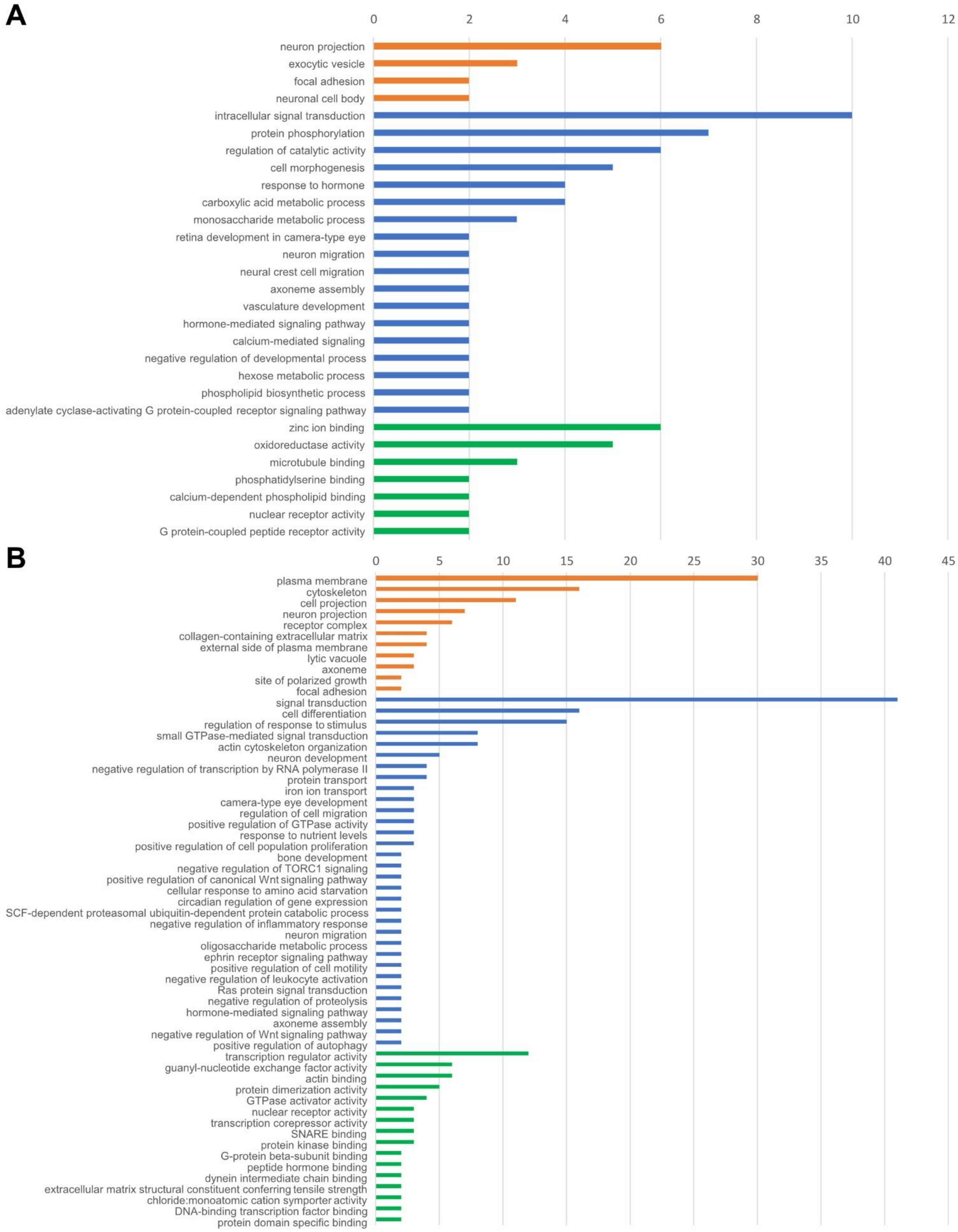


**Figure S16. Gene ontology (GO) enrichment analyses of melanocyte differentially expressed genes (DEGs) between iridescent and non-iridescent barbules (considering the merged populations of melanocytes 1 and 2). (A)** Common emerald dove. **(B)** Mallard duck. Enriched GO terms with at least two represented genes and with *P*-value ≤ 0.1 (Fisher's exact test) are reported (see **Data S5 and S6** for further details). The bars represent the number of genes within each GO term: cellular component (orange), biological process (blue), and molecular function (green).

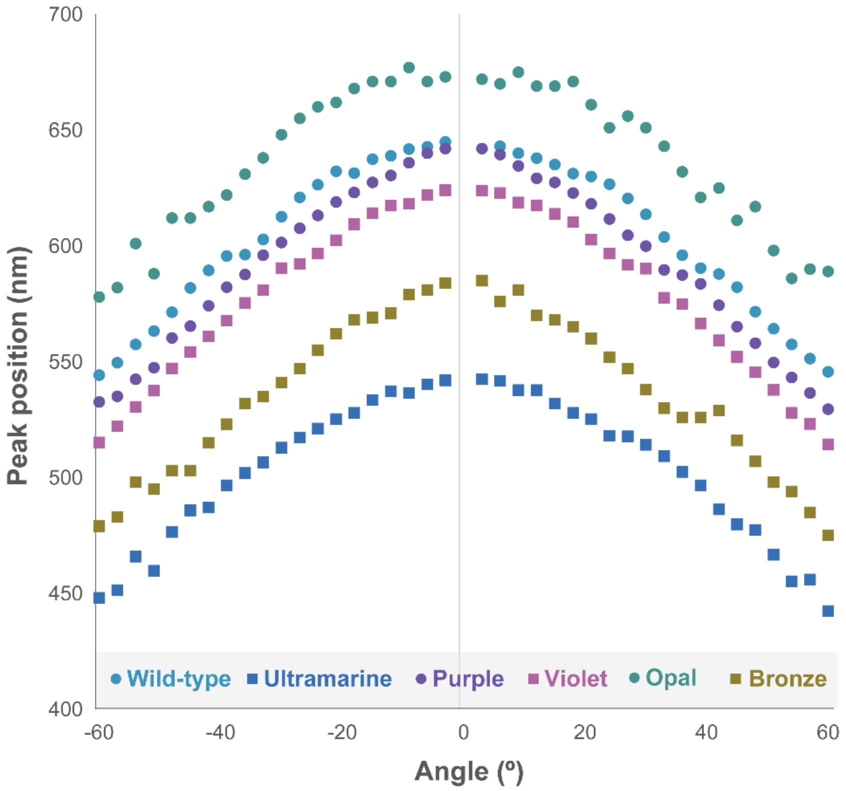


**Figure S17. Angle-dependent normalization comparison.** Measurement of the angle-dependent reflectance peak wavelength for area 3 (A3) of the eyespot for mutants exhibiting iridescence across incidence angles from –60º to 60º in 3º increments (excluding 0º). Cameo and Charcoal were excluded due to the absence of reflectance. Peak height was estimated from raw reflectance measures normalized with 200-point smoothing. This figure shows that peak positions are symmetrical between positive and negative angle measurements, allowing inferences using only one group of values.

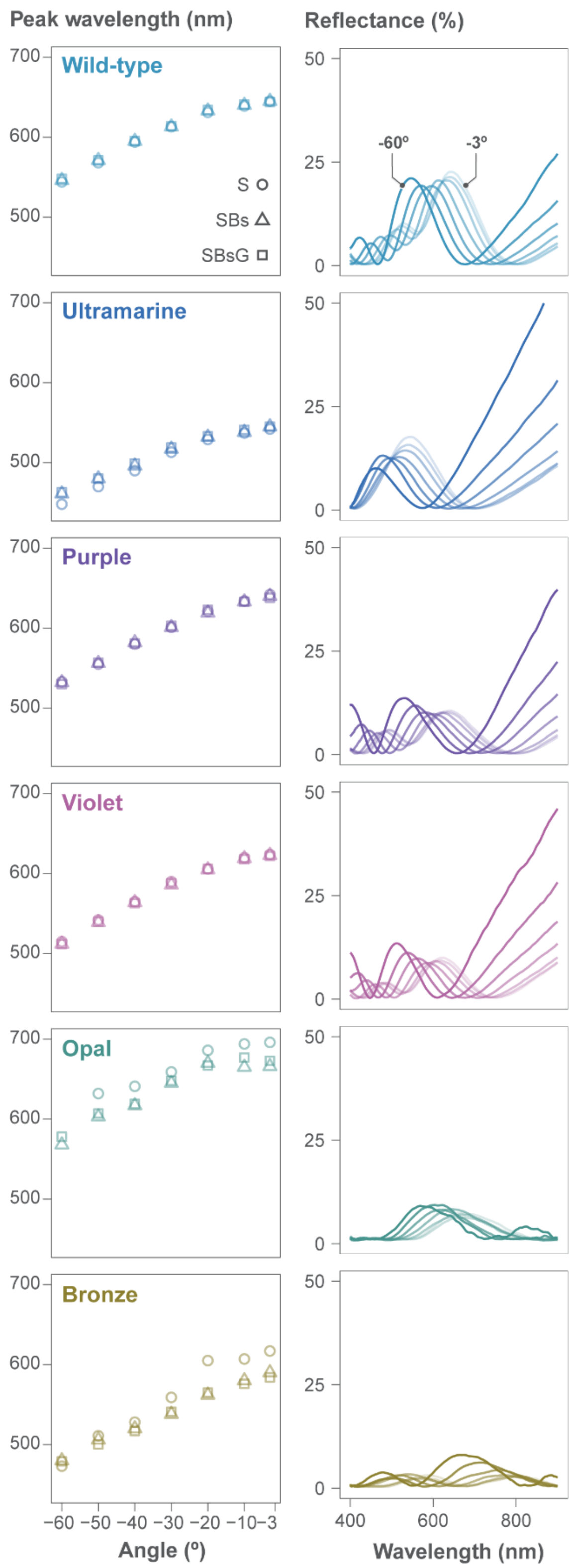


**Figure S18. Normalized angle-dependent reflectance measurements for area 3 (A3) of the eyespot.** Panels on the left represent peak wavelength for mutants exhibiting iridescence across incidence angles from -60º to -3º for three normalization strategies: 200-point smoothing (S, circle); 200-point smoothing and baseline (spline) subtraction (SBs, triangle); and 200-point smoothing and baseline subtraction with Gaussian fit (SBsG, square). Cameo and Charcoal were excluded due to the absence of reflectance. Panels on the right show the measurement of the angle-dependent reflectance considering

SBsG normalization with increasing transparency from -60 to -3º. This figure shows that data transformations do not substantially affect the peak wavelength identified across multiangle measurements.
